# Sialic acid-Dependent Binding and Viral Entry of SARS-CoV-2

**DOI:** 10.1101/2021.03.08.434228

**Authors:** Linh Nguyen, Kelli A. McCord, Duong T. Bui, Kim M. Bouwman, Elena N. Kitova, Dhanraj Kumawat, Gour C. Daskhan, Ilhan Tomris, Ling Han, Pradeep Chopra, Tzu-Jing Yang, Steven D. Willows, Andrew L. Mason, Todd L. Lowary, Lori J. West, Shang-Te Danny Hsu, S. Mark Tompkins, Geert-Jan Boons, Robert P. de Vries, Matthew S. Macauley, John S. Klassen

## Abstract

Emerging evidence suggests that host glycans influence infection by SARS-CoV-2. Here, we reveal that the receptor-binding domain (RBD) of the spike (S)-protein on SARS-CoV-2 recognizes oligosaccharides containing sialic acid (SA), with preference for the oligosaccharide of monosialylated gangliosides. Gangliosides embedded within an artificial membrane also bind the RBD. The monomeric affinities (*K*_d_ = 100-200 μM) of gangliosides for the RBD are similar to heparan sulfate, another negatively charged glycan ligand of the RBD proposed as a viral coreceptor. RBD binding and infection of SARS-CoV-2 pseudotyped lentivirus to ACE2-expressing cells is decreased upon depleting cell surface SA level using three approaches: sialyltransferase inhibition, genetic knock-out of SA biosynthesis, or neuraminidase treatment. These effects on RBD binding and pseudotyped viral entry are recapitulated with pharmacological or genetic disruption of glycolipid biosynthesis. Together, these results suggest that sialylated glycans, specifically glycolipids, facilitate viral entry of SARS-CoV-2.

---

Many viruses exploit carbohydrates attached to protein and lipid carriers, generally described as glycans, appended to host epithelial cells for viral entry^1^. Sialoglycans, acidic, sialic acid (SA)-containing glycans (e.g., gangliosides, mucin-type *O*-glycan, and complex *N*-glycans) densely displayed on the surface of mammalian cells^2^, act as co-receptors for a wide variety of viruses, including: orthomyxoviruses, paramyxoviruses, picornaviruses, reoviruses, polyomaviruses, adenoviruses, calicivirus, and parvoviruses^3^. SARS-CoV-2, which is responsible for the global outbreak of Coronavirus Disease 2019 (COVID-19) and a member of the coronavirus family, is believed to rely on a combination of angiotensin-converting enzyme 2 (ACE2) and glycans to bind and infect the lungs, as well as other tissues and organs^4–6^. Human coronaviruses generally rely on glycans to assist in cell entry^7^. For example, Middle East Respiratory Syndrome Coronavirus (MERS-CoV) binds sialoglycans to facilitate cellular entry^8^, the human betacoronaviruses OC43 and HKU1 engage sialoglycans with 9-*O*-acetylated SA as key receptors^9^, while Severe Acute Respiratory Syndrome Coronavirus (SARS-CoV-1) and CoV-NL63 exploit acidic heparan sulfate (HS) polysaccharides^10,11^.

There is emerging evidence that acidic glycans serve as co-receptors for SARS-CoV-2. Electrospray ionization mass spectrometry (ESI-MS) analysis revealed that binding of oligosaccharide fragments of heparin, a highly sulfated form of heparin sulfate (HS), to the receptor binding domain (RBD) of the transmembrane spike (S) glycoproteins of SARS-CoV-2 inhibits ACE2 binding^12^. Consistent with this finding, Kwon *et al.* showed that free HS inhibits SARS-CoV-2 infection of Vero cells^13^. Esko and coworkers reported that HS enhances the affinity of the SARS-CoV-2 RBD for ACE2, suggestive of HS acting as a more classical co-receptor^14^. Notably, destroying cellular HS enzymatically with heparanase, or competing with unfractionated heparin, both significantly reduced SARS-CoV-2 infection^14^. It was speculated early on that SARS-CoV-2 may exploit sialoglycans on cells^15^, and some evidence has emerged suggesting that SA can bind SARS-CoV-2. Results of biolayer interferometry showed that SA-conjugated gold nanoparticles exhibit high avidity for the SARS-CoV-2 S1 protein, which contains both the *N*-terminal domain (NTD) and RBD of the S protein^16^. Efforts to quantify these interactions on model sialylated species (e.g. 3’-sialyllactose), however, were challenging and only revealed weak binding (*K*_d_ ~mM)^16^. Given this weak binding, it is, perhaps, not surprising that traditional glycan microarray screening found no significant interactions between the RBD and sialylated *N*-glycans or gangliosides^4^. Beyond acidic glycans, recent work also suggests that the RBD mimics a galectin scaffold and can bind blood group A antigen^17^. Moreover, it was also recently reported that the sialylated *N*-and *O*-glycans on ACE2 may play a role in S-protein cell binding^18,19^.

In light of the suggested role for glycans in SARS-CoV-2 infection, we analyzed a large library of glycans as ligands for the RBD of SARS-CoV-2. Using catch-and-release electrospray ionization mass spectrometry (CaR-ESI-MS)^20–22^, a label-free method for quantifying weak, yet biologically-relevant, interactions within a complex mixture, we discovered that several classes of sialoglycans are bound by the RBD, with ganglioside oligosaccharides being the top hits. Cell-based studies reveal that RBD binding and SARS-CoV-2 pseudotyped virus entry of ACE2-expressing cells is decreased upon decreasing SA levels on cells pharmacologically, genetically, or enzymatically. Blocking glycolipid biosynthesis produced similar decreases in RBD binding and viral infection, pointing to RBD-glycolipid interactions as being important for SARS-CoV-2 infection of cells.

## Results

### CaR-ESI-MS screening of glycan libraries

The SARS-CoV-2 S protein possesses 26 potential glycosylation sites (22 *N*- and 4 *O*-glycosylation sites)^23^. This extensive degree of glycosylation, together with the appearance of multiple oligomeric states (monomer, dimer, trimer and hexamer) in ESI-MS analysis (Supplementary Fig. 1a), makes the application of CaR-ESI-MS screening to the S protein challenging. Consequently, the RBD, which contains only two *N*- (N331 and N343) and two *O*-glycosylation sites (T323 and S325), was selected for CaR-ESI-MS screening (Supplementary Fig. 1b,c)^24–27^. A total 140 glycans (Supplementary Fig. 2), which consist mostly of mammalian glycans (**1–133**) but also several non-mammalian rhamnose-containing oligosaccharides as negative controls (**134**–**140**), were used for screening against the SARS-CoV-2 RBD. To ensure that all species were of different molecular weights, the library was divided into sixteen sub-libraries of glycans (Library A – P, Supplementary Table 1 and Supplementary Fig. 3). In all cases, a reference protein (P_ref_) was added to control for non-specific glycan-RBD interactions during ESI^28^.

Numerous glycan ligands were released from RBD in CaR-ESI-MS performed at both at 25 °C (Supplementary Fig. 4 and 37 °C (Fig. 1a). The low relative abundances of released ligands are indicative of generally low affinities. No binding was detected for the non-mammalian rhamnose-containing glycans (**134**–**140**) or any detectable glycans released from P_ref,_. These results indicate that the low abundance ligands released from the RBD are the result of specific interactions and not false positives.

**Figure 1.**
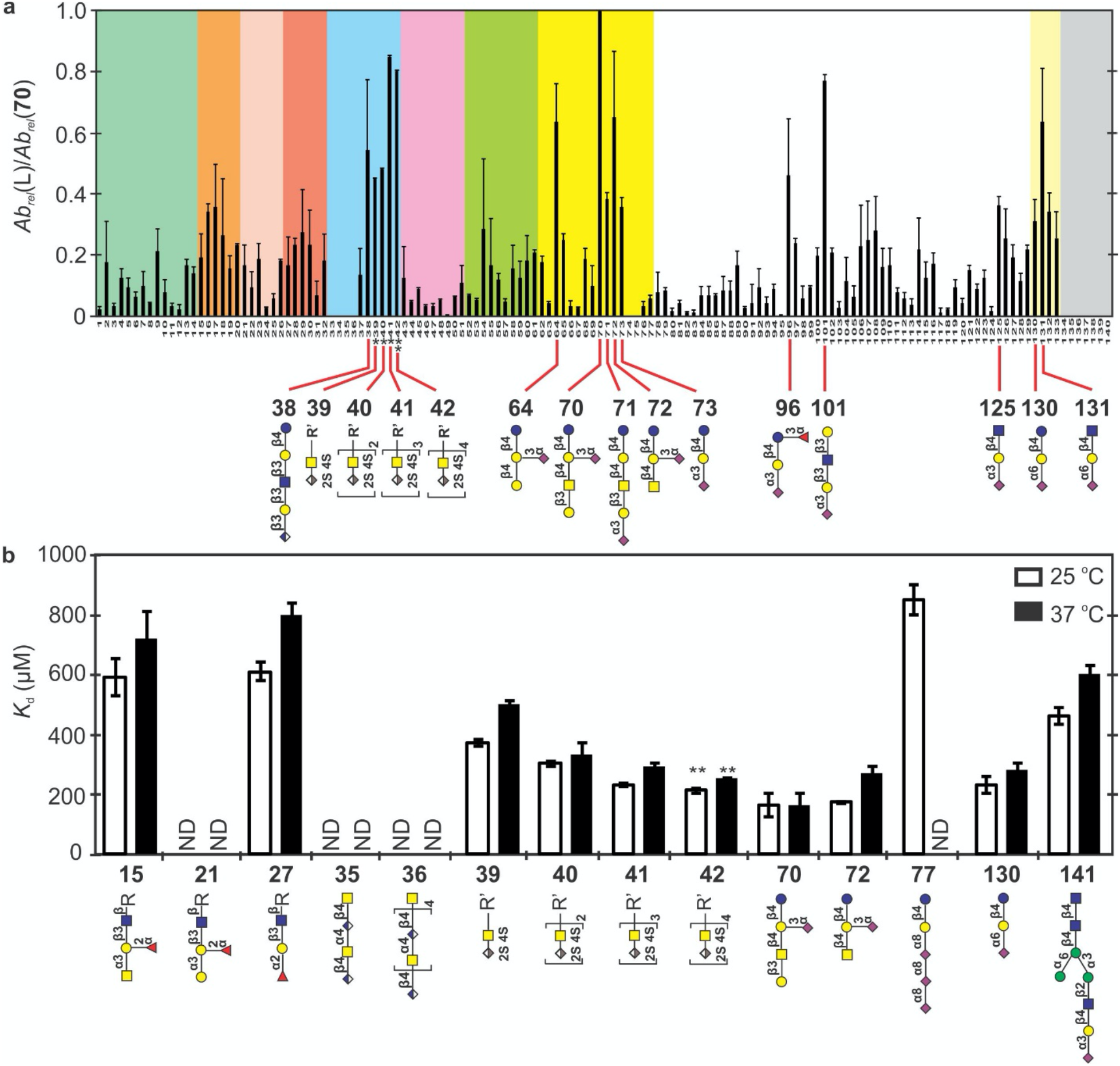
Glycan library screening and affinities for RBD. **a**, Normalized abundances of released glycans from the SARS-CoV-2 RBD by CaR-ESI-MS at 37°C. Summary of the charge-normalized (relative to 70) abundances of released ligands measured by CaR-ESI-MS screening of *Library* A – O against SARS-CoV-2 RBD. Measurements were performed in negative ion mode with a UHMR Orbitrap mass spectrometer at an HCD energy of 50 V on aqueous ammonium acetate (100 mM, pH 6.9) solutions of SARS-CoV-2 RBD (13 μM) and glycan library (containing 50 nM of each glycan). The different classes of oligosaccharides are distinguished by colour: mint green - Lewis antigens (**1**–**14**); light orange - blood group A antigens (**15**–**20**); light pink - blood group B antigens (**21**–**26**); tropical pink - blood group H antigens (**27**–**32**); ice blue - sulfated compounds (**34**–**38**) and heparan sulfates (**39**–**42**), light violet - antigen - related glycans (**43**–**51**), chartreuse - globo (**52**–**61**), yellow - ganglioside oligosaccharides (**62**–**77**), white - HMOs and other glycans (**78**–**129**), light yellow - Neu5Acα2-6-linked oligosaccharides (**130**–**133**), and grey rhamnose - containing compounds (**134**–**140**). * relative abundances (in CaR-ESI-MS) estimated from their relative (to **70**) *K*_d_ values. ** **42** sample also contained some **41** and the affinity reflects the estimated relative concentrations of **41** and **42**. **b,** Affinities of glycans (**15**, **21**, **27**, **35**, **36**, **39 – 42**, **70**, **72**, **77**, **130** and **141**) for RBD. *K*_d_ (μM) values measured by ESI-MS in aqueous ammonium acetate (100 mM, pH 6.9) solutions at 25°C (white bars) and 37°C (black bars). ND ≡ not detected.

At 37 °C, the pentasaccharide (**70**) of the ganglioside GM1 was the preferred ligand. Other gangliosides were also recognized, with a preference for monosialylated (**70** – **73**) over di-and trisialylated gangliosides. Acidic human milk oligosaccharides (HMOs) were consistently observed at high relative abundances and some of the ABH blood group antigens (in particular A and H types 2, 3 and 4) exhibit moderate CaR-ESI-MS signal. Interestingly, chondroitin sulfate (**33**, **34**) and heparosan-derived oligosaccharides containing GlcA (**35**, **36**), which are acidic glycans, showed little to no binding to RBD. These results, when considered with the preference for monosialylated over di-and trisialylated ganglioside oligosaccharides, argue against electrostatics being the dominant force underpinning RBD–ligand binding. Overall, the screening results obtained at 25 °C are similar to those obtained at 37 °C in terms of the glycans that are recognized, with only minor changes in the relative abundances for some ligands (Supplementary Fig. 4). HS glycans (**39**–**42**) were not included in CaR-ESI-MS screening due to difficulties in releasing them from RBD at the energies used to screen the other glycans; the higher energies required for HS release promote fragmentation of the other ligands. Therefore, the relative abundances of these HS structures shown in Fig. 1a were estimated from their affinities, measured by ESI-MS, relative to that of the highest affinity ligands identified by screening. Based on this analysis, HS **41** and **42** were predicted to have CaR-ESI-MS responses slightly lower than that of **70**.

### Affinities of the RBD towards select glycans

Quantifying interactions of ligands to individual glycoforms of RBD by ESI-MS is challenging due to the unknown concentration of each glycoform and the potential for spectral overlap (ligand-bound and free RBD species). Therefore, determining the affinities of RBD for glycans required the elucidation of glycoforms and an estimation of their respective concentrations, which was based on an assumption that all glycoforms have uniform ESI-MS response factors. A total of 77 distinct RBD MWs were identified by ESI-MS (Supplementary Table 2 and Supplementary Fig. 5a). According to reported glycomics studies, RBD produced from HEK293 cells has predominantly complex type *N*-glycans and core 1 and 2 mucin-type *O*-glycans (Supplementary Table 3)^24–27^. HILIC-UHPLC-FLD/ESI-MS analysis of the *N*-glycans released from the RBD sample used in this work identified 136 distinct structures, with 74 distinct compositions (Supplementary Table 4 and Supplementary Fig. 6). Based on the reported *O*-glycans and results of the current *N*-glycan analysis, the glycan composition of each RBD species was putatively assigned (Supplementary Table 5). It is found that the major glycoforms possess 2–8 Neu5Ac residues and are di- and trifucosylated (Supplementary Fig. 5b).

The affinities of the top hits (gangliosides **70** and **72**) and moderate binders (**15**, **21**, **27**, **77**, and **130**) identified by CaR-ESI-MS screening of the defined library, as well as the HS **39** – **42** for RBD were measured by ESI-MS at pH 6.9 and 25 °C and 37 °C. Two non-binders (**35** and **36**) were also included as negative controls. The reported affinities are the average values measured for all the RBD species detected (Fig. 1b and Supplementary Table 6). It is notable that RBD glycosylation has a minimal effect on the measured *K*_d_ (Supplementary Fig. 7), suggesting that the glycosylation sites are remote from the glycan binding site(s) and do not strongly influence the conformations of glycan binding motifs within RBD.

The measured affinities are all ≥100 μM, with ganglioside **70** (GM1) having the highest affinity (160 ± 40 μM) for RBD at 25 °C. For the HS **39** – **42**, the trend in affinities is **39** ≈ **40** > **41** ≈ **42**, with *K*_d_ ranging from 200 μM to 400 μM. With the exception of **77** (which was undetectable at 37 °C), the temperature dependence of the measured *K*_d_ was relatively minor, with changes of less than 50%. Based on the difference in affinities, the average association enthalpy change is only −3.4 kcal mol^−1^, which is modest for protein–glycan interactions; however, the entropy change is also favourable (with average entropy changes of 10–20 cal K^−1^ mol^−1^). The trend in affinities agree well with the trends in relative abundances measured by CaR-ESI-MS (Supplementary Fig. 8), establishing the reliability of CaR-ESI-MS for identifying glycan ligands and differentiating the high affinity ligands from the low affinity ligands and non-binders.

### Tissue-derived ***N***-glycan libraries

Direct evidence of binding between the S-protein or RBD and host *N-*glycans is lacking. To assess the *N-*glycan binding properties of RBD, libraries were prepared from lung and intestinal tissues and screened against RBD by CaR-ESI-MS at 25 and 37 °C. At 37 °C, screening of the lung tissue-derived *N-*glycans detected ligands with 19 distinct MWs (Supplementary Table 7), consisting of 13 mono- and six disialylated *N*-glycans, of which six are hybrid type and the remainder are complex type. The absence of any trisialylated *N*-glycan ligands may reflect their low abundances and low affinities, compared to those of the mono- and disialylated *N-*glycans (Fig. 2a). Two of the hits detected by CaR-ESI-MS, with *m/z* of 1549.55 ([M-H]^−^, HexNAc_3_Hex_3_ Fuc_1_Neu5Ac_1_) and 1889.65 ([M-H]^−^, HexNAc_3_Hex_6_Neu5Ac_1_), were not observed in HILIC-UHPLC-FLD analysis, again, suggesting their low abundances, yet high affinity (Fig. 2b). Ligands with 29 distinct MWs were detected from screening of the intestinal tissue *N*-glycans (Fig. 2c and Supplementary Table 8). The hits consist of 14 mono- (with one sulfated), seven di-, and five trisialylated *N*-glycans, of which six are hybrid type and the rest are complex type. Three of these hits (HexNAc_3_Hex_3_Neu5Ac_1_, HexNAc_3_Hex_5_Neu5Ac_1_, and HexNAc_3_Hex_6_Neu5Ac_1_) were not observed in HILIC-UHPLC-FLD analysis, presumably because of their low abundances. Notably, no neutral *N*-glycan ligands were detected for either library. With the exception of the glycan of composition HexNAc_4_Hex_5_Neu5Ac_2_, which was by far the most abundant glycan in the intestinal library (Fig. 2d), no *N*-glycans were released from P_ref_ in any of the CaR-ESI-MS measurements, indicating an absence of a contribution of non-specific binding to the remaining hits.

**Figure 2.**
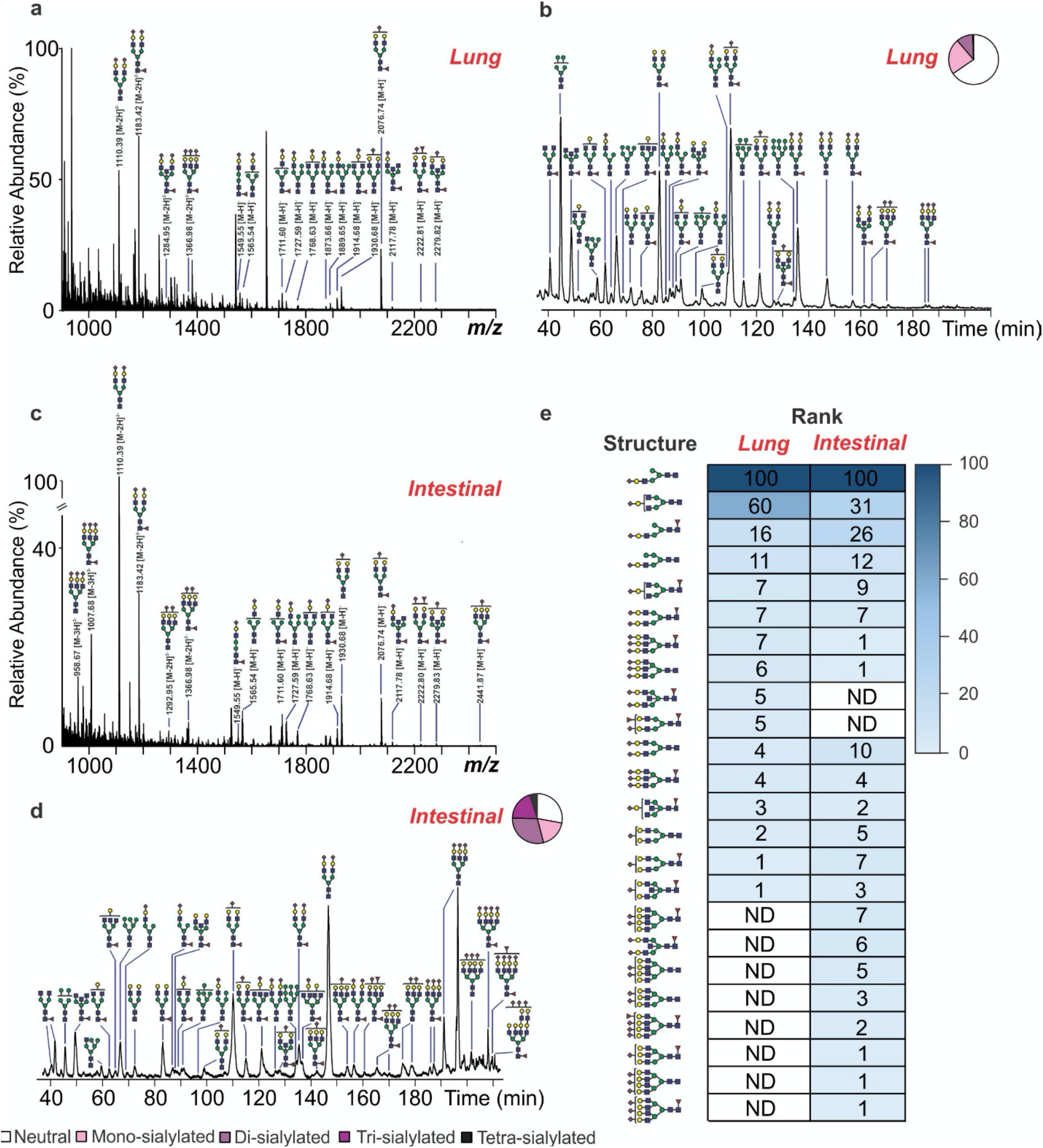
Screening of natural *N*-glycan libraries against RBD. **a,c,** CaR-ESI-MS screening results obtained for aqueous ammonium acetate solutions (100 mM, pH 6.9, 37 °C) of RBD (13 μM), P dimer of the Saga strain (P_ref_, 4 μM) and the *N*-glycan library from lung (200 μg mL^−1^) (**a**) and intestinal (125 μg mL^−1^) (**c**) tissues. Ions with *m/z* of 3000 – 4000 were subjected to HCD using a collision energy of 50 eV. **b,d,** Chromatogram of 2-AB-labeled *N*-glycans released from lung (**b**) and intestinal (**d**) tissues acquired using HILIC-UHPLC with fluorescence detection. The relative abundances of the Neu5Ac content of the *N*-glycans are indicated graphically. **e,** Heat maps of relative affinities of ligands identified by CaR-ESI-MS screening; ND ≡ not detected.

Based on the relative abundances of released *N*-glycans measured by CaR-ESI-MS and their relative concentrations in each library (deduced from UHPLC analysis), the relative affinities of the *N*-glycan ligands were ranked (Fig. 2e). From the resulting heat maps, it can be seen that RBD has a strong preference for monosialylated monoantennary *N*-glycans, with the clear top hit having the composition HexNAc_3_Hex_4_Neu5Ac_1_. Based on the calculated glucose unit values from elution times of *N*-glycans (the HPLC elution position, expressed as glucose unit value, for each glycan is related to the number and the linkage types of its constituent monosaccharides), this ligand is sialylated on the α-(1-3)-arm^29^. To relate the CaR-ESI-MS results acquired for the defined and natural *N*-glycan libraries, we also performed quantitative binding measurements on **141** (HexNAc_3_Hex_4_Neu5Ac_1_, with a Neu5Acα2-3 linkage), which has the same glycan composition as the top hit identified from screening. Affinities of 460 ± 30 μM and 600 ± 30 μM were measured at 25 °C and 37 °C, respectively (Fig 1b).

### CaR-ESI-MS screening of gangliosides

The CaR-ESI-MS screening revealed a preference for the oligosaccharide derived from gangliosides, with GM2 and GM1 being the top hits. As gangliosides are glycolipids normally embedded within a lipid bilayer, we screened the RBD against a library of six gangliosides (GM1, GM2, GM3, GD1a, GD2 and GT1b, each nominally 1% of total lipid), presented together in a nanodisc composed of 1,2-dimyristoyl-sn-glycero-3-phosphocholine (DMPC)^30^. To positively identify the gangliosides bound to RBD, ions centered at *m/z* 3,540 that fell within a 100 *m/z* window, which encompasses ions at charge state −9 of any RBD-ganglioside complexes present (Supplementary Fig. 9a), were isolated and then collisionally heated to release bound gangliosides (Supplementary Fig. 9b). Notably, signals corresponding to deprotonated GM1, GM2, and GM3 ions were measured; no signals corresponding to the disialylated gangliosides – GD1a, GD2, or GT1b – were detected. As a negative control, these measurements were repeated using identical experimental conditions but in the absence of RBD and no ganglioside ions were detected (Supplementary Fig. 9c). Even though GM3 was the most abundant of the gangliosides detected, this may not represent an intrinsic affinity for the RBD, but rather a greater release efficiency from the ND as the RBD complex, as suggested from a previous study^30^.

### Inhibition by SA suggests multiple glycan binding sites

The screening and affinity results reveal that the RBD binds preferentially to sialoglycans and HS oligomers and also binds to other structures, such as ABH antigens (Fig. 1a). The diversity of recognized structures raises the possibility that the RBD possesses multiple glycan binding sites, with distinct binding properties. Experimentally derived structural data has yet to be reported for RBD–glycan complexes and the only glycan binding site (for HS) was predicted from molecular docking^6^. With the goal of testing whether RBD possesses multiple glycan binding sites, we performed CaR-ESI-MS screening of the 16 sub-libraries of defined glycans against RBD in the presence of 20- and 50-fold excess (relative to RBD) of α-methyl glycoside of Neu5Ac (Neu5AcαMe) at 25 °C (Fig. 3). These conditions are expected to lead to a reduction in the concentration of RBD binding site(s); at least for those that recognize SA.

**Figure 3.**
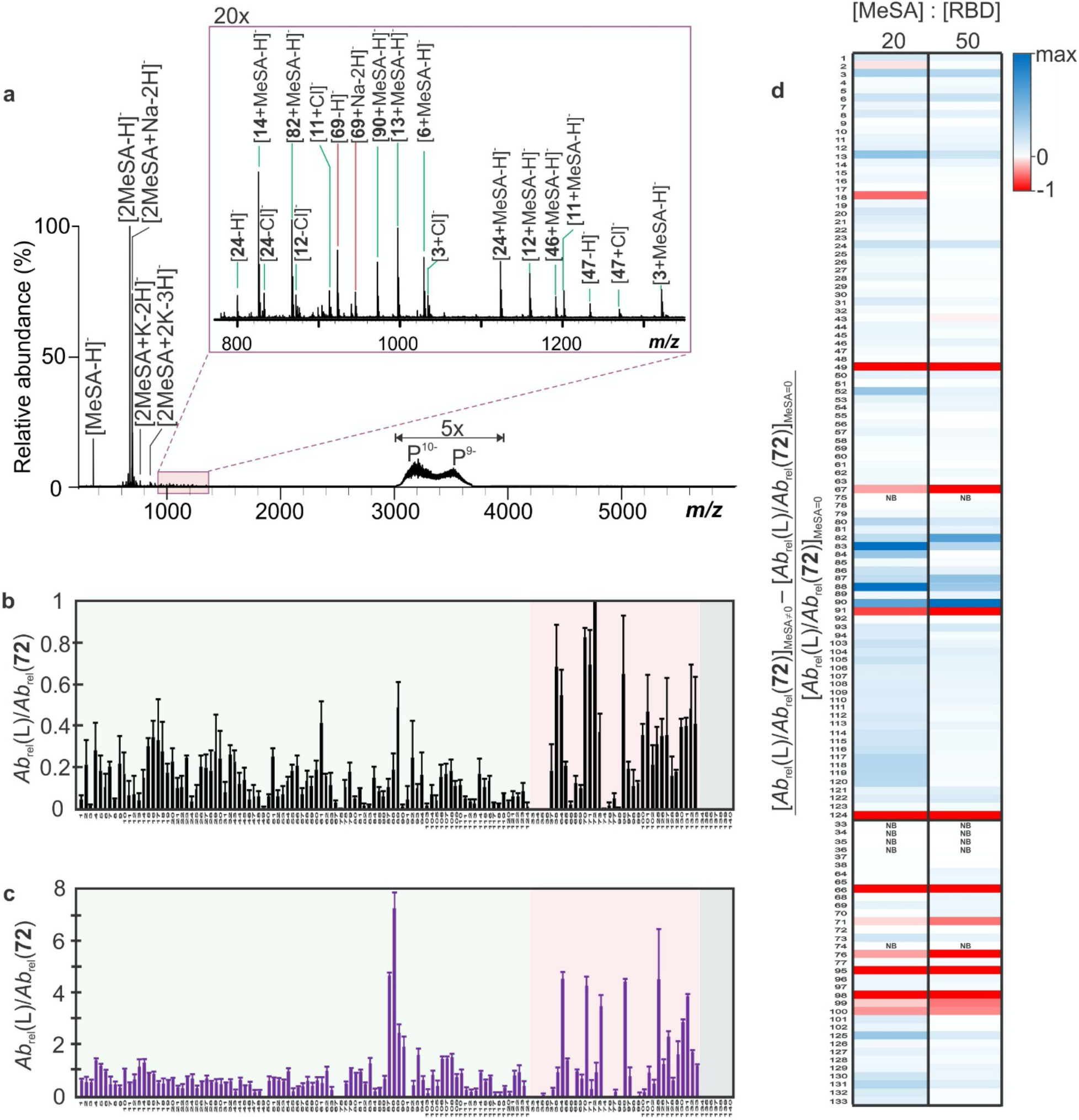
Inhibition of glycan binding to RBD by methyl-α-Neu5Ac. **a,** Representative CaR-ESI-MS screening obtained for aqueous ammonium acetate solutions (100 mM, pH 6.9, 25 °C) of RBD (10 μM), SNA (P_ref_, 4 μM), Neu5AcαMe (500 μM) and Library C (50 nM of each glycan). Ions with *m/z* 3000 – 4000 were subjected to HCD using a collision energy of 50 eV. **b,c**, Relative abundances of released glycans measured by CaR-ESI-MS for 16 sub-libraries (50 nM of each glycan) in 100 mM aqueous ammonium acetate solutions (pH 6.9, 25°C) containing RBD (10 μM) with 0 μM (**b**) and 500 μM (**c**) Neu5AcαMe (MeSA), in which light green, light pink and grey represent the neutral and acidic glycans and controls (Lib P), respectively. **d**, Heat maps of the change of normalized abundances (to **72**) of released glycans with the change of **72** after addition of SA measured by CaR-ESI-MS in the presence and absence of Neu5AcαMe (NB ≡ no binding).

The Neu5AcαMe concentration-dependent relative abundances of released ligands are summarized in Fig. 3d. Inspection of the glycan screening results reveals that, at high Neu5AcαMe concentration, the most abundant ligand released from RBD was not **72**, but rather the neutral pentasaccharide **88** (Galα(1-3)[Fucα(1-2)]Galβ(1-4)[Fucα(1-3)]Glc), which contains a fucosylated B type VI blood group at the terminus. Overall, the relative abundances measured for the neutral glycan ligands increased relative to those of the acidic ligands. These data suggest that the RBD possesses at least two glycan binding sites, with distinct preferences for neutral and acidic ligands.

### Sialic acid-dependent RBD-binding and viral entry

The results above suggest that SA-glycoconjugates are important for infection of cells by SARS-CoV-2. To test this hypothesis, we used ACE2-expressing HEK293 cells (Fig. 4a). When employing a trimeric RBD with a C-terminal mPlum fusion^31^, we observed robust binding of the SARS-CoV-2 RBD to ACE2^+^ cells, but no binding to ACE2^−^ cells (Fig. 4b). Pseudotyped SARS-CoV-2 lentivirus, encoding GFP, shows robust infection of ACE2^+^ 293 cells 24 hours after a one-hour incubation (Fig. 4c).

**Figure 4.**
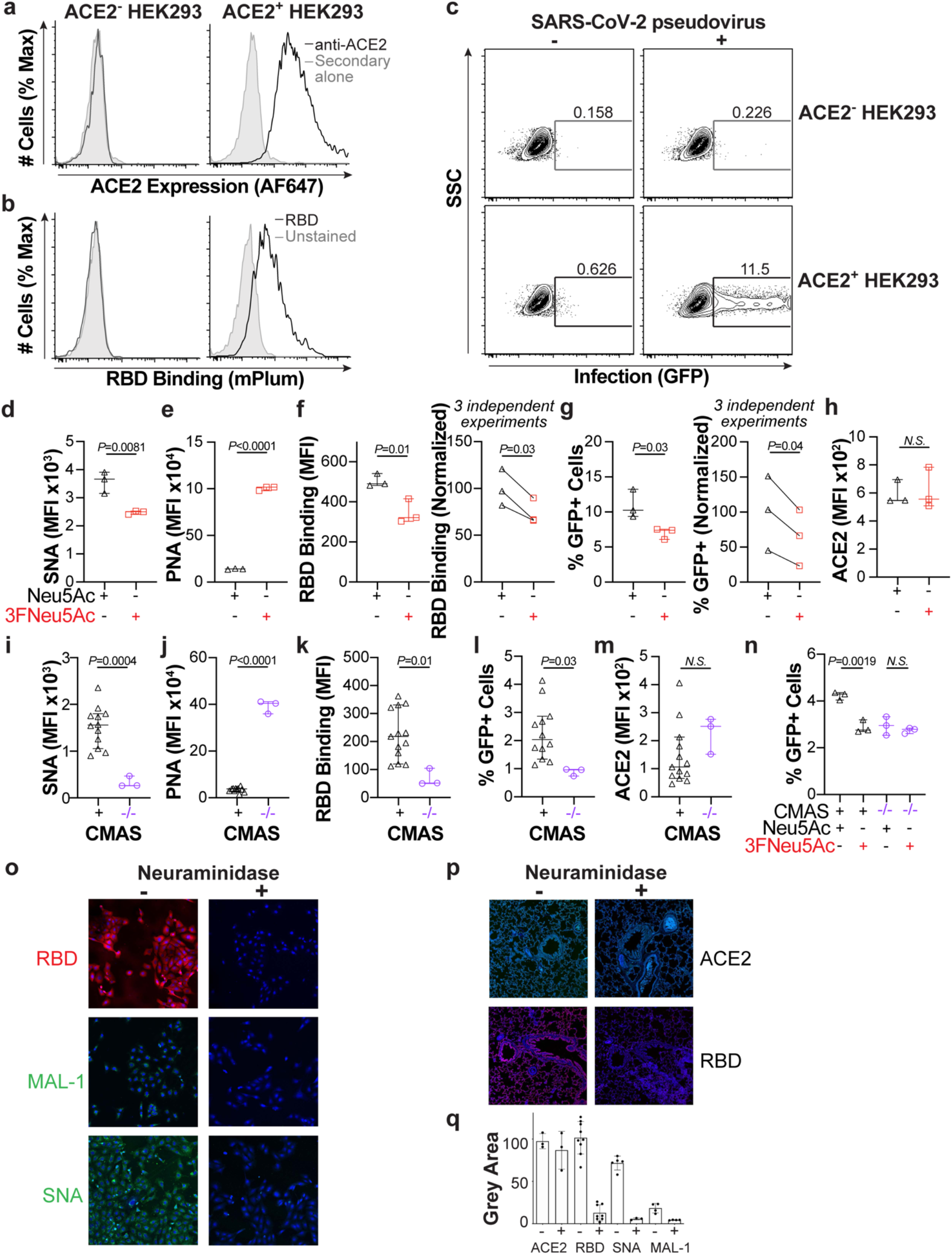
Decreasing sialic acid on ACE2^+^ cells decreases RBD binding and SARS-CoV-2 pseudotyped lentiviral infection. **a,b,** Expression of human ACE2 (**a**) and trimeric (**b**) and fluorescent RBD binding to WT (left) and ACE2^+^ (right) HEK293 cells as detected by flow cytometry. **c**, Flow cytometry gating of GFP-encoding pseudotyped SARS-CoV-2 lentivirus infection of WT (top) and ACE2^+^ (bottom) HEK293 cells. **d,e**, Changes in SA levels from 3FNeu5Ac treatment of HEK293 ACE2^+^ cells determined by SNA (**d**) and PNA (**e**) lectin staining by flow cytometry. **f**, 3FNeu5Ac treatment decreases RBD binding; shown are the mean fluorescence intensity values for three replicates within a single experiment (left) and the average of three independent experiments (right). The average RBD binding to control cells was set to 100%. **g,** 3FNeu5Ac treatment decreases SARS-CoV-2 pseudovirus infection of HEK293 ACE2^+^ cells; shown is % GFP^+^ cells for three replicates within a single experiment (left) and the averages of three independent experiments (right). **i,j,** Changes in SA levels within CMAS^−/−^ ACE2^+^ HEK293 cells determined by SNA (**i**) and PNA (**j**) lectin staining by flow cytometry. Each point represents the average of three replicates for each individual clone. **k-m,** Flow cytometry results for RBD binding (**k**), SARS-CoV-2 pseudovirus infection (**l**), and ACE2 expression (**m**) on CMAS^+^ versus CMAS^−/−^ ACE2^+^ HEK293 cells. Each point represents the average of three replicates for each individual clone. **n**, SARS-CoV-2 pseudovirus infection of CMAS^+^ versus CMAS^−/−^ ACE2^+^ HEK293 cells treated with or without 3FNeu5Ac. **o,p,** Pre-treatment of Vero-E6 cells (**o**) and ferret lung tissue sections (**p**) with neuraminidase decreases RBD binding. **q**, Quantification of fluorescence intensity for RBD binding and lectin staining for staining Vero-E6 cells and ferret lung tissue sections. Error bars represent ± standard deviation of three replicates. Statistical significance was calculated based on two-tailed unpaired Student’s *t*-test.

To test the role of SA in RBD binding and viral entry, we employed pharmacological, genetic, and enzymatic approaches to decrease the SA levels. ACE^+^ HEK293 Cells were first treated with the sialyltransferase (ST) inhibitor 3FNeu5Ac^32^, or its non-fluorinated analog as a control, for three days, which significantly decreased SA on the cells as measured by flow cytometry with the lectins SNA (Fig. 4d) and PNA (Fig. 4e). In 3FNeu5Ac-treated cells, RBD binding was consistently decreased (Fig. 4f) and pseudotyped lentivirus showed less infection (Fig. 4g). The decrease in infectivity was approximately 30–40%, which was consistent over numerous independent experiments. Using immunofluorescence staining of Vero-E6 cells, a decrease in RBD binding was also observed in cells treated with 3FNeu5Ac (Supplementary Fig. 10). Importantly, ST inhibition did not alter ACE2 levels (Fig. 4h and (Supplementary Fig. 10). Using a control lentivirus that does not encode the SARS-CoV-2 Spike protein, no differences were observed in viral entry upon 3FNeu5Ac treatment (Supplementary Fig. 11a).

We also genetically-ablated SA through isolation of a number of CMP sialic acid synthetase (CMAS) positive (CMAS^+^) and negative (CMAS^−/−^) clones by CRISPR/Cas9 within ACE2^+^ HEK293 cells, which were all tested in parallel to minimize concerns of clonal variability. Compared to the twelve isolated CMAS^+^ clones, the three CMAS^−/−^ clones exhibited greatly decreased SNA staining (Fig. 4i) and enhanced PNA staining (Fig. 4j), which is consistent with the abrogation of CMP-SA biosynthesis. RBD binding was decreased (Fig. 4k) and pseudotyped viral entry was likewise suppressed in the CMAS^−/−^ clones (Fig. 4l), for which there were no significant differences in ACE2 expression levels (Fig. 4m). A single CMAS^+^ and CMAS^−/−^ clone, with comparable ACE2 expression levels, were selected and fed with 3FNeu5Ac, with results showing that 3FNeu5Ac-treatment only decreased pseudotyped viral infectivity in the CMAS^+^ clone (Fig. 4n). The lack of an effect of 3FNeu5Ac in CMAS^−/−^ cells strongly suggests that the effects of 3FNeu5Ac on infectivity are due to the lower SA levels on the cells.

As a third method for reducing SA levels on cells, Vero-E6 cells (Fig. 4o) and Ferret lung tissue sections (Fig. 4p) were fixed and pre-treated with neuraminidase from *Vibrio cholerae* prior to assessing RBD binding by immunofluorescence staining. SARS-CoV-2 RBD binding was decreased by nearly 80% for both cells and tissue, despite no differences in ACE2 levels, with the expected abrogation of SA reported by SNA and MAA lectin staining (Fig. 5p). These results were consistent with three different sources of neuraminidase and, required overnight treatment with the neuraminidases (Supplementary Fig. 12).

**Figure 5.**
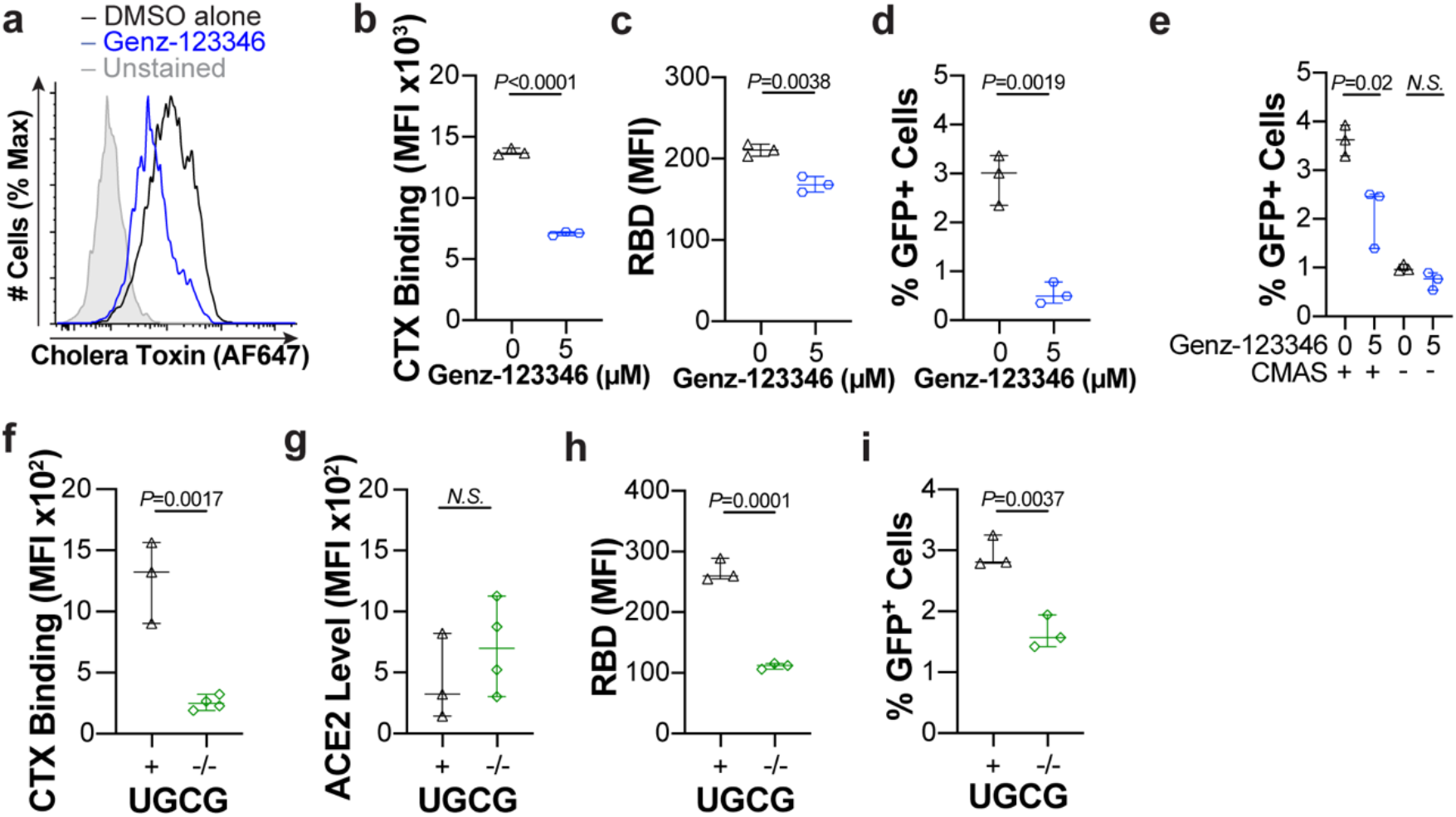
Pharmacological and genetic abrogation of glycolipids decrease RBD binding and SARS-CoV-2 viral infection. **a,b,** Changes in Cholera toxin (CTX) binding on ACE2^+^ HEK293 cells expressing after GENZ-123346 treatment determined by flow cytometry; representative results (**a**) and quantification of the MFI(**b**). **c**, RBD binding to ACE2^+^ HEK293 cells following GENZ-123346 treatment. **d**, SARS-CoV-2 pseudovirus infection in ACE2^+^ HEK293 cells following GENZ-123346 treatment. **e**, SARS-CoV-2 pseudovirus infection in CMAS^+^ and CMAS^−/−^ ACE2^+^ HEK293 cells following GENZ-123346 treatment. **f**, CTX staining of UGCG^+^ and UGCG^−/−^ ACE2^+^ HEK293 cell clones. **g**, ACE2 expression level on the UGCG^+^ and UGCG^−/−^ACE2^+^ HEK293 cell clones. **h**, RBD binding to UGCG^+^ and UGCG^−/−^ ACE2^+^ HEK293 cells. **i**, SARS-CoV-2 pseudovirus infection of UGCG^+^ and UGCG^−/−^ ACE2^+^ HEK293 cells. Error bars represent ± standard deviation of three replicates. Statistical significance was calculated based on two-tailed unpaired Student’s *t*-test.

### Glycolipids are critical for SARS-CoV-2 infection

We next investigated the potential for glycolipids to play a role in RBD binding and viral entry, given that gangliosides were the top hit in our binding assays. We used an inhibitor of UDP-glucose ceramide glycosyltransferase (UGCG), called GENZ-123346. ACE2^+^ HEK293 cells treated for two days with this inhibitor exhibited decreased Cholera toxin (CTX) staining by flow cytometry, which is indicative of decreased glycolipid levels on the cells (Fig. 5a,b). SARS-CoV-2 RBD binding (Fig. 5c) and pseudotyped viral infection (Fig. 5d) were tested in cells depleted of glycolipids with GENZ-123346, both of which showed significant decreases for the cells treated with the inhibitor. Moreover, GENZ-123346 failed to elicit an effect on pseudoviral entry with CMAS^−/−^ cells (Fig. 5e), suggesting the importance of SA-containing glycolipids. Infection of a control lentivirus not encoding the SARS-CoV-2 Spike protein did not show differences in infection upon treatment with GENZ-123346 (Supplementary Fig. 11b).

As a complementary approach, glycolipids were also depleted in cells by genetic ablation of UGCG by CRISPR/Cas9. Three UGCG^+^ and four UGCG^−/−^ clones were isolated, which stained positive and negative for CTX, respectively (Fig. 5f). Loss of UGCG did not significantly alter expression of ACE2 (Fig. 5g). A single UGCG^+^ and UGCG^−/−^ clone were selected with similar ACE2 expression levels for further testing. The UGCG^−/−^ cells showed decreased RBD binding (Fig. 5h) and were infected to a lesser extent by SARS-CoV-2 pseudotyped lentivirus (Fig. 5i). Overall, these results with genetic and pharmacological ablation of glycolipid biosynthesis suggest an important role for SA-containing glycolipids in SARS-CoV-2 infection of cells.

## Discussion

In this study, we leveraged CaR-ESI-MS, a sensitive and label- and immobilization-free assay, to study defined and natural glycan libraries to identify human glycan structures recognized by SARS-CoV-2 that may facilitate viral infection. Screening a defined glycan library against SARS-CoV-2 RBD revealed that several different classes of structures are recognized. Notably, RBD binds a variety of acidic glycans, with the highest preference for the oligosaccharides on the gangliosides GM1 and GM2. Screening of nanodiscs containing a mixture of gangliosides confirmed RBD binding to GM1 and GM2, as well as GM3. This is the first report of RBD binding to monosialylated gangliosides embedded in a lipid bilayer. The affinities of the monosialylated ganglioside oligosaccharides for SARS-CoV 2 RBD (e.g., 160 ± 40 μM for GM1 pentasaccharide at 25 °C) are similar and, in some instances, stronger than affinities reported for other ganglioside-binding viruses (e.g., respiratory syncytial virus and influenza virus strains) for which cell entry is SA-dependent^33,34^. It is noteworthy that, while other members of the Coronavirus family also bind glycolipids, they do so through their NTD; therefore, our observation of binding glycolipids through the RBD is an entirely new finding. Interestingly Hao *et al.* did not detect any binding of S_1_ to the glycans of glycosphingolipids, including gangliosides, in glycan microarray screening^4^. This absence of binding (with the array) might be due to the relatively low affinities of these interactions, or deleterious effects of glycan labeling on binding.

SARS-CoV-2 primarily affects the respiratory system, but it can also invade multiple organ systems. Recently, it was revealed that SARS-CoV-2 can invade the central nervous system (CNS)^35^. Infection of neurons by SARS-CoV-2 is highly relevant to our findings as gangliosides are prominent in human brain, at 10-to 30-fold higher levels than other organs^36^. Therefore, gangliosides possibly play an important role in mediating SARS-CoV-2 infection of the CNS.

Acidic human milk oligosaccharides (HMOs) were also observed at high relative abundances in CaR-ESI-MS screening, raising the possibility (based on an assumption HMOs can act as viral inhibitors) that breast fed newborns may be protected against SARS-CoV-2 infection. Indeed, in a study of 72 neonates breastfed by COVID-19 positive mothers, it was found that none tested positive for infection after 14 days^37^. Interestingly, some neutral glycans, including ABH blood group antigens (in particular A and H types 2, 3 and 4), were also found to exhibit moderate CaR-ESI-MS signal. Very recently, binding of RBD to blood group A was demonstrated^17^, which is consistent with our findings. Together, these may offer a clue as to why blood group A patients are at higher risk of hospitalization following SARS-CoV-2 infection^38^. Moreover, based on the competitive binding experiments with Neu5AcαMe, which competed away most of the sialoglycan but did not perturb binding of the neutral glycans, we suggest that the RBD may have at least two different glycan binding sites.

The affinities measured for the HS (**39** – **42**), which range from 200 μM to 400 μM (at 25°C), are similar to the *K*_d_ reported by Hao *et al*. for RBD and a long-chain heparin sample (240 μM, with MWs ranging from 17 kDa to 19 kDa)^4^ but weaker than the value reported by Liu *et al.* (1 μM) for heparin (average MW 15,700 Da)^5^. Notably, in the former SPR study, it was the RBD that was immobilized, while the glycan was immobilized in the latter. Therefore, simple avidity considerations likely explain this discrepancy in *K*_d_ values. Esko and co-workers reported that destruction of cellular glycosaminoglycans (GAGs) decreased SARS-CoV-2 infection, leading to the proposal that GAGs serve as a co-receptor. Notably, the affinities we report for ganglioside binding are as strong, or stronger, than for HS. Accordingly, our demonstration that reduction of SA or ganglioside levels on cells decreases RBD binding and infection of SARS-CoV-2 pseudotyped virus argues that sialoglycans may be equally as important as GAGs. Very recently, it was reported by others that, similar to what we observe, inhibition of UGCG diminished SARS-CoV-2 pseudotyped viral entry^39^. However, our findings go well beyond that by showing a direct binding interaction between the RBD and gangliosides, that inhibition of UGCG has no effect in cells lacking SA, and that genetic ablation of UGCG also causes a decrease in viral infection.

In summary, our findings suggest that sialylated glycans, which are abundant on all human cells, are bound through the RBD of SARS-CoV-2, thereby facilitating viral entry. These finding may have important implications in the tissue tropism of SARS-CoV-2, and could lead to new therapeutic approaches.

## Materials and Methods

### Ethics

The study protocol for acquisition of human tissue samples conformed to the ethical guidelines of the 1975 Declaration of Helsinki and was reviewed by the ethics review board at the University of Alberta (Pro00005105, Pro00085859 and Pro00035875).

The ferret lung tissues used in this study came from ferrets handled in an ABSL3 biocontainment laboratory. The research on these animals was conducted in compliance with the Dutch legislation for the protection of animals used for scientific purposes (2014, implementing EU Directive 2010/63) and other relevant regulations. The licensed establishment where this research was conducted (Erasmus MC) has an approved OLAW Assurance # A5051-01. Research was conducted under a project license from the Dutch competent authority and the study protocol (#17–4312) was approved by the institutional Animal Welfare Body. Animals were housed in Class III isolators in groups of 2.

### Proteins

Detailed information on the proteins used in this study and the preparation of stock solutions are given as Supplementary Information.

### Defined glycan library

The structures and MWs of the oligosaccharides contained within the defined library are described in Figure 1 and Table S1 (Supporting Information), along with their sources. With the exception of the HS compounds (**39**–**42**) and *N*-glycan standard (**141**), the glycans were divided into 16 MW-unique sub-libraries (Library A – P, Figure S1). To prepare the sub-libraries, a 1 mM stock solution of each glycan was first prepared by dissolving known mass of the glycan in ultrafiltered Milli-Q water (Millipore, Billerica, MA, USA). Aliquots of these solutions were then mixed, with ultrafiltered Milli-Q water, to give 50 μM stock solutions of each sub-library. All stock solutions were stored at –20 °C until needed.

### Synthesis

Additional details for preparation of these compounds is described in the supporting information.

#### *Synthesis of Methyl (5-acetamido-4,7,8,9-tetra-O-acetyl-3-dehydro-3,5-dideoxy-3-fluoro-5-β-D-glycero-D-galacto)onate* (143)

This compound was prepared similarly to as described elsewhere^40^.

#### *Synthesis of Methyl α-Neu5Ac* (146)

This compound was prepared similarly to as described elsewhere^41^.

### Natural glycan library

Two natural libraries of *N*-glycans, prepared from intestinal tissue (from a patient with Crohn’s Disease) and lung tissue, were screened against RBD. Lung tissue was taken post *ex vivo* perfusion of human lung (“right, peripheral”) of non-COVID19-related patient donor, then snap frozen and stored in liquid nitrogen. Complete details of the methods used to prepare the libraries are given as Supporting Information.

### Labeling of released *N*-glycans

The *N*-glycans released from RBD, lung or intestinal tissue were fluorescently labeled via reductive amination with 2-aminobenzamide (2-AB) (Sigma-Aldrich Canada)^42^. Complete details are given as Supporting Information.

### Mass Spectrometry

CaR-ESI-MS screening was performed in negative ion mode using a Q Exactive Ultra-High Mass Range Orbitrap mass spectrometer and the affinity measurements were carried out in positive ion mode on a Q Exactive Orbitrap mass spectrometer (Thermo Fisher Scientific, Bremen, Germany). Both instruments were equipped with a temperature-controlled nanoflow ESI (nanoESI) device; unless otherwise noted the solution temperature was 25°C^43^. CaR-ESI-MS measurements against ganglioside-containing nanodiscs were carried out in negative ion mode using a Synapt G2S quadrupole-ion mobility separation-time of flight (Q-IMS-TOF) mass spectrometer (Waters, Manchester, UK) nanoESI. NanoESI tips, with outer diameters (o.d.) of ~5 μm, were produced from borosilicate glass capillaries (1.0 mm o.d., 0.68 mm inner diameter) using a P-1000 micropipette puller (Sutter Instruments, Novato, CA). To perform nanoESI, a voltage of approximately +1 kV (positive mode) or −1 kV (negative mode) was applied to a platinum wire that was inserted inside the nanoESI tip (from the back) and in contact with the solution. An overview of the experimental and instrumental parameters used for the ESI-MS affinity measurements and CaR-ESI-MS screening experiments, along with data analysis procedures, are given as Supporting Information.

### Hydrophilic interaction-ultra high-performance liquid chromatography (HILIC-UHPLC) analysis of *N*-glycan libraries

The labeled *N*-glycans released from RBD and the intestinal and lung tissue were analyzed by HILIC-UHPLC on a Thermo Scientific™ Vanquish™ UHPLC system coupled with fluorescent (FLD) (Thermo Scientific, Waltham, MA, USA) and ESI-MS detectors (Thermo Q Exactive Orbitrap). Details of the experimental conditions and data analysis are provided as Supporting Information.

### Flow Cytometry

A 5-laser LSRFortessa™ X-20 Flow cytometer (BD Biosciences) was used for all flow cytometry data collection. Data was processed using FlowJo® Software Version 9.9.6.

### Production of Pseudovirus

GFP-encoding SARS-CoV-2 pseudotyped lentivirus was produced from HEK293T as reported^44^, using a plasmid encoding GFP (pCCNanoLuc2AeGFP), an envelope plasmid encoding SARS-CoV-2 spike protein (SARS-CoV-2), and a packaging plasmid encoding GagPol expression (pCRV1-NL-gagpol). Specifically, 1 × 10^6^ HEK293T cells were plated in a 6-well dish with 1.5 mL of DMEM growth media (Gibco) containing 10% Fetal Bovine Serum (FBS; Gibco), 100 U mL^−1^ Penicillin (Gibco), and 100 μg mL^−1^ Streptomycin (Gibco). The following day, pCCNanoLuc2AeGFP, pCRV1-NL-gagpol, and SARS-CoV-2 were mixed in a ratio of 0.7:0.7:0.4 with Opti-MEM® media (Gibco) to give 275 μL. TransIT®-LT1 reagent (Mirus Bio) was added according to the manufacturer’s protocol. Cells were incubated with this transfection mixture at 37 °C and 5% CO_2_ for 72 h. Following transfection, the supernatant was harvested, spun at 1,500 rcf at 4 °C for 5 min, then incubated at 4 °C in Lenti-X Concentrator (Clontech) for 1 h. The supernatant solution was spun at 1,500 rcf for 45 min at 4 °C. The final pellet was resuspended in 1/10th of the original volume of media using sterile PBS, titered, and stored at −80 °C.

### Pseudovirus Transduction

HEK293T cells expressing ACE2 (150,000)^45^ were plated in triplicate in a 24-well plate in 250 μL of DMEM growth media (Gibco), containing 10% Fetal Bovine Serum (FBS; Gibco), 100 U/mL Penicillin (Gibco), 100 μg mL^−1^ Streptomycin (Gibco), and 5μg mL^−1^ Blasticidin (InvivoGen). 10X concentrated pseudovirus (10 μL) was added to each 24-well and incubated for 1 or 8 h at 37 °C and 5% CO_2_. Cells were washed 1X in DMEM media and re-plated in the 24-well plate for a remaining 23 or 16 h to allow for expression of the GFP reported gene. Cells were removed from the plate via trypsin digest, centrifuged (300 rcf, 5 min), then resuspended in flow cytometry buffer (HBSS, 1% FBS, 500 μM EDTA). The viral titer of pseudovirus was determined by examining the % of GFP+ cells using flow cytometry.

### Production of Control Lentivirus

An empty lentiviral vector encoding mAmetrine was made using the Lentiviral backbone, RP172, as previously described^46^.

### Control Lentivirus Transduction

HEK293T cells expressing ACE2 (150,000)^45^ were plated in triplicate in a 24-well plate in 250 μL of DMEM growth media (Gibco), containing 10% Fetal Bovine Serum (FBS; Gibco), 100 U/mL Penicillin (Gibco), 100 μg mL^−1^ Streptomycin (Gibco), and 5μg mL^−1^ Blasticidin (InvivoGen). 10X concentrated control lentivirus (10 μL) was added to each 24-well and incubated for 1 hr at 37 °C and 5% CO_2_. Cells were washed 1X in DMEM media and re-plated in the 24-well plate for a remaining 23 or 16 h to allow for expression of the GFP reported gene. Cells were removed from the plate via trypsin digestion, centrifuged (300 rcf, 5 min), then resuspended in flow cytometry buffer (HBSS, 1% FBS, 500 μM EDTA). The viral titer of lentivirus was determined by examining the % of mAmetrine+ cells using flow cytometry.

### Lectin, Cholera toxin, and anti-ACE2 antibody staining by flow cytometry

Approximately 50,000 cells were seeded in a 96-well U-bottom flask and spun at 300 rcf for 5 min. The cell pellets were resuspended in a solution of in HBSS containing Calcium and Magnesium (Thermo Fisher) and 0.1% BSA (flow buffer), with either fluorescein-conjugated *Sambucus Nigra* Lectin (SNA, 1:1000, Vector Laboratories) or peanut agglutinin (PNA, 1:1000, Vector Laboratories) and incubated on ice for 20 min. After incubation, the cells were washed, and centrifuged (300 rcf, 5 min), and resuspended in flow buffer and analyzed by flow cytometry. The cells were stained with Fluorescein isothiocyanate (FITC)-conjugated cholera toxin B subunit (CTX, 1:1000, Sigma-Aldrich) at 1 μg mL^−1^ in flow buffer for 30 min at 4 °C. The cells were washed, resuspended in flow buffer, and analyzed by flow cytometry. For ACE2 staining, cells were resuspended in Goat anti-Human ACE2 antibody (1:500, R&D Systems) in the flow buffer. After a 20 min incubation at 4 °C, cells were centrifuged at 300 rcf for 5 min. The pellets were resuspended in a solution of anti-goat AF647 secondary antibody (Thermo Fischer) and incubated on ice for another 30 min. Cells were centrifuged at (300 rcf, 5 min) and the cell pellets were resuspended in the flow buffer. Differences in the median fluorescence intensity of AF647 were measured on the flow cytometer.

### Fluorescent SARS-CoV RBD Trimer Binding by flow cytometry

SARS-CoV-2 293T trimer containing the mPlum fluorophore was tested for cell binding at a final concentration of 10 μg/mL in DMEM containing 10% Fetal Bovine Serum. Cells were incubated at 4 °C for 45 minutes then centrifuged (300 rcf, 5 min). The cell pellets were resuspended in flow buffer and analyzed by flow cytometer for mPlum fluorescence in the PE-Cy5 channel.

### Immunofluorescent cell staining

Vero-E6 cells grown on coverslips were analyzed by immunofluorescent staining. Cells were fixed with 4% paraformaldehyde in PBS for 25 min at RT after which permeabilization was performed using 0,1% Triton in PBS. Subsequently, the coronavirus spike proteins were applied at 50 μg/ml for 1 hour at RT. Primary Strep-MAb classic chromeo-555 (IBA) and secondary Alexa-fluor-488 or −555 goat anti-mouse (Invitrogen) were applied sequentially with PBS washes in between. DAPI (Invitrogen) was used as nuclear staining. Samples were imaged on a Leica DMi8 confocal microscope equipped with a 10x HC PL Apo CS2 objective (NA. 0.40). Excitation was achieved with a Diode 405 or white light for excitation of Alexa488 and Alexa555, a pulsed white laser (80MHz) was used at 488 nm and 549 nm, and emissions were obtained in the range of 498-531 nm and 594-627 nm respectively. Laser powers were 10 – 20% with a gain of a maximum of 200. LAS Application Suite X was used as well as ImageJ for the addition of the scale bars. Where indicated, cells were treated with 2 mU of *Vibrio cholerae* (Sigma; #11080725001) or *Arthrobacter ureafaciens* (Sigma; #10269611001 or NEB; P0722) neuraminidase in 10 mM potassium acetate and 2.5 mg mL^−1^ Triton X-100, pH 4.2 at 37 °C O/N.

### Tissue staining

Serial sections of formalin-fixed, paraffin-embedded ferret lungs were obtained from the Department of Veterinary Pathobiology at Utrecht University and the Department of Viroscience at Erasmus University, respectively. Tissue was deparaffinized in xylene, rehydrated in a series of alcohol from 100%, 96% to 70%, and lastly in distilled water. Tissue slices were boiled in citrate buffer pH 6.0 for 10 min at 900 kW in a microwave for antigen retrieval and washed in PBS-T three times. The tissue slices were incubated with 3% BSA in PBS-T overnight at 4 °C. The next day, purified viral spike proteins (50 μg/ml) or antibodies were added to the tissues for 1 h at RT. With rigorous washing steps in between the secondary antibodies were applied for 45 min at RT. Where indicated tissue slides were treated with 2 mU of *Vibrio cholerae* neuraminidase in 10 mM potassium acetate and 2.5 mg mL^−1^ Triton X-100, pH 4.2 at 37 °C O/N.

### Pharmacological depletion of sialic acid

Stocks of peracetylated Neu5Ac or 3F-Neu5Ac were prepared in cell-culture grade dimethyl sulfoxide (DMSO) at 300 mM and stored at −20°C. 100,000 cells in 250 μL of media were seeded in a 24-well plate. The compound stocks were added to wells at a 1:1000 dilution to achieve a final concentration of 300 μM. Cells were incubated at 37 °C, 5% CO_2_ for 72 h to deplete cell surface SA levels.

### Generation of CMAS^−/−^and UGCG^−/−^Cells

crRNA was designed to target human CMAS (TGAGACGCCATCAGTTTCGA, Integrated DNA Technologies; IDT) or human UGCG (CCGATTACACCTCAACAAGA, IDT). HEK293T cells (500,000) expressing ACE2 were plated in a 6-well tissue culture plate in 1.5 mL of media. 24 hours later, gRNA (1 μM crRNA, 1 μM ATTO-550 labeled tracrRNA (IDT)) was boiled at 95 °C for 5 minutes. A solution of 20 pmol gRNA, 20 pmol Cas9 nuclease (IDT), 8 μL Cas9 PLUS reagent (IDT), 16 μL CRISPRMAX reagent (ThermoFisher) in 600 μL of Opti-MEM medium (Gibco) was added to cells and incubated at 37 °C, 5% CO_2_ for 24 h. The cells were removed from the 6-well plate via trypsin digestion, centrifuged (300 rcf, 5 min), then resuspended in 400 μL FACS buffer (HBSS containing 1% FBS and 500 μM EDTA). The top 5% ATTO-550 positive cells were sorted on a BD FACSMelody™ Cell Sorter into 96-well flat-bottom plates containing 200 μL of media at one cell per well and stored at 37 °C, 5% CO_2_. Approximately 2 weeks later, colonies were screened for CMAS^−/−^ cells on a flow cytometer using fluorescein-conjugated SNA and PNA, while UGCG^−/−^ cells were screened by FITC-conjugated cholera toxin B subunit (CTX, 1:1000, Sigma-Aldrich).

### Glycolipid Depletion

Stocks of Glucosylceramide Synthase Inhibitor (GENZ-123346, Toronto Research Chemicals) in cell-culture grade dimethyl sulfoxide (DMSO) were prepared at 5 mM and stored at −20 °C. 100,000 cells in 250 μL of media were seeded in a 24-well plate. The Genz stocks, or DMSO as a negative control, were added to wells at a 1:1000 dilution to achieve a final concentration of 5 μM. Cells were incubated with GENZ-123346 at 37 °C, 5% CO_2_ for 48 h to deplete glycolipids.

### Statistics and reproducibility

A Student’s *t*-test was used to assess statistical significance. All assays were conducted with replicates of *n* = 3.

## Data Availability

The authors declare that all data supporting the findings of this study are available within the paper and its supplementary information files. Please contact the corresponding authors (J.S.K) for access of raw data. This is stored electronically, and will be made available upon reasonable requests.

## Acknowledgements

The authors acknowledge the Canadian Glycomics Network, the Natural Sciences and Engineering Research Council of Canada, the Canada Foundation for Innovation, and the Alberta Innovation and Advanced Education Research Capacity Program for generous funding. MSM thanks Canadian Research Chairs for a tier 2 chair in chemical glycoimmunology. R.P.dV. is a recipient of an ERC Starting Grant from the European Commission (802780) and a Beijerinck Premium of the Royal Dutch Academy of Sciences. David J. Marchant (University of Alberta) is thanked for advice and critical reading the manuscript, Jayan Nagendran (University of Alberta) for generously providing the lung tissue, Kunimasa Susuki (University of Alberta) for preparation of lung homogenate, Paul D. Bieniasz (The Rockefeller University) and John L.M. Law (University of Alberta) for providing reagents (plasmids) for generating the pseudotyped virus and ACE2^+^ HEK293 cells, and Simonetta Sipione for advice related to glycolipids. mPlum-ER-3 was a gift from Michael Davidson (Addgene plasmid # 55966; http://n2t.net/addgene:55966;RRID:Addgene_55966). Plasmids for expression of SARS-CoV-2 Spike and RBD proteins were generously provided by Dr. Florian Krammer (Icahn School of Medicine at Mount Sinai, produced under NIAID NIAID Centers of Excellence for Influenza Research and Surveillance (CEIRS) contract HHSN272201400008C) and produced under CEIRS contract HHSN272201400004C awarded to S.M.T.

