## Supplementary Information for "Sialic acid-Dependent Binding and Viral Entry of SARS-CoV-2"

**Table of Content**

**Materials and Methods** ........ **S6**

Proteins ........ S6

Defined glycan library ........ S6

Phospholipid and glycosphingolipids ........ S7

Other reagents ........ S8

Synthesis of per-*O*-acetylated $3F_{ax}$-Neu5Ac ........ S8

Synthesis of protective derivative of methyl-α-Neu5Ac ........ S9

Synthesis of methyl-α-Neu5Ac ........ S9

Preparation of protein pellet from lung and intestinal tissues ........ S9

*N*-glycan sample preparation ........ S11

Preparation of gangliosides-containing nanodiscs ........ S11

Mass spectrometry ........ S12

CaR-ESI-MS library screening ........ S13

CaR-ESI-MS affinity ranking ........ S14

ESI-MS affinity measurements ........ S15

Labeling of *N*-glycan libraries ........ S16

Hydrophilic interaction-ultra high performance liquid chromatography (HILIC-UHPLC) analysis of natural *N*-glycan libraries ........ S17

Affinity ranking of *N*-glycan ligands detected by CaR-ESI-MS screening ........ S19

Assignment of glycan compositions of RBD glycoforms ........ S19

**Supplementary Tables** ........ **S21**

**Table 1.** Composition of the defined library and the corresponding sub-libaries used for CaR-ESI-MS screening ........ S21

**Table 2.** Summary of molecular weights of RBD glycoforms measured by ESI-MS ....... S27

**Table 3.** Summary of reported RBD *O*-glycosylation........ S31

**Table 4.** Summary of glycans identified by HILIC-UHPLC analysis of 2-AB labeled *N*-glycans released from RBD ........ S32

**Table 5.** Putative glycan compositions and relative abundances of the RBD glycoforms identified by ESI-MS ........ S35

**Table 6.** Dissociation constants ($K_d$) of glycan ligands for RBD........ S40

**Table 7.** Glycans identified by HILIC-UHPLC analysis of 2-AB labeled *N*-glycans released from lung tissue and results of CaR-ESI-MS screening against RBD........ S41

**Table 8.** Glycans identified by HILIC-UHPLC analysis of 2-AB labeled *N*-glycans released from intestinal tissue and results of CaR-ESI-MS screening against RBD ......... S42

**Supplementary Figures ........ S47**

**Figure 1.** Representative ESI mass spectra of S-protein and RBD........ S47

**Figure 2.** Structures of 140 defined glycans used in CaR-ESI-MS screening against RBD........ S48

**Figure 3.** Sub-libraries (A-O) of defined glycan library used for CaR-ESI-MS screening........ S50

**Figure 4.** CaR-ESI-MS screening against defined glycan libraries at 25 °C ........ S51

**Figure 5.** Zero-charge mass spectrum for an aqueous ammonium acetate solution of SARS-CoV-2 RBD........ S52

**Figure 6.** Chromatogram of 2-AB-labeled *N*-glycans released from RBD treated with PNGase F acquired using HILIC-UHPLC with fluorescence detection .......... S53

**Figure 7.** Dissociation constants of 10 RBD glycoforms .............................................. S54

**Figure 8.** Comparison of glycan affinities for RBD measured by ESI-MS and their relative abundances measured by CaR-ESI-MS screening .......................................... S55

**Figure 9.** CaR-ESI-MS screening ganglioside-containing nanodiscs against RBD ........ S56

**Figure 10.** Immunofluorescence staining in 3FNeu5Ac treated Vero-E6 cells. ................ S57

**Figure 11.** Infection of control lentivirus that does not encode SARS-CoV-2 in HEK293 cells expressing ACE2. ........................................................................................ S58

**Figure 12.** Immunofluorescence staining showing RBD binding in neuraminidase treated Vero-E6 cells ........................................................................................... S59

**Figure 13.** Structures of gangliosides (GM1, GM2, GM3, GD1a, GD2 and GT1b) and the phospholipid DMPC ......................................................................................... S60

**Figure 14.** Synthesis scheme of compounds **143** and **146** ............................................... S61

**Figure 15.** $^{1}$H NMR spectra of per-*O*-acetylated 3Fax-Neu5Ac (**143**) ............................. S62

**Figure 16.** $^{13}$C NMR spectra of per-*O*-acetylated 3Fax-Neu5Ac (**143**) ............................ S63

**Figure 17.** $^{1}$H NMR spectra of methyl-α-Neu5Ac (**146**) .................................................. S64

**Figure 18.** $^{13}$C NMR spectra of methyl-α-Neu5Ac (**146**) ................................................ S65

**Figure 19.** Influence of collision energy on glycan ligand release in CaR-ESI-MS screening .............................................................................................. S66

**Figure 20.** Reproducibility of CaR-ESI-MS glycan library screening data ....................... S67

**References ............................................................................................................. S68**

### Materials and Methods

**Proteins.** SARS-CoV-2 S protein[1] and RBD (consisting of residues 319-541 and a C-terminal hexa histidine tag)[2] were expressed in HEK cells and purified as described elsewhere. The lectins from *Sambucus nigra* (elder) and *Maackia amurensis*, lysozyme from chicken egg white and α-lactalbumin were purchased from Sigma-Aldrich Canada (Oakville, Canada). The P dimer of the Saga strain (GII.4, genotype, MW 69,734 Da) was produced as described elsewhere[3]. Each protein was dialyzed and concentrated against 200 mM ammonium acetate (pH 6.8) using Amicon 0.5 mL microconcentrator (EMD Millipore, Billerica, MA, USA) with a 10 kDa or 30 kDa MW cut-off and stored at -20 °C until used. The membrane scaffold protein (MSP) MSP1E1 (MW 27 494 Da) was expressed from the plasmid pMSP1E1 (Addgene, Cambridge, MA) and purified using a reported protocol[4]. The MSP1E1 protein was dialyzed into the Tris/HCl buffer (10 mM Tris, 100 mM NaCl, 1 mM EDTA, pH 7.4) and stored at -80 °C until needed. The concentrations of each stock solution were estimated by UV absorption at 280 nm.

**Defined glycan library.** The structures, MWs and sources of the purified oligosaccharides contained within the defined library are shown in Supplementary Figure 1,2, summarized in Supplementary Table 1. Glycans **1**, **2–5**, **7–14**, **33–38**, **43-77**, **79–83**, **85**, **88–91**, **93–103**, **105**, **109**, **112**, **113**, **116**, **122**, **127** and **129** were purchased from Elicityl SA (Crolles, France), **6**, **78**, **84**, **86**, **87**, **92**, **117**, **118**, **125**, and **131** were purchased from Dextra (Reading, UK), **104** and **123** were purchased from Carbosynth (San Diego, CA, USA) and **106–108**, **110**, **111**, **114**, **115**, **119–121**, **124**, **126**, **128**, **130**, **132**, and **133** were purchased from IsoSep (Tullinge, Sweden). Glycans **15–32**, **39–42** and **134-140** were prepared as described elsewhere[5-11]. The *N*-glycan **141** was purchased from Chemily Glycoscience (Atlanta, GA, USA).

With the exception of **39**–**42** and **141**, the glycans were divided into 16 MW-unique sub-libraries (*Library* A - P). To prepare the sub-libraries, 1 mM stock solutions of each glycan were first prepared by dissolving known mass of glycan in ultrafiltered Milli-Q water (Millipore, Billerica, MA, USA). Aliquots of these solutions were then mixed, with ultrafiltered Milli-Q water, to give 50 µM stock solutions of each sub-library. All stock solutions were stored at -20 °C until needed.

**Phospholipid and glycosphingolipids.** The gangliosides GM1 (β-D-Gal-(1→3)-β-D-GalNAc-(1→4)-[α-D-Neu5Ac-(2→3)]-β-D-Gal-(1→4)-β-D-Glc-ceramide, major isoforms d18:1-18:0 with MW 1545.88 and d20:1-18:0 with MW 1573.91 Da), GM2 (β-D-GalNAc-(1→4)-[α-D-Neu5Ac-(2→3)]-β-D-Gal-(1→4)-β-D-Glc-ceramide, major isoforms d18:1-18:0 with MW 1383.82 Da and d20:1-18:0 with MW 1411.86 Da) and GM3 (α-D-Neu5Ac-(2→3)-β-D-Gal-(1→4)-β-D-Glc-ceramide, major isoforms d18:1- 18:0 with MW 1180.74 Da and d20:1-18:0 with MW 1208.78 Da) were purchased from Cedarlane Labs (Burlington, Canada); GD1a (α-D-Neu5Ac-(2→3)-β-D-Gal-(1→3)-β-D-GalNAc-(1→4)-[α-D-Neu5Ac-(2→3)]-β-D-Gal-(1→4)-β-D-Glc-ceramide, major isoforms d18:1-18:0 with MWs 1836.97 Da and d20:1-18:0 with MW 1865.00 Da) and GT1b (α-Neu5Ac-(2→3)-β-D-Gal-(1→3)-β-D-GalNAc-(1→4)-[α-Neu5Ac-(2→8)-α-Neu5Ac-(2→3)]-β-D-Gal-(1→4)-β-D-Glc-ceramide, major isoforms d18:1-18:0 with MW 2128.07 Da and d20:1-18:0 with MW 2156.10 Da) were purchased from Sigma-Aldrich Canada (Oakville, Canada); GD2 (β-D-GalNAc-(1→4)-[α-D-Neu5Ac-(2→8)-α-D-Neu5Ac-(2→3)]-β-D-Gal-(1→4)- β-D-Glc-ceramide, major isoforms d18:1-18:0 with MW 1674.92 Da and d20:1-18:0 with MW 1702.95 Da) was purchased from MyBioSource Inc. (San Diego, CA). The phospholipid DMPC (1,2-dimyristoyl-sn-glycero-3-phosphocholine, MW 677.50 Da) was purchased from Avanti Polar Lipids (Alabaster, AL) (Supplementary Fig. 13). Stock solutions of each ganglioside (1 mM) and DMPC (20 mM) were prepared in HPLC grade methanol/chloroform (1:1, v/v, Thermo Fisher, Ottawa, Canada).

**Other reagents.** Select-Fluor and $NaIO_4$ was purchased from Sigma-Aldrich Canada (Oakville, Canada) and $D_2O$ was purchased from Deutero GmbH (Kastellaun, Germany).

**Synthesis of per-*O*-acetylated 3F$_{ax}$-Neu5Ac**. Synthesis and characterization of *N*-Acetyl-2,3-didehydro-2-deoxyneuraminic acid (DANA) were previously reported by Cairo and co-workers[12]. DANA (350 mg, 0.75 mol) was dissolved in $CH_3NO_2$ (3.5 mL) and $H_2O$ (500 µL) with Select-Fluor (1.1 g, 3.0 mol) and the reaction mixture was stirred at room temperature for 2 d. The reaction was monitored by TLC and after completion, the reaction mixture was quenched with saturated aq. $NaHCO_3$. The compound was extracted with EtOAc (2x20 mL) and then crude was purified by silica column chromatography (EtOAc/Hexane, 7:3) to afford of **142a** (125 mg, 38 %) and **142b** (100 mg, 30 %). A solution of **142a** (100 mg, 0.02 mol) in anhydrous pyridine (2 mL) was stirred with acetic anhydride (1.5 mL) at 0 °C for 5 h under argon atmosphere. The reaction mixture was then warmed to room temperature and stirred overnight (Supplementary Fig. 2a). After the reaction completion, the solution was concentrated and co-evaporated with toluene. The product was then purified with silica column chromatography (EtOAc/ Hexane, 7:3) to afford desired compound **143** (100 mg, 92 %). $^1$H NMR (500 MHz, $CDCl_3$) δ 5.59 (ddd, *J* = 28, 11, 2.5 Hz, 1H), 5.38–5.32 (m, 2H), 5.36 (dt, *J* = 6.5, 2.5 Hz, 1H), 4.96 (dd, *J* = 49, 2.5 1H), 4.58 (dd, *J* = 12.5, 2.5 Hz, 1H), 4.29–4.14 (m, 3H), 3.87 (s, 3H), 2.20 (s, 3H), 2.19 (s, 3H), 2.14 (s, 3H), 2.07 (s, 3H), 2.06 (s, 3H), 1.95 (s, 3H) (Supplementary Figure 3); $^{13}$C NMR (125 MHz, $CDCl_3$) δ 170.62, 170.52, 170.49, 170.24, 167.06, 165.06, 87.49, 71.82, 71.27, 67.98, 62.06, 53.47, 45.76, 31.55, 23.27, 22.61, 20.85, 20.77, 20.63, 20.50, 14.08 (Supplementary Figure 4); HRMS (ESI-MS) calculated for *m/z* $[M+Na]^+$ cald for $C_{22}H_{30}FNNaO_{14}$: 574.1543, found: 574.1538.

**Synthesis of protected derivative of methyl-α-Neu5Ac.** Synthesis of per-*O*-acetylated methyl ester glycosyl chloro derivative (**144**) was achieved in quantitative yield in three-step from the *N*-acetyl neuraminic acid through following a procedure reported by Daskhan and co-workers[13]. A solution of glycosyl chloride **144** (450 mg, 0.88 mmol) and 4Å powdered molecular sieves (1 gm) in anhydrous MeOH (12 mL) was stirred at room temperature under an argon atmosphere. After 15-20 minutes, AgOTf (453 mg, 1.76 mmol) was added to the reaction mixture and stirred for 4 h in the dark. The crude product was filtered through a ceiled pad and washed with $CH_2Cl_2$ (3x150 mL), brine (50 mL), water (2x50 mL) and aqueous $Na_2S_2O_3$ (10%, m/v). The crude product was purified by silica column chromatography (EtOAc/Hexane, 9.5:0.5) to afford of compound **145** (323 mg, 72%) as a colourless solid. TLC: EtOAc:hexane = 9.5:0.5. $R_f$=0.42 (EtOA). $^1$H NMR (500 MHz, $CDCl_3$) δ 5.47-5.44 (m, 1H), 5.35 (dd, $J$ = 8.5, 2.5 Hz, 1H), 5.13 (d, $J$ = 9.5 Hz, 1H), 4.91-4.85 (m, 1H), 4.34 (dd, $J$ = 12.5, 2.5 Hz, 1H), 4.16-4.13 (m, 1H), 4.09 (dd, $J$ = 10.5, 5.5 Hz, 1H), 3.83 (s, $CO_2Me$, 3H), 3.34 (s, OMe, 3H), 2.59 (dd, $J_{4\text{-}3}$eq 5.0, $J_3$ax-$_3$eq 12.5 Hz, 1H, H3$_{eq}$) 2.17 (s, 3H), 2.16 (s, 3H), 2.06 (s, 3H), 2.05 (s, 3H), 1.96 (dd, $J_{4\text{-}3}$ax = 13.0, 25.5 Hz, 1H, H3$_{ax}$), 1.90 (s, 3H, NCOC$H_3$); $^{13}$C NMR (125 MHz, $CDCl_3$) δ 171.01, 170.66, 170.18, 170.14, 170.02, 168.16, 98.98, 72.47, 69.07, 68.43, 67.32, 62.39, 52.76 ($CO_2Me$), 52.48 (OMe), 49.51, 37.91, 23.23, 21.13, 20.87, 20.78. 1D NMR data matched with previously reported literature[14].

**Synthesis of methyl-α-Neu5Ac.** Methyl 5-acetamido-3,5-dideoxy-d-glycero-α-D-galacto-2-onulopyranose (**145**) (100 mg, 0.19 mmol) was dissolved in dry MeOH (10 mL), then NaOMe was added to the solution and pH of the reaction mixture was adjusted to 9.0-9.5. The mixture was left stirring at room temperature for overnight under an Ar atmosphere. Solvent was evaporated under reduced pressure. The crude product was subjected to de-esterification dissolving in 2N NaOH (2 mL) and stirred for 24 h at room temperature. Reaction mixture was neutralized with addition of amberlite IR-120 resin ($H^+$), filtered and lyophilized. The crude product was dissolved in $H_2O$ (0.5 mL) and purified by BioGel P-2 column eluted with water to compound **146** as a colourless solid after lyophilization. Yield: (56.9 mg, 89%). TLC: EtOAc:acetic acid:MeOH:$H_2O$ = 6:1:2:1. R*f* = 0.25. $^1$H NMR (500 MHz, $D_2O$): δ 3.86 (ddd, 1H), 3.84 (dd, $J$ = 9.5, 2.0 Hz, 1H), 3.78 (dd, $J$ = 20.0, 10.0 Hz, 1H), 3.68 (dd, $J$ = 9.5, 2.0 Hz, 1H), 3.65-3.64 (m, 1H), 3.62 (dd, $J$ = 12.5, 6.5 Hz, 1H), 3.56 (dd $J$ = 9.5, 2.0 Hz, 1H), 3.31 (s, OMe, 3H), 2.69 (dd, $J_{4\text{-}3}$eq 4.5, $J_3$ax$_{\text{-}3}$eq 12.5 Hz, 1H, H3$_{eq}$), 2.01 (s, 3H, NCOC$H_3$) 1.60 (dd, $J_{4\text{-}3}$ax = 12.5, 24.5 Hz, 1H, H3$_{ax}$) (Supplementary Fig. 17); $^{13}$C NMR (125 MHz, $D_2O$) δ 174.29, 171.58, 101.50, 73.44, 72.51, 69.08 (2C), 63.47, 52.72, 52.48 (OMe), 40.22, 23.30 (Supplementary Fig. 18). 1D NMR data matched with previously reported literature[15].

**Preparation of protein pellet from lung and intestinal tissues.** Protein extraction from lung or intestinal tissue from a patient with Crohn's disease was carried out using an adaptation of a previously reported protocol[16]. Briefly, the harvested sample was snap-frozen immediately and then stored at -80 $^o$C until homogenization in ice cold water using PowerGen 125 homogenizer (Fisher Scientific, Ottawa, ON, Canada). Then 2.7 times the aqueous sample volume of methanol and 1.3 times the aqueous sample volume of chloroform was sequentially added to the homogenized sample and mixed vigorously at room temperature. The homogenate was centrifuged at 3000 x g for 10 min followed by removal of supernatant. The excess methanol/chloroform from the pellet was removed under a stream of nitrogen gas for 2 min. After addition of 0.6 M TRIS buffer (pH 8.5, adjusted with dilute acetic acid), the sample was replaced under a stream of nitrogen and evaporated. The sample was then re-suspended with 200 mM ammonium acetate, followed by centrifuge at 4000 rpm, 10 min. The supernatant was lyophilized and stored at -20 $^o$C until needed.

***N*-glycan sample preparation.** To prepare the *N*-glycan libraries, 400 µg of SARS-CoV-2 RBD or 1 g of protein pellet extracted from lung or intestinal tissue was dissolved in 500 µL of 8M urea in 100 mM Tris-HCl (pH 8.0) containing 3 mM EDTA, and incubated at room temperature for 1 h. The denatured RBD were then reduced with 10 µL of 500 mM dithiothreitol (DTT, Sigma-Aldrich, Canada) at room temperature for 1 h followed by alkylation with 23 µL of 500 mM iodoacetamide (IAA, Sigma-Aldrich, Canada) at room temperature for 20 min in the dark. The reaction was quenched by adding 10 µL of 250 mM DTT, and the solution buffer exchanged using a PD MiniTrap G-25 column (GE Healthcare, Buckinghamshire, UK) according to the manufacturer's instructions. The glycoproteins solution was subsequently digested with trypsin/chymotrypsin [Substrate/Enzyme (weight/weight) = 50] in 50 mM ammonium bicarbonate (pH 8.0) for 18 h at 37 °C. The reaction was quenched by heat inactivation at 100 °C for 10 min.

Released *N*-glycans were liberated by incubation of resulting glycopeptides with amidase PNGase F (New England BioLabs, MA, USA) at 37 °C for 18 h and purified using porous graphitized carbon (Hypercarb cartridges, 100 mg, 1 mL volume, Thermo Fisher Scientific). The porous graphitized carbon cartridge was pre-equilibrated with 1 mL 80% acetonitrile containing 0.1% trifluoroacetic acid (TFA) followed by 2 mL of water. The samples were added to the cartridges followed by washing with 2 mL of water. *N*-glycans were then eluted with 1 mL of 25% acetonitrile containing 0.1% TFA. The lyophilized sample was stored at -20 $^{o}$C until needed.

**Preparation of ganglioside-containing nanodiscs**. Nanodisc (ND) composed of DMPC and a mixture of GM1, GM2, GM3, GD1a, GD2 and GT1b (each nominally 1% of total lipid) was prepared according to the protocol described by Sligar and coworkers[4]. Briefly, DMPC and the six gangliosides were diluted in methanol at the desired molar ratios. The lipids were dried under gentle stream of $N_2$ to form a lipid film, and then re-suspended in a buffer (pH 7.4) of 20 mM

TrisHCl, 0.5 mM EDTA, 100 mM NaCl and 25 mM sodium cholate (Sigma-Aldrich Canada). The membrane scaffold protein MSP1E1 was then added to the mixture at a 1:100 [MSP1E1:(gangliosides + DMPC)] molar ratio. The ND self-assembly process was initiated by adding pre-washed Bio-Beads (Bio-Rad, Mississauga, Canada) and the mixture was incubate for 3 h at room temperature using an orbital shaker. Following incubation, the supernatant, which contained the NDs, was removed and purified by gel-filtration chromatography using a Superdex 200 10/300 size-exclusion column (GE-Healthcare Life Sciences, Piscataway, NJ) equilibrated with 200 mM ammonium acetate (pH 6.8). Finally, the fraction corresponding to the NDs was collected, concentrated, dialyzed into 200 mM ammonium acetate (pH 6.8) using an Amicon microconcentrator (EMD Millipore, Billerica, MA) with a 30 kDa MW cut off and stored at -20 °C until used. The concentrations of the ND stock solutions were estimated from the concentration of MSP1E1, which was measured by UV absorption at 280 nm, and taking into account that each ND consists of two copies of MSP1E1.

**Mass spectrometry**. The CaR-ESI-MS glycan library screening measurements were performed in negative mode using a Q Exactive Orbitrap mass spectrometer with Ultra High Mass Range (Q Exactive UHMR, Thermo Fisher Scientific). The capillary temperature was 200 °C, and the S-lens RF level was 100; an automatic gain control target of 5 $\times 10^5$ and maximum injection time of 200 ms were used. The resolving power for full MS and high energy CID (HCD) were set to 3125 and 25,000, respectively. HCD spectra were acquired using collision energies ranging from 10 eV to 60 eV. The ESI-MS affinity measurements were performed in positive ion mode on a Q Exactive Orbitrap mass spectrometer (Thermo Fisher Scientific). The capillary temperature was 200 °C and the S-lens RF level was 100; an automatic gain control target of 5 $\times 10^5$ and maximum injection time of 200 ms were used. The resolving power is 17,500. For both instruments, data acquisition and pre-processing was performed using Xcalibur version 4.1; ion intensities were extracted using in-house software (SWARM)[17]. The CaR-ESI-MS analysis of ganglioside nanodiscs was carried out in negative ion mode using a Synapt G2S quadrupole-ion mobility separation-time of flight (Q-IMS-TOF) mass spectrometer (Waters, Manchester, UK). The source temperature was 60 °C; the cone voltage was 60 V, the trap voltage was 5 V (for MS) or 50 V (for CID), and the transfer voltage was 2 V. Argon was used for CID at a Trap ion guide pressure of $1.42 \times 10^{-2}$ mbar. Data acquisition and processing were performed using MassLynx software (version 4.1).

All three instruments were equipped with a modified nanoflow ESI (nanoESI) source. NanoESI tips with an outer diameter (o.d.) of ~5 μm were pulled from borosilicate glass (1.0 mm o.d., 0.78 mm inner diameter) with a P-1000 micropipette puller (Sutter Instruments, Novato, CA). A platinum wire was inserted into the nanoESI tip, making contact with the sample solution. A voltage of approximately +1 kV (positive mode) or -1 kV (negative mode) was applied to the platinum wire. The temperature of the solution in the nanoESI tip was controlled using a home-built temperature-controlled device[18].

**CaR-ESI-MS library screening.** For all screening experiments, aqueous solutions composed of the volatile buffer ammonium acetate (100 mM, pH 6.9), RBD (13 μM) and sub-library (50 nM of each glycan) were used. The SNA lectin, which has strict preference for α2-6 linked sialosides, served as $P_{ref}$ for libraries containing neutral glycans and those containing Neu5Acα2-3 and Neu5Acα2-8 (Libraries A-M, O and P) and MAA, which is specific for certain α2-3 linked sialosides, was used for Library N, which contained Neu5Acα2-6 glycans. A range of HCD energies (10 – 60 eV) were used to perform ligand release form the RBD. The onset of ligand release was observed at energies between 22 eV and 32 eV. Although there were a few exceptions, the trends in relative abundances of the released ligands were similar at collision energies up to

~50 eV (Supplementary Fig. 19) . Also, the absolute signal for released ligands increased with energy up to 50 eV. At higher energies, the abundances of released ligands were found to decrease due to the secondary fragmentation. Consequently, 50 eV was selected as the optimal collision energy for library screening. The reproducibility of the CaR-ESI-MS screening results using the selected experimental conditions was assessed using Library D (Supplementary Fig. 20). The relative standard deviations of the 18 glycan ligands in the library ranged from 2% to 27% (4 replicates), which is in the acceptable-to-excellent range for analytical methods[19]. Moreover, the trend in relative abundances of released ligands remained constant across the 4 measurements.

**CaR-ESI-MS affinity ranking.** To rank affinities of released ligands identified across all sub-libraries, the relative abundance ($Ab_{rel}$) of ligands were calculated from eq S1:

$$Ab_{rel}(\mathrm{L}_j) = Ab(\mathrm{L}_j)/Ab_{\mathrm{P33}} \qquad \text{(S1)}$$

where $\mathrm{L}_j$ is a given glycan in the defined library, $Ab(\mathrm{L}_j)$ is the total, charge-normalized abundance of the deprotonated ions of $\mathrm{L}_j$ and any adducts (e.g., chloride and nitrate adduct) detected and $Ab_{\mathrm{P33}}$ is the total, charge-normalized abundance of the most abundant RBD glycoform (P33, MW 31,926 Da).

**ESI-MS affinity measurements.** Dissociation constants ($K_\mathrm{d}$) glycan ligands for RBD were measured by the direct ESI-MS binding assay[20]. As the number of RBD glycan binding sites is not known and no RBD species with multiple bound ligands were detected, the ESI-MS data were analyzed assuming a single site. For a monovalent protein-ligand (PL) interaction (eq S2), the affinity ($K_\mathrm{d}$, eq S3) can be calculated from the ratio ($R$) of total $Ab$ of L-bound to free P ions (eq S3) measured by ESI-MS:

$$\mathrm{P} + \mathrm{L} \rightleftharpoons \mathrm{PL} \qquad \text{(S2)}$$

$$K_d = \frac{[P][L]}{[PL]} = \frac{[L]_0}{R} - \frac{[P]_0}{R+1} \quad (S3)$$

$$R = \frac{Ab(PL)}{Ab(P)} = \frac{[PL]}{[P]} \quad (S4)$$

where $[P]_0$ and $[L]_0$ are initial concentrations of P and L, respectively. The abundance ratio $R$ measured by ESI-MS is taken to be equal to the equilibrium concentration ratio in the solution. Because RBD consists of multiple species with distinct glycan compositions (referred to here as $P_x$, where x = 1-77) and glycosylation can, in principle, influence binding, the affinities for L binding to individual RBD species (eqs S5a-x) were determined:

$$P_1 + L \rightleftharpoons P_1L \quad (S5a)$$

$$\vdots \qquad \qquad \vdots$$

(S4x)

$$P_x + L \rightleftharpoons P_xL \quad (S5x)$$

The corresponding equations of mass balance equations are:

$$[L]_0 = [L] + \sum_x [P_xL] \quad (S6a)$$

$$[P_1]_0 = [P_1] + [P_1L] \quad (S6b)$$

$$[P_2]_0 = [P_2] + [P_2L] \quad (S6c)$$

$$\vdots \qquad \qquad \vdots$$

$$[P_x]_0 = [P_x] + [P_xL] \quad (S6x)$$

where $[P_x]_0$ is the initial concentration of a given $P_x$ species. Initial concentrations of individual RBD species were estimated from their relative abundances measured by ESI-MS, assuming uniform response factors, eq S7:

$$[P_x]_0 = \frac{Ab(P_x)}{\sum_x Ab(P_x)}[P]_0 \quad (S7)$$

The affinity of given $P_x$ species ($K_{dx}$) can be calculated from eq S7:

$$K_{dx} = \frac{[P_x][L]}{[P_xL]} = \frac{[L]_0}{R_x} - \frac{1}{R_x}\sum_x \frac{R_x[P_x]_0}{(1+R_x)} \quad \text{(S8)}$$

where $R_x$ is the total abundance ratio of ligand-bound and free $P_x$ ions, eq S8:

$$R_x = \frac{Ab(P_xL)}{Ab(P_x)} = \frac{[P_xL]_{eq}}{[P_x]_{eq}} \quad \text{(S9)}$$

When detected, the contribution of non-specific $P_x$-L binding, which occurred during ESI, to the mass spectrum was corrected using the $P_{ref}$ method[21].

In some instances, signal for one ligand-bound $P_x$ complex overlapped with signal for another (free) $P_x$ species. Spectral overlap was corrected for by considering the abundance ratio of the two $P_x$ species (e.g. $P_x$ and $P_{x+1}$) in the absence of L ($r$, eq S10) and assuming that $P_x$ and $P_{x+1}$ exhibit identical affinities for L:

$$r = \frac{Ab(P_{x+1})}{Ab(P_x)} \quad \text{(S10)}$$

The corresponding $r$ value was then used to calculate the true $R_x$ value ($R_{x,cor}$), corrected for spectral overlap:

$$R_{x,cor} = \frac{Ab(P_xL)}{Ab(P_x)} - r \quad \text{(S11)}$$

which can be used in eq S8 to calculate $K_{dx}$. Notably, there is good agreement between $R_x$ values measured in the absence of spectra overlap and those corrected for overlap, which indicates that glycosylation has at best a minor influence on glycan ligand binding.

**Labeling of *N*-glycan libraries**. The *N*-glycans released from RBD, lung or intestinal tissue were fluorescently labeled via reductive amination with 2-aminobenzamide (2-AB) (Sigma-Aldrich Canada). The 2-AB and sodium cyanoborohydride were dissolved in 70:30 (%, v/v) DMSO–acetic acid to furnish concentrations of 0.37 and 1 M, respectively. The labeling solution was added to the *N*-glycan sample followed by incubation at 65 °C for 3 h. The excess labeling reagent was removed by 30 mg Oasis HLB cartridge (Waters, Milford, MA, USA). The cartridge was conditioned by 2 × 1 mL of acetonitrile. The labeled *N*-glycans were diluted tenfold with acetonitrile and added to a conditioned cartridge. The loaded cartridge was washed with 2 × 1 mL of acetonitrile and eluted with 2 × 400 µL of 20% acetonitrile. The purified *N*-glycans were lyophilized and stored at −20 °C prior to analysis.

**Hydrophilic interaction-ultra high performance liquid chromatography (HILIC-UHPLC) analysis of natural *N*-glycan libraries.** The labeled *N*-glycans released RBD and the intestinal and lung tissue were analyzed by HILIC-UHPLC on a Thermo Scientific™ Vanquish™ UHPLC system coupled with fluorescent (FLD) (Thermo Scientific, Waltham, MA, USA) and ESI-MS detectors (Thermo Q Exactive Orbittrap). Compounds separation was achieved using a Waters Acquity UPLC BEH Glycan column (1.7 µm, 2.1 × 150 mm, Waters). The eluents were ammonium formate 100 mM (pH 4.5) (solvent A) and acetonitrile (solvent B). The separation was performed at 60 ºC. The following gradient was used for both FLD and MS detection: t = 0 min, 25% solvent A (0.2 mL/min); t = 175, 35% solvent A (0.2 mL/min); t = 210, 45% solvent A (0.2 mL/min); t = 210.1, 100% solvent A (0.2 mL/min); t = 224, 100% solvent A (0.2 mL/min); t = 225, 25% solvent A (0.2 mL/min); t = 240, 25% solvent A (0.2 mL/min). The excitation and emission wavelengths were set at 330 nm and 420 nm respectively. During LC-MS analysis, following parameters were used: probe heater temperature of 250 ºC, sheath gas flow rate of 40 arbitrate units (arb), aux gas flowrate of 10 arb, capillary temperature of 275 ºC and spray voltage of 3.5 kV. The mass spectra was acquired in positive mode with m/z range of 250-3000 at resolution of 70 000. The automatic gain control (AGC) target was set at $1\times10^6$ and maximum injection time of 100 ms were used. HCD mass spectra were acquired in the data-dependent mode for the 5 most abundant ions with resolution of 17,500. AGC target, maximum injection time, and isolation window were set at $2 \times 10^5$, 200 ms and 2.0 *m/z*, respectively. HCD normalized collision energy was 27%. The data were recorded by Xcalibur (Thermo, Version 4.1) and interpreted using GlycoWorkbench software.

The glycan composition corresponding to each peak identified by HILIC-UHPLC analysis was preliminarily assigned according to the MS data and biosynthetic pathways of *N*-glycan[22]. The structures of separated isomers were then assigned according to the measured retention times[23,24]. The fractional abundance ($F_j$) of a given labeled *N*-glycan within the mixture was determined from the relative fluorescent signal in the chromatogram, eq S12:

$$F_j = \frac{AF_j}{\sum_j AF_j} 100\% \tag{S12}$$

where $AF_j$ is the peak area of glycan *j*. In cases where a chromatographic peak contained multiple *N*-glycan species with distinct molecular weights (MWs), analysis of corresponding extracted-ion chromatogram (EIC) enabled the determination of the fractional abundance ($f_j$) of individual glycans (or mixture of compositional isomers) that comprise a given peak, eq S13:

$$f_j = \frac{\sum_n A_j^{n+}}{\sum_j \sum_n A^{n+}{}_{j,co-eluting}} \tag{S13}$$

where $A_j$ is the peak area of the monoisotopic ion of glycan *j* (summed over all charge states, *n*) and $A_{j,co\text{-}eluting}$ is sum of all monoisotopic ion peak areas corresponding to co-eluting *N*-glycan species. The fractional abundance of each glycan *j* ($AF_j$) within the chromatographic peak was calculated using eq S14:

$$AF_j = f_j \sum_j AF_{j,co-eluting} \qquad \text{(S14)}$$

where $AF_{j,co\text{-}eluting}$ corresponds to the total fluorescent signal of co-eluted *N*-glycans.

**Affinity ranking of *N*-glycan ligands detected by CaR-ESI-MS screening.** The ligand affinities (individual glycans or mixtures of compositional isomers) for RBD detected by CaR-ESI-MS were ranked based on their charge normalized fractional abundances ($FAb_j$) and fractional abundances determined by HILIC-UHPLC ($F_j$). For each glycan ligand (or mixture), the relative affinity ($K_{rel,j}$),which was calculated using eq S15:

$$K_{rel,j} = \frac{FAb_j}{F_j} \qquad \text{(S15)}$$

The ligand affinity ranking values ($r_j$) reported for each GBP (Figure 5) correspond to the $K_{rel,j}$ values normalized to the maximum value $K_{rel,max}$, eq S16:

$$r_j = \frac{K_{rel,j}}{K_{rel,max}} \qquad \text{(S16)}$$

**Assignment of glycan compositions of RBD glycoforms.** The data were analyzed using the measured MWs of intact protonated RBD. The MW of deglycosylated RBD was calculated based on the elemental composition ($C_{1170}H_{1744}N_{318}O_{335}S_9$) corresponding to residues 319-541 of the S-protein, plus the hexa His tag and 4 disulfide bonds. Possible glycan compositions were simulated for the numbers of *N*-acetylhexosamines (N ≡ HexNAc*, N*-acetylgalactosamine and *N*-acetylglucosamine), hexoses (H ≡ Hex, glucose and galactose), fucoses (F ≡ Fuc) and *N*-acetylneuraminic acids (S ≡ Neu5Ac). Possible values of N, H, F and S were calculated by considering the composition of *N*-glycans established by HILIC-UHPLC-FLD/ESI-MS and reported *O*-glycans (Supplementary Tables S3-S4)[25-28]. Possible MWs of RBD glycoforms were then calculated from the sum of aforementioned MW of RBD and MWs of glycan residues from each possible N_H_F_S combination (Supplementary Table S5).

**Supplementary Table 1.** Composition of the defined library (composed of purified, free oligosaccharides) and the corresponding sub-libraries used for CaR-ESI-MS screening. Glycan codes, structures and molecular weights (MW) are indicated. The source of each glycan is given in the Materials and Methods (Supporting Information) section.

| **Glycan Code** | **Structure** | **Library** | **MW (Da)** |
|---|---|---|---|
| **1** | Galβ(1-3)[Fucα(1-4)]GlcNAc | B, G | 529.20 |
| **2** | Galβ(1-3)[Fucα(1-4)]GlcNAcβ(1-3)Gal | I | 691.25 |
| **3** | Galβ(1-3)[Fucα(1-4)]GlcNAcβ(1-3)Galβ(1-4)[Fucα(1-3)]Glc | C, L | 999.36 |
| **4** | GalNAcα(1-3)[Fucα(1-2)]Galβ(1-3)[Fucα(1-4)]GlcNAc | B | 878.39 |
| **5** | Galα(1-3)[Fucα(1-2)]Galβ(1-3)[Fucα(1-4)]GlcNAc | B | 837.31 |
| **6** | Fucα(1-2)Galβ(1-3)[Fucα(1-4)]GlcNAc | B, F | 675.26 |
| **7** | Fucα(1-2)Galβ(1-3)[Fucα(1-4)]GlcNAcβ(1-3)Gal | E | 837.31 |
| **8** | Galβ(1-4)[Fucα(1-3)]GlcNAc | D | 529.20 |
| **9** | Galβ(1-4)[Fucα(1-3)]GlcNAcβ(1-3)Gal | H | 691.25 |
| **10** | Galβ(1-4)[Fucα(1-3)]GlcNAcβ(1-3)Galβ(1-4)[Fucα(1-3)]Glc | B | 999.36 |
| **11** | GalNAcα(1-3)[Fucα(1-2)]Galβ(1-4)[Fucα(1-3)]GlcNAc | C | 878.39 |
| **12** | Galα(1-3)[Fucα(1-2)]Galβ(1-4)[Fucα(1-3)]GlcNAc | C | 837.31 |
| **13** | Fucα(1-2)Galβ(1-4)[Fucα(1-3)]GlcNAc | C, G | 675.26 |
| **14** | Fucα(1-2)Galβ(1-4)[Fucα(1-3)]GlcNAcβ(1-3)Gal | D | 837.31 |
| **15** | GalNAcα(1-3)[Fucα(1-2)]Galβ(1-3)GlcNAcβ1-O$(CH_2)_6CH=CH_2$ | F | 842.39 |
| **16** | GalNAcα(1-3)[Fucα(1-2)]Galβ(1-4)GlcNAcβ1-O$(CH_2)_6CH=CH_2$ | G | 842.39 |
| **17** | GalNAcα(1-3)[Fucα(1-2)]Galβ(1-3)GalNAcα1-O$(CH_2)_6CH=CH_2$ | H | 842.39 |
| **18** | GalNAcα(1-3)[Fucα(1-2)]Galβ(1-3)GalNAcβ1-O$(CH_2)_6CH=CH_2$ | I | 842.39 |
| **19** | GalNAcα(1-3)[Fucα(1-2)]Galβ(1-3)Galβ1-O$(CH_2)_6CH=CH_2$ | F | 801.36 |

| | | | |
|---|---|---|---|
| **20** | GalNAcα(1-3)[Fucα(1-2)]Galβ1-4Galβ1-O(CH$_2$)$_6$CH=CH$_2$ | G | 801.36 |
| **21** | Galα(1-3)[Fucα(1-2)]Galβ1-3GlcNAcβ1-O(CH$_2$)$_6$CH=CH$_2$ | H | 801.36 |
| **22** | Galα(1-3)(Fuca1-2)Galβ1-4GlcNAcβ1-O(CH$_2$)$_6$CH=CH$_2$ | I | 801.36 |
| **23** | Galα(1-3)(Fuca1-2)Galβ1-3GalNAcα1-O(CH$_2$)$_6$CH=CH$_2$ | J | 801.36 |
| **24** | Galα(1-3)[Fucα(1-2)]Galβ(1-3)GalNAcβ1-O(CH$_2$)$_6$CH=CH$_2$ | C | 801.36 |
| **25** | Galα(1-3)(Fuca1-2)Galβ1-3Galβ1-O(CH$_2$)$_6$CH=CH$_2$ | F | 760.33 |
| **26** | Galα(1-3)(Fuca1-2)Galβ1-4Galβ1-O(CH$_2$)$_6$CH=CH$_2$ | G | 760.33 |
| **27** | Fucα(1-2)Galβ1-3GlcNAcβ1-O(CH$_2$)$_6$CH=CH$_2$ | F | 639.31 |
| **28** | Fucα(1-2)Galβ1-4GlcNAcβ1-O(CH$_2$)$_6$CH=CH$_2$ | G | 639.31 |
| **29** | Fucα(1-2)Galβ1-3GalNAcα1-O(CH$_2$)$_6$CH=CH$_2$ | H | 639.31 |
| **30** | Fucα(1-2)Galβ1-3GalNAcβ1-O(CH$_2$)$_6$CH=CH$_2$ | I | 639.31 |
| **31** | Fucα(1-2)Galβ(1-3)Galβ1-O(CH$_2$)$_6$CH=CH$_2$ | F | 598.28 |
| **32** | Fucα(1-2)Galβ(1-4)Galβ1-O(CH$_2$)$_6$CH=CH$_2$ | G | 598.28 |
| **33** | ΔUAβ(1-3)GalNAc4,6S (diE) | A | 539.02 |
| **34** | ΔUA2Sβ(1-3)GalNAc4,6S (triS) | A | 618.98 |
| **35** | D-GlcAβ(1-4)GlcNAcα(1-4)GlcAβ(1-4)GlcNAc | A | 758.22 |
| **36** | D-GlcAβ(1-4)[Glc NAcα(1-4)GlcAβ(1-4)]$_4$GlcNAc | A | 1895.55 |
| **37** | GlcAβ(1-3)Galβ(1-4)Glc | A | 518.15 |
| **38** | GlcAβ(1-3)Galβ(1-3)GlcNAcβ(1-3)Galβ(1-4)Glc | A | 883.28 |
| **39** | IdoA2Sβ(1-4)GlcNS6Sα(CH$_2$)$_5$NH$_2$ | NA [a] | 680.07 |
| **40** | (IdoA2Sβ(1-4)GlcNS6S)$_2$α(CH$_2$)$_5$NH$_2$ | NA [a] | 1433.92 |
| **41** | (IdoA2Sβ(1-4)GlcNS6S)$_3$α(CH$_2$)$_5$NH$_2$ | NA [a] | 1834.01 |
| **42** | (IdoA2Sβ(1-4)GlcNS6S)$_4$α(CH$_2$)$_5$NH$_2$ | NA [a,b] | 2410.99 |
| **43** | GalNAcβ(1-3)GalNAcβ(1-3)Galα(1-4)Galβ(1-4)Glc | E | 910.33 |
| **44** | GalNAcβ(1-3)GalNAcβ(1-3)Gal | E | 586.22 |
| **45** | GalNAcα(1-3)GalNAcβ(1-3)Galα(1-3)Galβ(1-4)Glc | F | 910.33 |
| **46** | Galα(1-3)Galβ(1-4)GlcNAcβ(1-3)Galβ(1-4)Glc | C | 869.30 |
| **47** | Galα(1-3)[Galβ(1-4)GlcNAcβ(1-3)]$_2$Galβ(1-4)Glc | C | 1234.43 |
| **48** | Galα(1-3)[Galβ(1-4)GlcNAcβ(1-3)]$_3$Galβ(1-4)Glc | D | 1599.56 |

| | | | |
|---|---|---|---|
| **49** | Galα(1-3)[Galβ(1-4)GlcNAcβ(1-3)]$_4$Galβ1-4Glc | F | 1964.70 |
| **50** | Galα(1-3)Galβ(1-3)GlcNAc | D | 545.19 |
| **51** | Galα(1-3)Galβ(1-4)[Fucα(1-3)]GlcNAc | J | 691.25 |
| **52** | Galα(1-4)Galβ(1-4)Glc | D | 504.17 |
| **53** | GalNAcβ(1-3)Galα(1-3)Galβ(1-4)Glc | C | 707.25 |
| **54** | Galβ(1-3)GalNAcβ(1-3)Galα(1-3)Galβ(1-4)Glc | D | 869.30 |
| **55** | Galβ(1-3)GalNAcβ(1-3)Galα(1-4)Galβ(1-4)Glc | E | 869.30 |
| **56** | GalNAcβ(1-3)Galα(1-4)Galβ(1-4)Glc | E | 707.25 |
| **57** | Galα(1-4)Galβ(1-4)GlcNAc | G, J | 545.19 |
| **58** | Fucα(1-2)Galβ(1-3)GalNAcβ(1-3)Galα(1-4)Galβ(1-4)Glc | D | 1015.36 |
| **59** | Galα(1-3)[Fucα(1-2)]Galβ(1-3)GalNAcβ(1-3)Galα(1-4)Galβ(1-4)Glc | D | 1177.41 |
| **60** | GalNAcα(1-3)[Fucα(1-2)]Galβ(1-3)GalNAcβ(1-3)Galα(1-4)Galβ(1-4)Glc | D | 1218.44 |
| **61** | Neu5Acα(2-3)Galα(1-4)Galβ(1-4)Glc | E | 795.26 |
| **62** | Galβ(1-3)GalNAcβ(1-4)Galβ(1-4)Glc | D | 707.25 |
| **63** | GalNAcβ(1-4)Galβ(1-4)Glc | E | 545.19 |
| **64** | Galβ(1-4)[Neu5Acα(2-3)]Galβ1-4Glc) | D | 795.26 |
| **65** | Fucα(1-2)Galβ(1-3)GalNAcβ(1-4)[Neu5Acα(2-3)]Galβ1-4Glc | D | 1144.40 |
| **66** | Neu5Acα(2-3)Galβ(1-3)GalNAcβ(1-4)[Neu5Acα(2-3)]Galβ(1-4)Glc | D | 1289.44 |
| **67** | Galβ(1-3)GalNAcβ(1-4)[Neu5Acα(2-8)]Neu5Acα(2-3)Galβ(1-4)Glc | E | 1289.44 |
| **68** | GalNAcβ(1-4)[Neu5Acα(2-8)Neu5Acα(2-3)]Galβ(1-4)Glc | D | 1127.38 |
| **69** | Neu5Acα(2-8)Neu5Acα(2-3)Galβ(1-4)Glc | C | 924.30 |
| **70** | Galβ(1-3)GalNAcβ(1-4)[Neu5Acα(2-3)]Galβ(1-4)Glc | D | 998.34 |
| **71** | Neu5Acα(2-3)Galβ(1-3)GalNAcβ(1-4)Galβ(1-4)Glc | E | 998.34 |
| **72** | GalNAcβ(1-4)[Neu5Acα(2-3)]Galβ(1-4)Glc | I | 836.29 |
| **73** | Neu5Acα(2-3)Galβ(1-4)Glc | C, M | 633.21 |

| | | | |
|---|---|---|---|
| **74** | Neu5Acα(2-8)Neu5Acα(2-3)Galβ(1-3)GalNAcβ(1-4)[Neu5Acα(2-3)]Galβ(1-4)Glc | D | 1580.53 |
| **75** | Galβ(1-3)GalNAcβ(1-4)[Neu5Acα(2-8)Neu5Acα(2-8)Neu5Acα(2-3)]Galβ(1-4)Glc | E | 1580.53 |
| **76** | GalNAcβ(1-4)[Neu5Acα(2-8)Neu5Acα(2-8)Neu5Acα(2-3)]Galβ1-4Glc | D | 1418.48 |
| **77** | Neu5Acα(2-8)Neu5Acα(2-8)Neu5Acα(2-8)Galβ(1-4)Glc | D | 1215.40 |
| **78** | Galβ(1-3)Galβ(1-4)Glc | I | 545.19 |
| **79** | Galβ(1-4)GlcNAcβ(1-3)Galβ(1-4)GlcNAc | B | 748.27 |
| **80** | Galα(1-3)Galβ(1-4)Glc | B, C | 504.17 |
| **81** | Galβ(1-3)GlcNAcβ(1-3)Gal | B | 545.19 |
| **82** | Galβ(1-4)GlcNAcβ(1-3)Gal | C | 545.19 |
| **83** | Galβ(1-4)Glcβ1-1βGal | E | 504.17 |
| **84** | Galβ(1-3)Galβ(1-4)Galβ(1-3)Gal | E | 666.22 |
| **85** | Galβ(1-3)GalNAcβ(1-3)Gal | F | 545.19 |
| **86** | Manα(1-6)[Man α(1-3)]Manα(1-6)[Manα(1-6)]Man | F | 828.27 |
| **87** | Manα(1-6)[Man α(1-3)]Manα(1-6)Man | F | 666.22 |
| **88** | Galα(1-3)[Fucα(1-2)]Galβ(1-4)[Fucα(1-3)]Glc | B | 796.28 |
| **89** | Fucα(1-2)Galβ(1-4)[Fucα(1-3)]Glc | B, L | 634.23 |
| **90** | Galα(1-3)Galβ(1-4)[Fucα(1-3)]Glc | C | 650.22 |
| **91** | Fucα(1-2)Galβ(1-4)Glcβ(1-1)β[Fucα(1-2)]Gal | C | 796.28 |
| **92** | GlcNAcβ(1-4)[Fucα(1-6)]GlcNAc | E | 570.23 |
| **93** | Fucα(1-2)Galβ(1-4)[Fucα(1-2)]Glc | E | 634.23 |
| **94** | GalNAcα(1-3)[Fucα(1-2)]Galβ(1-4)[Fucα(1-3)]Glc | F | 837.31 |
| **95** | Neu5Acα(2-3)Galβ(1-4)[Fucα(1-3)]GlcNAcβ(1-3)Gal | B | 982.35 |
| **96** | Neu5Acα(2-3)Galβ(1-4)[Fucα(1-3)]Glc | B | 779.27 |
| **97** | Neu5Acα(2-3)Galβ(1-3)GalNAcβ(1-3)Gal | J | 836.29 |
| **98** | Neu5Acα(2-3)Galβ(1-4)Glcβ1-1β[Neu5Acα(2-3)]Gal | E | 1086.36 |
| **99** | Neu5Acα(2-8)Neu5Acα(2-3)Galβ(1-3)GlcNAcβ(1-3)Gal | E | 1127.38 |

| | | | |
|---|---|---|---|
| **100** | Neu5Acα(2-3)Galβ(1-3)GalNAcβ(1-3)Galα(1-4)Galβ(1-4)Glc | E | 1160.40 |
| **101** | Neu5Acα(2-3)Galβ(1-3)GlcNAcβ(1-3)Gal | G | 836.29 |
| **102** | Neu5Acα(2-3)Galβ(1-4)GlcNAcβ(1-3)Gal | H | 836.29 |
| **103** | Fucα(1-2)Galβ(1-4)Glc | K | 488.17 |
| **104** | GlcNAcβ(1-3)Galβ(1-4)Glc | K | 545.20 |
| **105** | GalNAcα(1-3)[Fucα(1-2)]Galβ(1-4)Glc | K | 691.25 |
| **106** | Galβ(1-3)GlcNAcβ(1-3)Galβ(1-4) Glc | K | 707.25 |
| **107** | Fucα(1-2)Galβ(1-3)GlcNAcβ(1-3)Galβ(1-4)Glc | K | 853.31 |
| **108** | Fucα(1-2)Galβ(1-3)[Fucα(1-4)]GlcNAcβ(1-3)Galβ(1-4)Glc | K | 999.36 |
| **109** | Galβ(1-4)GlcNAcβ(1-3)[Galβ(1-4)GlcNAcβ(1-3)]Galβ(1-4)Glc | K | 1072.38 |
| **110** | GalNAcα(1-3)[Fucα(1-2)]Galβ(1-3)[Fucα(1-4)]GlcNAcβ(1-3)Galβ(1-4)Glc | K | 1202.44 |
| **111** | Galβ(1-4)[Fucα(1-3)]GlcNAcβ(1-6)[Fucα(1-2)Galβ(1-3)GlcNAcβ(1-3)]Galβ(1-4)Glc | K | 1364.50 |
| **112** | Galβ(1-4)GlcNAcβ(1-3)Galβ(1-4)GlcNAcβ(1-3)Galβ(1-4)GlcNAcβ(1-3)Galβ(1-4)Glc | K | 1437.51 |
| **113** | Galβ(1-4)[Fucα(1-3)]Glc | L | 488.17 |
| **114** | Galβ(1-4)GlcNAcβ(1-3)Galβ(1-4)Glc | L | 707.25 |
| **115** | Galβ(1-3)[Fucα(1-4)]GlcNAcβ(1-3)Galβ(1-4)Glc | L | 853.31 |
| **116** | GalNAcα(1-3)[Fucα(1-2)]Galβ(1-3)GlcNAcβ(1-3)Galβ(1-4)Glc | L | 1056.39 |
| **117** | Galβ(1-4)GlcNAcβ(1-6)[Galβ(1-4)GlcNAcβ(1-3)]Galβ(1-4)Glc | L | 1072.38 |
| **118** | Galβ(1-4)[Fucα(1-3)]GlcNAcβ(1-6)[Galβ(1-3)GlcNAcβ(1-3)]Galβ(1-4)Glc | L | 1218.44 |
| **119** | Galβ(1-3)[Fucα(1-4)]GlcNAcβ(1-3)Galβ(1-4)[Fucα(1-3)]GlcNAcβ(1-3)Galβ(1-4)Glc | L | 1364.50 |
| **120** | Galβ(1-4)[Fucα(1-3)]GlcNAcβ(1-6)[Fucα(1-2)Galβ(1-3)[Fucα(1-4)]GlcNAcα(1-3)]Galα(1-4)Glc | L | 1510.55 |

| | | | |
|---|---|---|---|
| **121** | Galβ(1-4)[Fucα(1-3)]GlcNAcβ(1-3)Galβ(1-4)-βGlc | O | 853.31 |
| **122** | Galβ(1-4)[Fucα(1-3)]GlcNAcβ(1-3)Galβ(1-4)[Fucα(1-3)]Glc | O | 999.36 |
| **123** | Galα(1-4)GlcNAcβ(1-3)Galβ(1-4)GlcNAcβ(1-3)Galβ(1-4)Glc | O | 1072.38 |
| **124** | Galβ(1-3)GlcNAcα(1-3)Galβ(1-4)[Fucα(1-3)]Glc | O | 1144.40 |
| **125** | Neu5Acα(2-3)Galβ(1-4)GlcNAc | M | 674.24 |
| **126** | Neu5Acα(2-3)Galβ(1-3)GlcNAcβ(1-3)Galβ(1-4)Glc | M | 998.34 |
| **127** | Neu5Acα(2-3)Galβ(1-3)[Fucα(1-4)]GlcNAc | M | 820.30 |
| **128** | Neu5Ac(2-3)Galβ(1-3)[Fucα(1-4)]GlcNAcβ(1-3)Galβ(1-4)Glc | M | 1144.40 |
| **129** | Neu5Acα(2-3)Galβ(1-4)GlcNAcβ(1-3)Galβ(1-4)Glc | O | 998.34 |
| **130** | Neu5Acα(2-6)Galβ(1-4)Glc | N | 633.21 |
| **131** | Neu5Acα(2-6)Galβ(1-4)GlcNAc | N | 674.24 |
| **132** | Neu5Acα(2-6)[Galβ(1-3)]GlcNAcβ(1-3)Galβ(1-4)Glc | N | 998.34 |
| **133** | Fucα(1-2)Galβ(1-3)[Neu5Acα(2-6)]GlcNAcβ(1-3)Galβ(1-4)Glc | N | 1144.40 |
| **134** | Rhaα(1-3)2MeRhaPMP | P[c,d] | 432.47 |
| **135** | Rhaα(1-3)GlcNAcOc | P[e] | 481.46 |
| **136** | 2,4MeFucα(1-3)Rhaα(1-3)RhaPMP | P[c,d,f] | 605.20 |
| **137** | 2,3,4MeFucα(1-3)Rhaα(1-3)RhaPMP | P[c,d,f] | 619.22 |
| **138** | 3,6MeGlcβ(1-4)Rhaα(1-2)RhaPMP | P[c,d] | 621.22 |
| **139** | 6MeGlcβ(1-4)2,3MeRhaα(1-2)3MeRhaPMP | P[c,d] | 649.38 |
| **140** | Galβ(1-5)Galβ(1-4)Rhaα(1-3)GlcNAcOc | P[e] | 804.80 |
| **141** | Neu5Acα(2-3)Galβ(1-4)GlcNAcβ(1-2)Manα(1-3)][Manα(1-6)]Manβ(1-4)GlcNAcβ(1-4)GlcNAc | NA[a] | 1566.56 |

a. NA ≡ not applicable; binding of these glycans to RBD was evaluated by direct ESI-MS analysis.

b. The relative abundance of **41** in the **42** sample was estimated to be 94% by ESI-MS.

c. PMP ≡ *p*-methoxyphenyl. d. Me ≡ methyl. e. Oc ≡ Octyl. f. Bn ≡ benzyl.

**Supplementary Table 2.** Summary of molecular weights (MWs) of RBD glycoforms ($P_x$) measured by ESI-MS performed in positive ion mode on a Q Exactive Orbitrap mass spectrometer.

| $P_x$ | Measured mass-to-charge ratios (*m/z*) | | | Measured MW (Da) |
|---|---|---|---|---|
| | z = 9 | z = 10 | z = 11 | |
| P1 | 3447.5 | 3102.8 | 2821.0 | 31018 ± 1 |
| P2 | 3451.8 | 3106.5 | 2824.2 | 31056 ± 2 |
| P3 | 3463.6 | 3117.4 | 2834.1 | 31164 ± 1 |
| P4 | 3467.1 | 3120.8 | 2837.2 | 31202 ± 2 |
| P5 | 3475.4 | 3127.8 | 2843.7 | 31268 ± 1 |
| P6 | 3479.6 | 3132.0 | 2847.4 | 31310 ± 2 |
| P7 | 3483.6 | 3135.3 | 2850.4 | 31343 ± 1 |
| P8 | 3486.0 | 3137.2 | 2852.3 | 31364 ± 2 |
| P9 | 3487.9 | 3139.3 | 2854.2 | 31384 ± 2 |
| P10 | 3491.7 | 3142.6 | 2857.1 | 31416 ± 1 |
| P11 | 3496.0 | 3146.7 | 2860.8 | 31456 ± 2 |
| P12 | 3497.9 | 3148.2 | 2862.1 | 31472 ± 1 |
| P13 | 3499.8 | 3150.0 | 2863.7 | 31490 ± 1 |
| P14 | 3501.5 | 3151.6 | 2865.4 | 31506 ± 2 |
| P15 | 3504.2 | 3154.0 | 2867.3 | 31530 ± 1 |
| P16 | 3506.2 | 3155.6 | 2868.9 | 31547 ± 1 |
| P17 | 3507.9 | 3157.2 | 2870.3 | 31562 ± 1 |
| P18 | 3509.8 | 3158.9 | 2872.0 | 31578 ± 2 |

| | | | | |
|---|---|---|---|---|
| P19 | 3512.5 | 3161.2 | 2873.8 | 31602 ± 2 |
| P20 | 3512.6 | 3161.2 | 2873.8 | 31603 ± 2 |
| P21 | 3515.9 | 3164.5 | 2876.9 | 31635 ± 1 |
| P22 | 3518.0 | 3166.3 | 2878.6 | 31653 ± 1 |
| P23 | 3520.3 | 3168.5 | 2880.5 | 31674 ± 1 |
| P24 | 3522.1 | 3170.1 | 2882.0 | 31691 ± 2 |
| P25 | 3524.0 | 3171.9 | 2883.6 | 31709 ± 1 |
| P26 | 3528.5 | 3175.8 | 2887.3 | 31749 ± 1 |
| P27 | 3532.2 | 3179.1 | 2890.2 | 31781 ± 1 |
| P28 | 3536.6 | 3183.2 | 2893.7 | 31821 ± 2 |
| P29 | 3538.4 | 3184.8 | 2895.3 | 31837 ± 1 |
| P30 | 3540.2 | 3186.4 | 2896.9 | 31854 ± 1 |
| P31 | 3542.0 | 3188.1 | 2898.5 | 31871 ± 2 |
| P32 | 3545.1 | 3190.4 | 2900.5 | 31895 ± 2 |
| P33 | 3548.3 | 3193.6 | 2903.4 | 31926 ± 1 |
| P34 | 3550.1 | 3195.6 | 2904.9 | 31944 ± 2 |
| P35 | 3553.2 | 3197.6 | 2907.1 | 31968 ± 2 |
| P36 | 3554.6 | 3199.3 | 2908.5 | 31983 ± 1 |
| P37 | 3556.6 | 3201.0 | 2910.1 | 32000 ± 1 |
| P38 | 3558.4 | 3202.7 | 2911.6 | 32018 ± 1 |
| P39 | 3561.0 | 3205.0 | 2913.6 | 32040 ± 1 |
| P40 | 3564.7 | 3208.3 | 2916.8 | 32073 ± 1 |
| P41 | 3569.4 | 3212.3 | 2920.4 | 32114 ± 2 |

| | | | | |
|---|---|---|---|---|
| P42 | 3570.7 | 3213.9 | 2921.8 | 32128 ± 1 |
| P43 | 3572.9 | 3215.6 | 2923.4 | 32147 ± 1 |
| P44 | 3577.2 | 3219.7 | 2927.0 | 32186 ± 1 |
| P45 | 3580.8 | 3222.8 | 2929.9 | 32218 ± 1 |
| P46 | 3582.7 | 3224.5 | 2931.6 | 32236 ± 1 |
| P47 | 3589.0 | 3230.2 | 2936.6 | 32291 ± 1 |
| P48 | 3590.7 | 3232.0 | 2938.3 | 32309 ± 2 |
| P49 | 3593.4 | 3233.9 | 2940.2 | 32330 ± 2 |
| P50 | 3597.1 | 3237.5 | 2943.3 | 32365 ± 1 |
| P51 | 3601.5 | 3241.3 | 2946.8 | 32405 ± 1 |
| P52 | 3605.3 | 3244.8 | 2949.9 | 32438 ± 1 |
| P53 | 3609.6 | 3248.6 | 2953.2 | 32476 ± 2 |
| P54 | 3611.5 | 3250.4 | 2955.1 | 32495 ± 1 |
| P55 | 3613.3 | 3252.1 | 2956.6 | 32511 ± 1 |
| P56 | 3621.3 | 3259.3 | 2963.0 | 32583 ± 1 |
| P57 | 3625.9 | 3263.2 | 2966.7 | 32623 ± 1 |
| P58 | 3629.4 | 3266.7 | 2969.8 | 32656 ± 1 |
| P59 | 3634.0 | 3270.8 | 2973.4 | 32697 ± 1 |
| P60 | 3637.5 | 3274.0 | 2976.4 | 32729 ± 1 |
| P61 | 3645.8 | 3281.3 | 2983.2 | 32803 ± 1 |
| P62 | 3649.9 | 3285.1 | 2986.7 | 32841 ± 2 |
| P63 | 3653.8 | 3288.5 | 2989.5 | 32874 ± 2 |
| P64 | 3658.1 | 3292.6 | 2993.1 | 32914 ± 2 |

| | | | | |
|---|---|---|---|---|
| P65 | 3661.7 | 3295.8 | 2996.2 | 32947 ± 1 |
| P66 | 3665.7 | 3299.5 | 2999.4 | 32988 ± 2 |
| P67 | 3670.1 | 3303.2 | 3003.1 | 33023 ± 2 |
| P68 | 3675.0 | 3307.0 | 3006.6 | 33062 ± 2 |
| P69 | 3678.1 | 3310.5 | 3009.6 | 33094 ± 2 |
| P70 | 3686.2 | 3317.8 | 3016.3 | 33167 ± 2 |
| P71 | 3694.3 | 3325.0 | 3022.7 | 33239 ± 2 |
| P72 | 3702.5 | 3332.3 | 3029.5 | 33313 ± 2 |
| P73 | 3707.5 | 3335.8 | 3032.8 | 33353 ± 2 |
| P74 | 3726.7 | 3354.1 | 3049.1 | 33530 ± 2 |
| P75 | 3734.7 | 3361.5 | 3056.2 | 33605 ± 2 |
| P76 | 3766.1 | 3390.6 | 3082.4 | 33894 ± 2 |
| P77 | 3798.7 | 3420.0 | 3108.9 | 34186 ± 2 |

a. Listed MWs are averages of the values calculated for charge states +9, +10, and +11.

b. Uncertainties correspond to one standard deviation.

**Supplementary Table 3.** Summary of reported RBD *O*-glycosylation (at T323 and S325).

| *O*-glycan composition | Shajahan [a] | Zhang [b] | Sanda [c] | Zhao [d] |
|---|---|---|---|---|
| $HexNAc_1$ | T323 | T323/S325 | N/A | T323/S325 |
| $HexNAc_1Hex_1$ | T323 | T323/S325 | T323/S325 | T323 |
| $HexNAc_1Hex_1Neu5Ac_1$ | T323/S325 | T323/S325 | T323/S325 | T323 |
| $HexNAc_1Hex_1Neu5Ac_2$ | T323 | T323/S325 | T323/S325 | T323 |
| $HexNAc_2$ | N/A | T323/S325 | N/A | N/A |
| $HexNAc_2Hex_1$ | N/A | T323 | N/A | T323 |
| $HexNAc_2Hex_2$ | N/A | N/A | N/A | T323 |
| $HexNAc_2Hex_2Neu5Ac_1$ | T323 | N/A | T323/S325 | N/A |
| $HexNAc_2Hex_2Neu5Ac_2$ | T323 | N/A | N/A | N/A |

a. Reference S25.

b. Reference S26.

c. Reference S27.

d. Reference S28.

**Supplementary Table 4.** Summary of glycans identified HILIC-UHPLC analysis of 2-AB labeled *N*-glycans released from RBD.[a,b]

| No. | Glycan composition | Putative structure | Number of isomers | Relative abundance (%) |
|---|---|---|---|---|
| 1 | $\mathrm{HexNAc_4Hex_5Fuc_1Neu5Ac_1}$ | 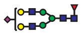 | 6 | 100 |
| 2 | $\mathrm{HexNAc_4Hex_5Fuc_1Neu5Ac_2}$ | 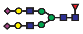 | 3 | 57.2 |
| 3 | $\mathrm{HexNAc_4Hex_5Fuc_2Neu5Ac_1}$ | 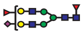 | 5 | 43.2 |
| 4 | $\mathrm{HexNAc_4Hex_4Fuc_1Neu5Ac_1}$ | 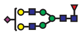 | 4 | 38.7 |
| 5 | $\mathrm{HexNAc_4Hex_5Fuc_2}$ | 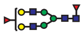 | 3 | 30.5 |
| 6 | $\mathrm{HexNAc_4Hex_5Fuc_1}$ | 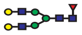 | 1 | 28.3 |
| 7 | $\mathrm{HexNAc_5Hex_4Fuc_1Neu5Ac_1}$ | 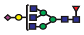 | 6 | 26 |
| 8 | $\mathrm{HexNAc_5Hex_3Fuc_1}$ | 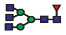 | 3 | 25.4 |
| 9 | $\mathrm{HexNAc_5Hex_4Fuc_2}$ | 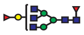 | 2 | 24.4 |
| 10 | $\mathrm{HexNAc_4Hex_3Fuc_1}$ | 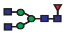 | 4 | 23.4 |
| 11 | $\mathrm{HexNAc_4Hex_4Fuc_2}$ | 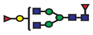 | 3 | 21.9 |
| 12 | $\mathrm{HexNAc_2Hex_5}$ | 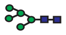 | 1 | 20.2 |
| 13 | $\mathrm{HexNAc_5Hex_4Fuc_1}$ | 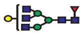 | 6 | 18.9 |
| 14 | $\mathrm{HexNAc_5Hex_6Fuc_1Neu5Ac_1}$ | 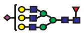 | 5 | 17.9 |
| 15 | $\mathrm{HexNAc_6Hex_3Fuc_2}$ | 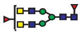 | 2 | 17.1 |
| 16 | $\mathrm{HexNAc_5Hex_6Fuc_1Neu5Ac_2}$ | 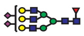 | 4 | 14.8 |
| 17 | $\mathrm{HexNAc_5Hex_6Fuc_1}$ | 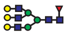 | 1 | 14.7 |
| 18 | $\mathrm{HexNAc_4Hex_3Fuc_2}$ | 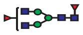 | 1 | 13.0 |
| 19 | $\mathrm{HexNAc_5Hex_4Fuc_2Neu5Ac_1}$ | 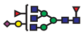 | 2 | 12.4 |
| 20 | $\mathrm{HexNAc_4Hex_4Fuc_1}$ | 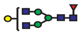 | 3 | 12.3 |
| 21 | $\mathrm{HexNAc_5Hex_3Fuc_2}$ | 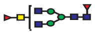 | 2 | 9.6 |
| 22 | $\mathrm{HexNAc_6Hex_3Fuc_1}$ | 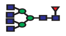 | 3 | 9.2 |
| 23 | $\mathrm{HexNAc_6Hex_3Fuc_1Neu5Ac_1}$ | 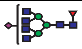 | 1 | 8.1 |
| 24 | $\mathrm{HexNAc_5Hex_3Fuc_1Neu5Ac_1}$ | 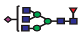 | 1 | 7.5 |

| | | | | |
|---|---|---|---|---|
| 25 | $HexNAc_6Hex_3Fuc_2Neu5Ac_1$ | 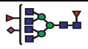 | 1 | 7.3 |
| 26 | $HexNAc_5Hex_6Fuc_2Neu5Ac_1$ | 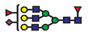 | 4 | 5.2 |
| 27 | $HexNAc_5Hex_4Fuc_1S_1$ | 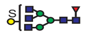 | 1 | 5.1 |
| 28 | $HexNAc_5Hex_6Fuc_1Neu5Ac_3$ | 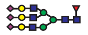 | 2 | 4.9 |
| 29 | $HexNAc_3Hex_3Fuc_1$ | 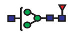 | 2 | 4.8 |
| 30 | $HexNAc_5Hex_6Fuc_2$ | 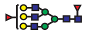 | 1 | 4.8 |
| 31 | $HexNAc_6Hex_4Fuc_1Neu5Ac_1$ | 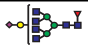 | 1 | 4.4 |
| 32 | $HexNAc_4Hex_5Fuc_4$ | 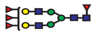 | 1 | 3.8 |
| 33 | $HexNAc_5Hex_4Fuc_1Neu5Ac_1S_1$ | 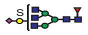 | 2 | 3.7 |
| 34 | $HexNAc_6Hex_5Fuc_1Neu5Ac_2$ | 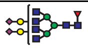 | 1 | 3.3 |
| 35 | $HexNAc_3Hex_4Fuc_1Neu5Ac_1$ | 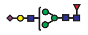 | 2 | 3.2 |
| 36 | $HexNAc_5Hex_4Fuc_3$ | 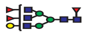 | 1 | 2.9 |
| 37 | $HexNAc_6Hex_5Fuc_2$ | 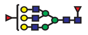 | 1 | 2.7 |
| 38 | $HexNAc_4Hex_5Neu5Ac_1$ | 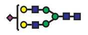 | 2 | 2.3 |
| 39 | $HexNAc_5Hex_3Fuc_1S_1$ | 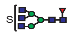 | 2 | 2.3 |
| 40 | $HexNAc_6Hex_3Fuc_1S_1$ | 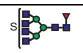 | 1 | 2.3 |
| 41 | $HexNAc_4Hex_5Fuc_3Neu5Ac_1$ | 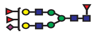 | 2 | 2.0 |
| 42 | $HexNAc_3Hex_5Fuc_1Neu5Ac_1$ | 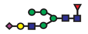 | 1 | 1.6 |
| 43 | $HexNAc_6Hex_3Fuc_1S_2$ | 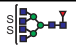 | 1 | 1.6 |
| 44 | $HexNAc_4Hex_5Fuc_2Neu5Ac_2$ | 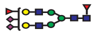 | 2 | 1.5 |
| 45 | $HexNAc_3Hex_4Fuc_1$ | 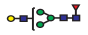 | 2 | 1.4 |
| 46 | $HexNAc_5Hex_5Fuc_1Neu5Ac_1$ | 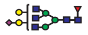 | 1 | 1.4 |
| 47 | $HexNAc_6Hex_5Fuc_1Neu5Ac_1$ | 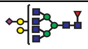 | 1 | 1.4 |
| 48 | $HexNAc_4Hex_3Fuc_1Neu5Ac_1$ | 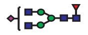 | 1 | 1.2 |
| 49 | $HexNAc_5Hex_6Fuc_2Neu5Ac_2$ | 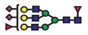 | 2 | 1.1 |
| 50 | $HexNAc_2Hex_3$ | 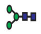 | 1 | 1.0 |
| 51 | $HexNAc_4Hex_5Fuc_4Neu5Ac_1$ | 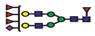 | 1 | 1.0 |
| 52 | $HexNAc_5Hex_6Fuc_3$ | 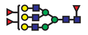 | 1 | 0.9 |
| 53 | $HexNAc_3Hex_3$ | 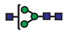 | 1 | 0.8 |

| | | | | |
|---|---|---|---|---|
| 54 | $HexNAc_6Hex_7Fuc_1Neu5Ac_2$ | 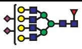 | 1 | 0.7 |
| 55 | $HexNAc_5Hex_6Fuc_3Neu5Ac_1$ | 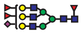 | 1 | 0.5 |
| 56 | $HexNAc_6Hex_7Fuc_1Neu5Ac_1$ | 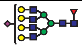 | 1 | 0.5 |
| 57 | $HexNAc_7Hex_5$ | 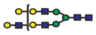 | 1 | 0.3 |
| 58 | $HexNAc_2Hex_1Neu5Ac_1$ | | 1 | 0.2 |
| 59 | $HexNAc_2Hex_3Fuc_1$ | 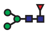 | 1 | 0.1 |
| 60 | $HexNAc_3Hex_4$ | 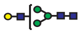 | 2 | 0.1 |
| 61 | $HexNAc_3Hex_4Fuc_2$ | 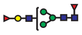 | 1 | 0.1 |
| 62 | $HexNAc_2Hex_4$ | 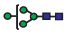 | 1 | 0 |
| 63 | $HexNAc_2Hex_4Fuc_1$ | 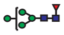 | 1 | 0 |
| 64 | $HexNAc_2Hex_6$ | 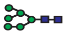 | 1 | 0 |
| 65 | $HexNAc_3Hex_3Fuc_2$ | 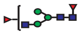 | 1 | 0 |
| 66 | $HexNAc_3Hex_5Fuc_1$ | 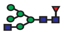 | 1 | 0 |
| 67 | $HexNAc_3Hex_5Fuc_2$ | 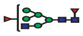 | 1 | 0 |
| 68 | $HexNAc_3Hex_6Fuc_1$ | 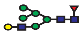 | 1 | 0 |
| 69 | $HexNAc_3Hex_6P_1$ | 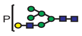 | 1 | 0 |
| 70 | $HexNAc_4Hex_3$ | 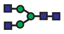 | 1 | 0 |
| 71 | $HexNAc_4Hex_3Fuc_1S_1$ | 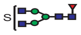 | 1 | 0 |
| 72 | $HexNAc_4Hex_5$ | 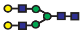 | 1 | 0 |
| 73 | $HexNAc_4Hex_6Fuc_1$ | 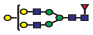 | 1 | 0 |
| 74 | $HexNAc_6Hex_7Fuc_1Neu5Ac_3$ | 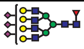 | 1 | 0 |

a. S indicates sulfate group, P indicates phosphate group.

b. *N*-glycans may have either core or antennary fucosylation.

**Supplementary Table 5.** Putative glycan compositions and relative abundances of the RBD glycoforms ($P_x$) identified by ESI-MS.

<table>
<tr><th rowspan="2">$P_x$</th><th rowspan="2">Measured MW (Da)</th><th rowspan="2">Relative abundance[a] (%)</th><th rowspan="2">Putative glycan composition N_H_F_S</th><th rowspan="2">Calculated MW (Da)</th><th colspan="4">Possible glycan combinations</th></tr>
<tr><th colspan="2">O-glycans</th><th colspan="2">N-glycans</th></tr>
<tr><td rowspan="2">P1</td><td rowspan="2">31018</td><td rowspan="2">1</td><td rowspan="2">10_10_2_4</td><td rowspan="2">31020</td><td colspan="2">2202</td><td rowspan="2">4411</td><td rowspan="2">4411</td></tr>
<tr><td>1101</td><td>1101</td></tr>
<tr><td rowspan="2">P2</td><td rowspan="2">31056</td><td rowspan="2">1</td><td rowspan="2">10_12_4_2</td><td rowspan="2">31054</td><td colspan="2">2201</td><td>4520</td><td>4521</td></tr>
<tr><td>1100</td><td>1100</td><td>4521</td><td>4521</td></tr>
<tr><td rowspan="2">P3</td><td rowspan="2">31164</td><td rowspan="2">1</td><td rowspan="2">10_10_3_4</td><td rowspan="2">31166</td><td colspan="2">1102</td><td>5420</td><td>4512</td></tr>
<tr><td>1101</td><td>1102</td><td>4411</td><td>4420</td></tr>
<tr><td rowspan="2">P4</td><td rowspan="2">31202</td><td rowspan="2">1</td><td rowspan="2">12_14_2_1</td><td rowspan="2">31201</td><td colspan="2">2201</td><td>5610</td><td>5610</td></tr>
<tr><td>1100</td><td>1100</td><td>5610</td><td>5611</td></tr>
<tr><td rowspan="2">P5</td><td rowspan="2">31268</td><td rowspan="2">5</td><td rowspan="2">9_11_2_5</td><td rowspan="2">31270</td><td colspan="2">1102</td><td>4511</td><td>4512</td></tr>
<tr><td>1100</td><td>1102</td><td>3511</td><td>4512</td></tr>
<tr><td rowspan="2">P6</td><td rowspan="2">31310</td><td rowspan="2">11</td><td rowspan="2">10_10_2_5</td><td rowspan="2">31311</td><td colspan="2">1100</td><td>4512</td><td>5411</td></tr>
<tr><td>1101</td><td>1101</td><td>4311</td><td>4512</td></tr>
<tr><td rowspan="2">P7</td><td rowspan="2">31343</td><td rowspan="2">7</td><td rowspan="2">10_12_2_4</td><td rowspan="2">31344</td><td colspan="2">2201</td><td rowspan="2">4511</td><td rowspan="2">4512</td></tr>
<tr><td>1101</td><td>1101</td></tr>
<tr><td>P8</td><td>31364</td><td>1</td><td>12_15_2_1</td><td>31363</td><td>1100</td><td>2201</td><td>4610</td><td>5610</td></tr>
<tr><td rowspan="2">P9</td><td rowspan="2">31384</td><td rowspan="2">20</td><td rowspan="2">11_11_2_4</td><td rowspan="2">31385</td><td colspan="2">2201</td><td rowspan="2">4512</td><td rowspan="2">5411</td></tr>
<tr><td>1101</td><td>1101</td></tr>
<tr><td rowspan="2">P10</td><td rowspan="2">31416</td><td rowspan="2">26</td><td rowspan="2">11_13_2_3</td><td rowspan="2">31418</td><td colspan="2">2201</td><td rowspan="2">4511</td><td rowspan="2">5411</td></tr>
<tr><td>1100</td><td>1101</td></tr>
<tr><td>P11</td><td>31456</td><td>2</td><td>10_10_3_5</td><td>31457</td><td>1102</td><td>1102</td><td>4310</td><td>4512</td></tr>
<tr><td rowspan="2">P12</td><td rowspan="2">31472</td><td rowspan="2">2</td><td rowspan="2">10_11_2_5</td><td rowspan="2">31473</td><td colspan="2">2202</td><td rowspan="2">4411</td><td rowspan="2">4512</td></tr>
<tr><td>1101</td><td>1101</td></tr>
<tr><td>P13</td><td>31490</td><td>26</td><td>12_14_2_2</td><td>31492</td><td colspan="2">2202</td><td>5610</td><td>5610</td></tr>
</table>

<table>
<tr><td></td><td></td><td></td><td></td><td></td><td>2201</td><td>2201</td><td>4510</td><td>4510</td></tr>
<tr><td rowspan="2">P14</td><td rowspan="2">31506</td><td rowspan="2">4</td><td rowspan="2">10_13_2_4</td><td rowspan="2">31506</td><td colspan="2">1101</td><td>4511</td><td>6712</td></tr>
<tr><td>2201</td><td>2201</td><td>4511</td><td>4512</td></tr>
<tr><td rowspan="2">P15</td><td rowspan="2">31530</td><td rowspan="2">3</td><td rowspan="2">11_11_3_4</td><td rowspan="2">31531</td><td colspan="2">2202</td><td rowspan="2">4521</td><td rowspan="2">5411</td></tr>
<tr><td>1101</td><td>1101</td></tr>
<tr><td rowspan="2">P16</td><td rowspan="2">31547</td><td rowspan="2">2</td><td rowspan="2">11_12_2_4</td><td rowspan="2">31547</td><td colspan="2">2202</td><td rowspan="2">4411</td><td rowspan="2">5611</td></tr>
<tr><td>1101</td><td>1101</td></tr>
<tr><td rowspan="2">P17</td><td rowspan="2">31562</td><td rowspan="2">7</td><td rowspan="2">11_13_3_3</td><td rowspan="2">31564</td><td colspan="2">2202</td><td rowspan="2">4521</td><td rowspan="2">5610</td></tr>
<tr><td>1101</td><td>1101</td></tr>
<tr><td rowspan="2">P18</td><td rowspan="2">31578</td><td rowspan="2">2</td><td rowspan="2">11_14_2_3</td><td rowspan="2">31580</td><td colspan="2">2201</td><td>4610</td><td>5611</td></tr>
<tr><td>1101</td><td>2201</td><td>3511</td><td>5610</td></tr>
<tr><td rowspan="2">P19</td><td rowspan="2">31602</td><td rowspan="2">4</td><td rowspan="2">10_10_4_5</td><td rowspan="2">31603</td><td colspan="2">2202</td><td>4531</td><td>4311</td></tr>
<tr><td>1102</td><td>1102</td><td>4411</td><td>4411</td></tr>
<tr><td rowspan="2">P20</td><td rowspan="2">31603</td><td rowspan="2">8</td><td rowspan="2">12_12_3_3</td><td rowspan="2">31605</td><td colspan="2">2202</td><td rowspan="2">4521</td><td rowspan="2">6512</td></tr>
<tr><td>1101</td><td>1101</td></tr>
<tr><td rowspan="2">P21</td><td rowspan="2">31635</td><td rowspan="2">44</td><td rowspan="2">10_12_2_5</td><td rowspan="2">31635</td><td colspan="2">2202</td><td rowspan="2">4511</td><td rowspan="2">4512</td></tr>
<tr><td>1101</td><td>1101</td></tr>
<tr><td rowspan="2">P22</td><td rowspan="2">31653</td><td rowspan="2">3</td><td rowspan="2">10_13_3_4</td><td rowspan="2">31652</td><td colspan="2">2202</td><td rowspan="2">3511</td><td rowspan="2">5621</td></tr>
<tr><td>1101</td><td>1101</td></tr>
<tr><td rowspan="2">P23</td><td rowspan="2">31674</td><td rowspan="2">5</td><td rowspan="2">11_11_2_5</td><td rowspan="2">31676</td><td colspan="2">2202</td><td rowspan="2">4512</td><td rowspan="2">5411</td></tr>
<tr><td>1101</td><td>1101</td></tr>
<tr><td rowspan="2">P24</td><td rowspan="2">31691</td><td rowspan="2">1</td><td rowspan="2">11_12_3_4</td><td rowspan="2">31693</td><td colspan="2">2202</td><td rowspan="2">4521</td><td rowspan="2">5511</td></tr>
<tr><td>1101</td><td>1101</td></tr>
<tr><td rowspan="2">P25</td><td rowspan="2">31709</td><td rowspan="2">8</td><td rowspan="2">11_13_2_4</td><td rowspan="2">31709</td><td colspan="2">2202</td><td rowspan="2">4511</td><td rowspan="2">5611</td></tr>
<tr><td>1101</td><td>1101</td></tr>
<tr><td rowspan="2">P26</td><td rowspan="2">31749</td><td rowspan="2">22</td><td rowspan="2">12_12_2_4</td><td rowspan="2">31750</td><td colspan="2">2202</td><td>5511</td><td>5511</td></tr>
<tr><td>2201</td><td>2201</td><td>4411</td><td>4411</td></tr>
<tr><td rowspan="2">P27</td><td rowspan="2">31781</td><td rowspan="2">65</td><td rowspan="2">10_12_3_5</td><td rowspan="2">31781</td><td colspan="2">2202</td><td rowspan="2">4512</td><td rowspan="2">4521</td></tr>
<tr><td>1101</td><td>1102</td></tr>
<tr><td>P28</td><td>31821</td><td>30</td><td>11_11_3_5</td><td>31822</td><td colspan="2">2202</td><td>4512</td><td>5421</td></tr>
</table>

| | | | | | | | | |
|---|---|---|---|---|---|---|---|---|
| | | | | | 1102 | 2202 | 4311 | 4520 |
| P29 | 31837 | 6 | 11_12_2_5 | 31838 | 2202 | | 4512 | 5511 |
| | | | | | 1101 | 1101 | | |
| P30 | 31854 | 9 | 11_13_3_4 | 31855 | 2202 | | 4521 | 5611 |
| | | | | | 1101 | 1101 | | |
| P31 | 31871 | 2 | 11_14_2_4 | 31871 | 2202 | | 4511 | 5511 |
| | | | | | 1101 | 1101 | | |
| P32 | 31895 | 14 | 12_12_3_4 | 31896 | 2202 | | 4521 | 6511 |
| | | | | | 1101 | 2201 | 4521 | 5411 |
| P33 | 31926 | 100 | 10_12_2_6 | 31926 | 2202 | | 4512 | 4512 |
| | | | | | 1102 | 1102 | 4511 | 4511 |
| P34 | 31944 | 1 | 12_15_2_3 | 31945 | 2201 | 2201 | 3511 | 5611 |
| P35 | 31968 | 37 | 11_11_2_6 | 31967 | 1102 | | 4512 | 6512 |
| | | | | | 1102 | 2202 | 4310 | 4512 |
| P36 | 31983 | 2 | 11_12_3_5 | 31984 | 1101 | 1102 | 4521 | 5511 |
| P37 | 32000 | 49 | 11_13_2_5 | 32000 | 2202 | | 4511 | 5612 |
| | | | | | 1102 | 2201 | 4511 | 4511 |
| P38 | 32018 | 2 | 11_14_3_4 | 32018 | 2201 | 2202 | 3511 | 4520 |
| P39 | 32040 | 30 | 12_12_2_5 | 32041 | 2202 | | 5411 | 5612 |
| | | | | | 1101 | 2201 | 4511 | 5411 |
| P40 | 32073 | 83 | 12_14_2_5 | 32074 | 2202 | | 5611 | 5611 |
| | | | | | 2201 | 2201 | 4511 | 4511 |
| P41 | 32114 | 27 | 13_13_2_4 | 32115 | 2202 | | 5611 | 6511 |
| | | | | | 2201 | 2201 | 4511 | 5411 |
| P42 | 32128 | 4 | 11_12_2_6 | 32129 | 2202 | | 4411 | 5613 |
| | | | | | 1101 | 1101 | | |
| P43 | 32147 | 42 | 11_13_3_5 | 32146 | 2202 | | 4521 | 5612 |

| | | | | | | | | |
|---|---|---|---|---|---|---|---|---|
| | | | | | 1101 | 2202 | 4511 | 4521 |
| P44 | 32186 | 12 | 12_12_3_5 | 322187 | 2202 | | 5420 | 5613 |
| | | | | | 2201 | 2202 | 4521 | 5411 |
| P45 | 32218 | 33 | 10_12_2_7 | 32217 | 1102 | | 4512 | 5613 |
| | | | | | 1102 | 1102 | 4511 | 4512 |
| P46 | 32236 | 26 | 12_15_4_3 | 32237 | 1101 | 2202 | 5621 | 5630 |
| P47 | 32291 | 61 | 11_13_2_6 | 32291 | 2202 | | 4512 | 5612 |
| | | | | | 1102 | 2201 | 4511 | 4512 |
| P48 | 32309 | 3 | 11_14_3_5 | 32308 | 1101 | 2202 | 4521 | 3511 |
| P49 | 32330 | 23 | 12_12_2_6 | 32332 | 2202 | | 5411 | 5613 |
| | | | | | 2202 | 2202 | 4511 | 5411 |
| P50 | 32365 | 74 | 12_14_2_5 | 32365 | 2202 | | 5611 | 5612 |
| | | | | | 2201 | 2201 | 4511 | 4512 |
| P51 | 32405 | 11 | 13_13_2_5 | 32406 | 2202 | | 5611 | 6512 |
| | | | | | 1101 | 2201 | 4511 | 5412 |
| P52 | 32438 | 45 | 11_13_3_6 | 32437 | 2202 | | 4521 | 5613 |
| | | | | | 1102 | 2202 | 4521 | 4511 |
| P53 | 32476 | 2 | 12_12_3_6 | 32478 | 1102 | 2202 | 4512 | 5420 |
| P54 | 32495 | 4 | 12_13_2_6 | 32494 | 2202 | | 5511 | 5613 |
| | | | | | 2201 | 2202 | 4411 | 4512 |
| P55 | 32511 | 37 | 12_14_3_5 | 32511 | 2202 | | 5613 | 5522 |
| | | | | | 2201 | 2202 | 4511 | 4521 |
| P56 | 32583 | 68 | 11_13_2_7 | 32582 | 2202 | | 4512 | 5613 |
| | | | | | 2202 | 2202 | 4511 | 4512 |
| P57 | 32623 | 9 | 12_12_4_6 | 32624 | 2202 | | 5613 | 5631 |
| | | | | | 1101 | 2202 | 4512 | 4531 |
| P58 | 32656 | 56 | 12_14_2_6 | 32656 | 2202 | | 5612 | 5612 |
| | | | | | 2202 | 2202 | 4511 | 4511 |
| P59 | 32697 | 11 | 13_13_2_6 | 32697 | 2202 | | 5612 | 6512 |
| | | | | | 1101 | 2202 | 4511 | 6512 |

<table>
<tr><td>P60</td><td>32729</td><td>51</td><td>13_15_2_5</td><td>32730</td><td>2201</td><td>2202</td><td>4511</td><td>5611</td></tr>
<tr><td>P61</td><td>32803</td><td>18</td><td>12_14_3_6</td><td>32802</td><td>2202</td><td>2202</td><td>4511</td><td>4521</td></tr>
<tr><td>P62</td><td>32841</td><td>3</td><td>13_13_3_6</td><td>32843</td><td>2202</td><td>2202</td><td>4521</td><td>5411</td></tr>
<tr><td rowspan="2">P63</td><td rowspan="2">32874</td><td rowspan="2">46</td><td rowspan="2">11_13_2_8</td><td rowspan="2">32873</td><td colspan="2">1102</td><td>5613</td><td>5613</td></tr>
<tr><td>1102</td><td>2202</td><td>4512</td><td>4512</td></tr>
<tr><td>P64</td><td>32914</td><td>2</td><td>12_12_4_7</td><td>32915</td><td>2202</td><td>2202</td><td>5430</td><td>5613</td></tr>
<tr><td rowspan="2">P65</td><td rowspan="2">32947</td><td rowspan="2">92</td><td rowspan="2">12_14_2_8</td><td rowspan="2">32946</td><td colspan="2">2202</td><td>5621</td><td>5613</td></tr>
<tr><td>1102</td><td>2202</td><td>4521</td><td>5613</td></tr>
<tr><td>P66</td><td>32987</td><td>2</td><td>13_13_2_7</td><td>32988</td><td>1101</td><td>1102</td><td>5612</td><td>6512</td></tr>
<tr><td>P67</td><td>33022</td><td>8</td><td>13_15_2_6</td><td>33021</td><td>2202</td><td>2202</td><td>4511</td><td>5611</td></tr>
<tr><td>P68</td><td>33062</td><td>2</td><td>14_14_2_6</td><td>33062</td><td>2202</td><td>2202</td><td>5411</td><td>5611</td></tr>
<tr><td>P69</td><td>33093</td><td>29</td><td>12_14_3_7</td><td>33093</td><td>1102</td><td>2202</td><td>4521</td><td>5612</td></tr>
<tr><td>P70</td><td>33168</td><td>37</td><td>13_15_3_6</td><td>33167</td><td>2202</td><td>2202</td><td>4521</td><td>5611</td></tr>
<tr><td rowspan="2">P71</td><td rowspan="2">33239</td><td rowspan="2">65</td><td rowspan="2">12_14_2_8</td><td rowspan="2">33239</td><td colspan="2">2202</td><td>5613</td><td>5613</td></tr>
<tr><td>2202</td><td>2202</td><td>4512</td><td>4512</td></tr>
<tr><td>P72</td><td>33313</td><td>2</td><td>13_15_2_7</td><td>33313</td><td>2201</td><td>2202</td><td>4511</td><td>5613</td></tr>
<tr><td>P73</td><td>33355</td><td>1</td><td>14_14_2_6</td><td>33355</td><td>2202</td><td>2202</td><td>5430</td><td>5612</td></tr>
<tr><td>P74</td><td>33530</td><td>11</td><td>12_14_2_9</td><td>33530</td><td>1102</td><td>2202</td><td>4512</td><td>5613</td></tr>
<tr><td>P75</td><td>33605</td><td>23</td><td>13_15_2_8</td><td>33604</td><td>2202</td><td>2202</td><td>4511</td><td>5613</td></tr>
<tr><td>P76</td><td>33894</td><td>2</td><td>13_15_2_9</td><td>33895</td><td>1102</td><td>2202</td><td>5612</td><td>5613</td></tr>
<tr><td>P77</td><td>34187</td><td>1</td><td>13_15_2_10</td><td>34186</td><td>1102</td><td>2202</td><td>5613</td><td>5613</td></tr>
</table>

a. From the total abundances of each glycoform (in three charge states, +9 to +11), which were normalized by P33 (processed using Thermo BioPharma Finder Software$^{TM}$).

**Supplementary Table 6.** Dissociation constants ($K_d$) of glycan ligands (**15**, **21**, **27**, **35**, **36**, **39-42**, **70**, **72**, **77**, **130** and **141**) measured by ESI-MS in 100 mM ammonium acetate (pH 6.9) at 25 °C and 37 °C.[a,b]

| Glycan code | $K_d$ (µM) at 25 °C | $K_d$ (µM) at 37 °C |
|---|---|---|
| **15** | 590 ± 60 | 720 ± 90 |
| **21** | ND | ND |
| **27** | 610 ± 30 | 800 ± 40 |
| **35** | ND | ND |
| **36** | ND | ND |
| **39** | 370 ± 10 | 500 ± 10 |
| **40** | 300 ± 10 | 330 ± 40 |
| **41** | 230 ± 4 | 290 ± 10 |
| **42**[c] | 210 ± 10 | 250 ± 4 |
| **70** | 160 ± 40 | 160 ± 40 |
| **72** | 170 ± 5 | 270 ± 20 |
| **77** | 850 ± 50 | ND |
| **130** | 230 ± 30 | 280 ± 20 |
| **141** | 460 ± 30 | 600 ± 30 |

a. Affinities determined from measurements performed at a minimum of three different initial concentrations. Errors correspond to 1 standard deviation.

b. ND ≡ not detected.

c. The $K_d$ was calculated based on the estimated fractional abundance of **42** in the sample.

**Supplementary Table 7.** Summary of glycans identified HILIC-UHPLC analysis of 2-AB labeled *N*-glycans released from lung tissue and their relative abundances (as released ligands) in CaR-ESI-MS screening against RBD at 25 $^{o}$C and 37 $^{o}$C. The relative abundance ratios (to glycan number **27**) from CaR-ESI-MS screening are indicated as: ○ , ratio ~1 – 10; ◍, 10 – 100; ●, larger than 100.[a,b]

| No. | Glycan composition | Putative structure | Number of isomers | Relative abundance (%) | CaR-ESI-MS | |
|---|---|---|---|---|---|---|
| | | | | | 25 $^{o}$C | 37 $^{o}$C |
| 1 | HexNAc$_2$Hex$_2$Fuc$_1$ | 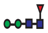 | 1 | 69.2 | ND | ND |
| 2 | HexNAc$_2$Hex$_3$ | 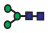 | 1 | 20.1 | ND | ND |
| 3 | HexNAc$_2$Hex$_3$Fuc$_1$ | 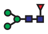 | 1 | 33.3 | ND | ND |
| 4 | HexNAc$_2$Hex$_4$ | 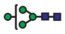 | 2 | 63.4 | ND | ND |
| 5 | HexNAc$_2$Hex$_4$Fuc$_1$ | 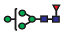 | 2 | 18.9 | ND | ND |
| 6 | HexNAc$_2$Hex$_6$ | 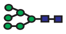 | 1 | 16.2 | ND | ND |
| 7 | HexNAc$_2$Hex$_7$ | 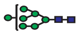 | 1 | 6.2 | ND | ND |
| 8 | HexNAc$_2$Hex$_8$ | 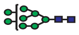 | 1 | 12.0 | ND | ND |
| 9 | HexNAc$_2$Hex$_9$ | 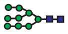 | 1 | 4.5 | ND | ND |
| 10 | HexNAc$_3$Hex$_3$ | 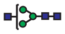 | 2 | 13.9 | ND | ND |
| 11 | HexNAc$_3$Hex$_3$Fuc$_1$ | 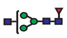 | 1 | 15.2 | ND | ND |
| 12 | HexNAc$_3$Hex$_4$ | 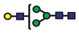 | 1 | 4.7 | ND | ND |
| 13 | HexNAc$_3$Hex$_4$Fuc$_1$Neu5Ac$_1$ | 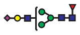 | 1 | 4.1 | ◍ | ◍ |
| 14 | HexNAc$_3$Hex$_4$Neu5Ac$_1$ | 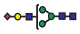 | 1 | 5.7 | ◍ | ○ |
| 15 | HexNAc$_3$Hex$_5$ | 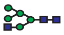 | 1 | 5.8 | ND | ND |
| 16 | HexNAc$_3$Hex$_5$Fuc$_1$Neu5Ac$_1$ | 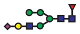 | 2 | 16.7 | ○ | ○ |

| | | | | | | |
|---|---|---|---|---|---|---|
| 17 | HexNAc$_3$Hex$_5$Neu5Ac$_1$ | 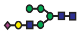 | 1 | 5.5 | ND | ND |
| 18 | HexNAc$_4$Hex$_3$Fuc$_1$ | 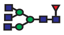 | 1 | 18.0 | ND | ND |
| 19 | HexNAc$_4$Hex$_4$ | 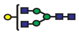 | 1 | 13.2 | ND | ND |
| 20 | HexNAc$_4$Hex$_4$Fuc$_1$ | 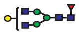 | 1 | 12.6 | ND | ND |
| 21 | HexNAc$_4$Hex$_4$Fuc$_1$Neu5Ac$_1$ | 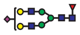 | 2 | 12.2 | ◍ | ○ |
| 22 | HexNAc$_4$Hex$_4$Neu5Ac$_1$ | 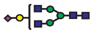 | 1 | 2.8 | ○ | ○ |
| 23 | HexNAc$_4$Hex$_5$ | 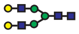 | 1 | 9.3 | ND | ND |
| 24 | HexNAc$_4$Hex$_5$Fuc$_1$ | 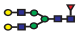 | 2 | 61.3 | ND | ND |
| 25 | HexNAc$_4$Hex$_5$Fuc$_1$Neu5Ac$_1$ | 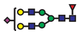 | 3 | 100.0 | ◍ | ◍ |
| 26 | HexNAc$_4$Hex$_5$Fuc$_1$Neu5Ac$_2$ | 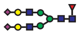 | 2 | 40.6 | ● | ● |
| 27 | HexNAc$_4$Hex$_5$Fuc$_2$Neu5Ac$_1$ | 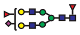 | 1 | 1.0 | ○ | ○ |
| 28 | HexNAc$_4$Hex$_5$Neu5Ac$_1$ | 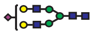 | 1 | 8.3 | ◍ | ◍ |
| 29 | HexNAc$_4$Hex$_5$Neu5Ac$_2$ | 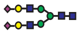 | 2 | 21.8 | ● | ◍ |
| 30 | HexNAc$_5$Hex$_3$Fuc$_1$ | 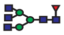 | 2 | 21.9 | ND | ND |
| 31 | HexNAc$_5$Hex$_4$ | 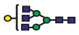 | 1 | 5.7 | ND | ND |
| 32 | HexNAc$_5$Hex$_4$Fuc$_1$ | 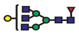 | 1 | 7.2 | ND | ND |
| 33 | HexNAc$_5$Hex$_4$Fuc$_1$Neu5Ac$_1$ | 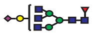 | 1 | 0.5 | ○ | ○ |
| 34 | HexNAc$_5$Hex$_5$Fuc$_1$Neu5Ac$_1$ | 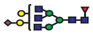 | 1 | 3.4 | ○ | ○ |
| 35 | HexNAc$_5$Hex$_5$Fuc$_1$Neu5Ac$_2$ | 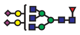 | 1 | 2.0 | ○ | ○ |
| 36 | HexNAc$_5$Hex$_6$Fuc$_1$Neu5Ac$_1$ | 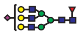 | 1 | 1.2 | ND | ND |
| 37 | HexNAc$_5$Hex$_6$Fuc$_1$Neu5Ac$_2$ | 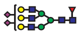 | 5 | 5.9 | ○ | ○ |
| 38 | HexNAc$_5$Hex$_6$Fuc$_1$Neu5Ac$_3$ | 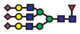 | 4 | 2.6 | ○ | ○ |
| 39 | HexNAc$_5$Hex$_6$Fuc$_1$Neu5Ac$_4$ | 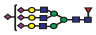 | 1 | 0.4 | ND | ND |
| 40 | HexNAc$_5$Hex$_6$Neu5Ac$_3$ | 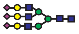 | 1 | 0.8 | ○ | ○ |

| | | | | | | |
|---|---|---|---|---|---|---|
| 41 | $HexNAc_6Hex_7Fuc_1Neu5Ac_2$ | 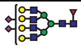 | 2 | 0.6 | ND | ND |
| 42 | $HexNAc_6Hex_7Fuc_1Neu5Ac_4$ | 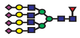 | 1 | 0.1 | ND | ND |
| 43 | $HexNAc_7Hex_5$ | 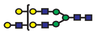 | 1 | 0.3 | ND | ND |
| 44 | $HexNAc_7Hex_8Fuc_1Neu5Ac_1$ | 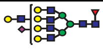 | 1 | 0.1 | ND | ND |
| 45 | $HexNAc_7Hex_8Fuc_1Neu5Ac_2$ | 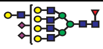 | 1 | 0.1 | ND | ND |
| 46 | $HexNAc_8Hex_6$ | 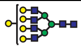 | 1 | 0.3 | ND | ND |
| 47 | $HexNAc_8Hex_6Neu5Ac_1$ | 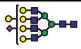 | 1 | 0.1 | ND | ND |
| 48 | $HexNAc_3Hex_3Fuc_1Neu5Ac_1$ | 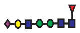 | ND | ND | ○ | ○ |
| 49 | $HexNAc_3Hex_6Neu5Ac_1$ | 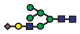 | ND | ND | ○ | ○ |

a. ND ≡ not detected

b. *N*-glycans could have either core or antennary fucosylation.

**Supplementary Table 8.** Summary of glycans identified HILIC-UHPLC analysis of 2-AB labeled *N*-glycans released from intestinal tissue and their presence in CaR-ESI-MS screening against RBD at 25ºC and 37 ºC. The relative abundance ratios (to glycan number **44**) of the released glycans from CaR-ESI-MS screening are indicated as: ○ , ratio ~1 – 10; ◍, 10 – 100; ●, larger than 100.[a,b]

| No. | Glycan composition | Putative structure | Number of isomers | Relative abundance (%) | CaR-ESI-MS | |
|---|---|---|---|---|---|---|
| | | | | | 25 ºC | 37 ºC |
| 1 | HexNAc2Hex2 |  | 1 | 7.8 | ND | ND |
| 2 | HexNAc2Hex2Fuc1 | 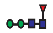 | 1 | 14.9 | ND | ND |
| 3 | HexNAc2Hex3 | 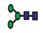 | 1 | 4.7 | ND | ND |
| 4 | HexNAc2Hex3Fuc1 | 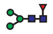 | 1 | 6.6 | ND | ND |
| 5 | HexNAc2Hex4 | 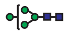 | 2 | 7.4 | ND | ND |
| 6 | HexNAc2Hex5 | 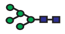 | 2 | 15.0 | ND | ND |
| 7 | HexNAc2Hex6 | 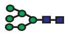 | 1 | 14.9 | ND | ND |
| 8 | HexNAc2Hex7 | 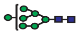 | 1 | 4.2 | ND | ND |
| 9 | HexNAc2Hex8 | 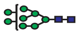 | 1 | 3.3 | ND | ND |
| 10 | HexNAc2Hex9 | 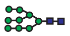 | 1 | 11.5 | ND | ND |
| 11 | HexNAc3Hex3Fuc1 | 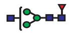 | 1 | 4.9 | ND | ND |
| 12 | HexNAc3Hex4 | 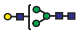 | 1 | 0.4 | ND | ND |
| 13 | HexNAc3Hex4Neu5Ac1 | 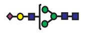 | 2 | 1.3 | ○ | ◍ |
| 14 | HexNAc3Hex4Fuc1Neu5Ac1 | 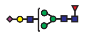 | 2 | 3.8 | ◍ | ◍ |
| 15 | HexNAc3Hex5Neu5Ac1 | 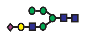 | 2 | 6.4 | ○ | ◍ |
| 16 | HexNAc4Hex3Fuc1 | 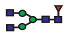 | 2 | 16.0 | ND | ND |
| 17 | HexNAc4Hex4Neu5Ac1 | 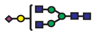 | 1 | 1.3 | ○ | ◍ |
| 18 | HexNAc4Hex4Fuc1 | 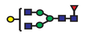 | 2 | 16.0 | ND | ND |
| 19 | HexNAc4Hex4Fuc1Neu5Ac1 | 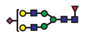 | 2 | 6.1 | ◍ | ◍ |
| 20 | HexNAc4Hex5 | 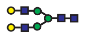 | 1 | 6.3 | ND | ND |

| | | | | | | |
|---|---|---|---|---|---|---|
| 21 | HexNAc4Hex5Neu5Ac1 | 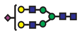 | 2 | 33.6 | ◍ | ◍ |
| 22 | HexNAc4Hex5Neu5Ac2 | 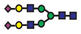 | 3 | 100.0 | ● | ● |
| 23 | HexNAc4Hex5Fuc1 | 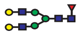 | 1 | 19.9 | ND | ND |
| 24 | HexNAc4Hex5Fuc1Neu5Ac1 | 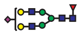 | 2 | 48.0 | ◍ | ND |
| 25 | HexNAc4Hex5Fuc1Neu5Ac1S1 | 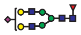 | 1 | 3.9 | ○ | ND |
| 26 | HexNAc4Hex5Fuc1Neu5Ac2 | 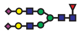 | 4 | 42.4 | ◍ | ◍ |
| 27 | HexNAc5Hex3 | 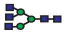 | 1 | 2.6 | ND | ND |
| 28 | HexNAc5Hex4Neu5Ac1 | 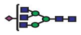 | 1 | 0.9 | ○ | ND |
| 29 | HexNAc5Hex4Fuc1 | 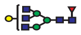 | 1 | 6.7 | ND | ND |
| 30 | HexNAc5Hex4Fuc1Neu5Ac1 | 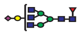 | 2 | 4.2 | ○ | ○ |
| 31 | HexNAc5Hex5Fuc1 | 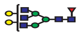 | 1 | 5.3 | ND | ND |
| 32 | HexNAc5Hex5Fuc1Neu5Ac1 | 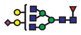 | 1 | 4.5 | ○ | ND |
| 33 | HexNAc5Hex5Fuc1Neu5Ac2 | 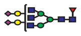 | 1 | 6.0 | ○ | ND |
| 34 | HexNAc5Hex6Neu5Ac1 | 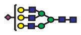 | 1 | 2.8 | ○ | ND |
| 35 | HexNAc5Hex6Neu5Ac2 | 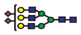 | 3 | 22.9 | ○ | ND |
| 36 | HexNAc5Hex6Neu5Ac3 | 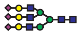 | 2 | 46.6 | ◍ | ◍ |
| 37 | HexNAc5Hex6Fuc1 | 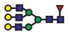 | 1 | 7.6 | ND | ND |
| 38 | HexNAc5Hex6Fuc1Neu5Ac1 | 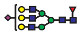 | 3 | 10.1 | ○ | ○ |
| 39 | HexNAc5Hex6Fuc1Neu5Ac2 | 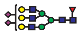 | 5 | 30.6 | ◍ | ○ |
| 40 | HexNAc5Hex6Fuc1Neu5Ac3 | 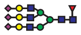 | 2 | 50.0 | ◍ | ◍ |
| 41 | HexNAc5Hex6Fuc2Neu5Ac3 | 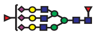 | 1 | 6.8 | ND | ND |
| 42 | HexNAc6Hex3Fuc1 | 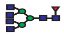 | 2 | 19.4 | ND | ND |
| 43 | HexNAc6Hex7Neu5Ac2 | 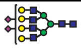 | 1 | 2.9 | ○ | ○ |
| 44 | HexNAc6Hex7Neu5Ac3 | 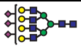 | 2 | 9.7 | ○ | ○ |
| 45 | HexNAc6Hex7Neu5Ac4 | 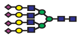 | 2 | 9.8 | ND | ND |
| 46 | HexNAc6Hex7Fuc1Neu5Ac2 | 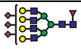 | 1 | 1.1 | ○ | ○ |
| 47 | HexNAc6Hex7Fuc1Neu5Ac3 | 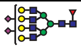 | 3 | 14.6 | ○ | ○ |
| 48 | HexNAc6Hex7Fuc1Neu5Ac4 | 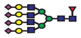 | 2 | 16.2 | ND | ND |
| 49 | HexNAc6Hex7Fuc2Neu5Ac3 | 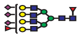 | 1 | 3.0 | ○ | ○ |

| | | | | | | |
|---|---|---|---|---|---|---|
| 50 | $HexNAc_6Hex_7Fuc_2Neu5Ac_4$ | 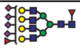 | 2 | 9.3 | ND | ND |
| 51 | $HexNAc_8Hex_9Neu5Ac_3$ | 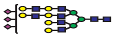 | 1 | 7.0 | ND | ND |
| 52 | $HexNAc_3Hex_3Fuc_1Neu5Ac_1$ | 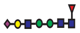 | ND | ND | ○ | ○ |
| 53 | $HexNAc_3Hex_6Neu5Ac_1$ | 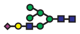 | ND | ND | ○ | ○ |

a. ND ≡ not detected

b. *N*-glycans could have either core or antennary fucosylation.

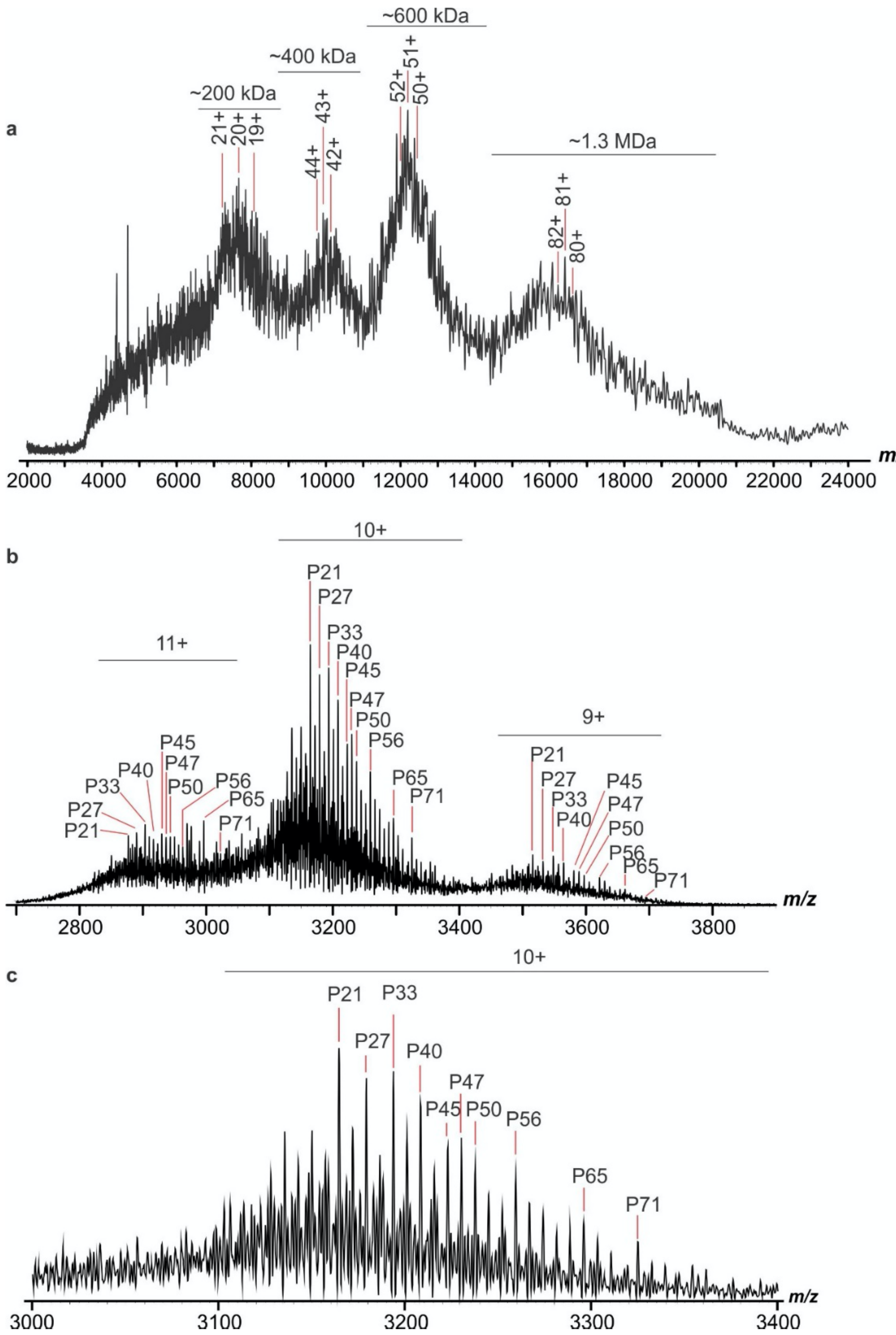


**Supplementary Figure 1. a,b,** Representative ESI mass spectra acquired in positive mode for an aqueous ammonium acetate solution (100 mM, pH 6.9, 25 $^{o}$C) of S-protein (5 μM) (**a**), and RBD (5 μM) (**b**). **c,** Expanded region (m/z 3000 to 3400) of the mass spectrum shown in (**b**).

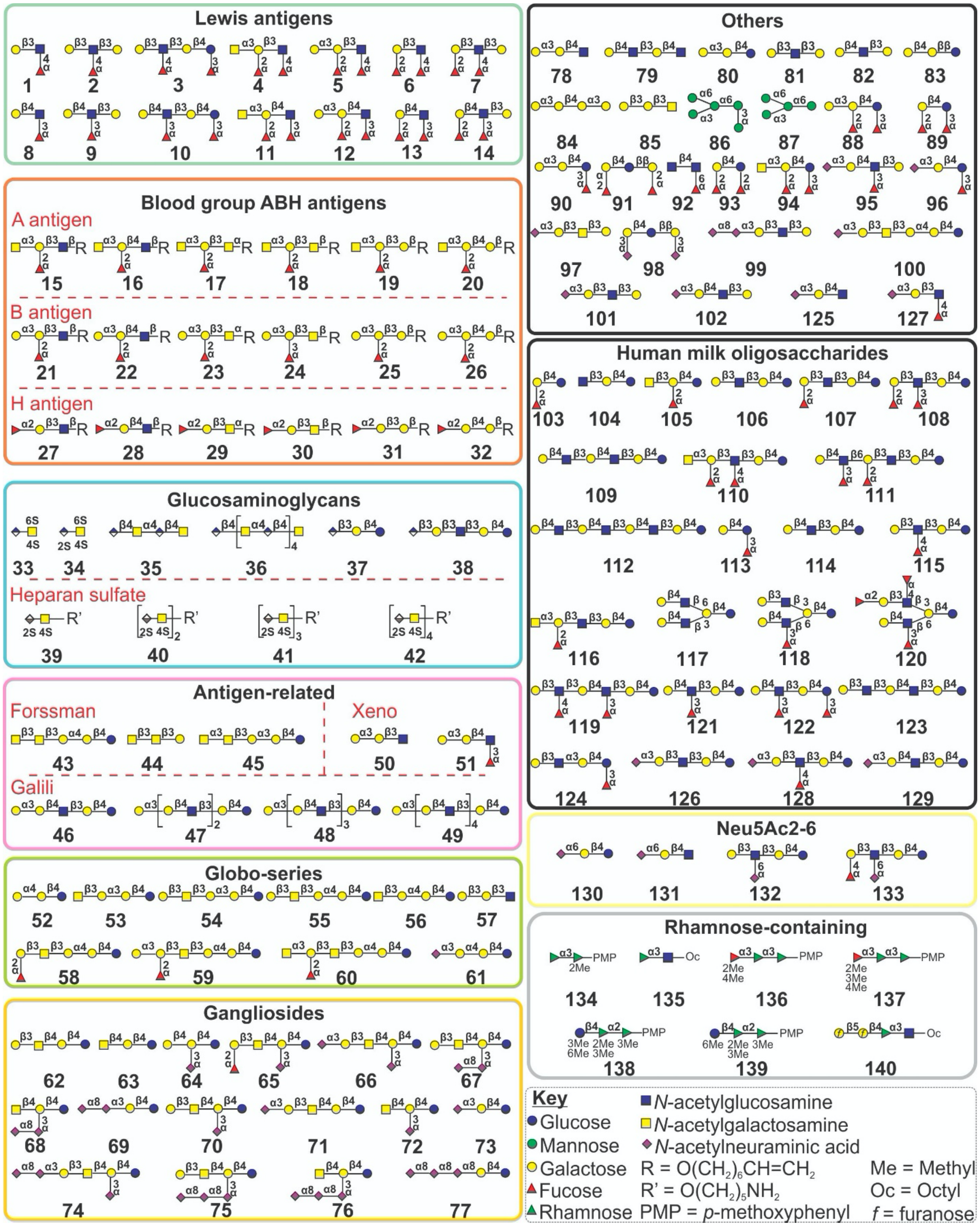


**Supplementary Figure 2.** Structures of 140 defined glycans used in CaR-ESI-MS screening against RBD. These glycans comprise 9 groups: Lewis antigens, blood group ABH antigens, antigen-related glycans, globo-series glycans, ganglioside oligosaccharides, human milk oligosaccharides, Neu5Acα2-6 linked glycans, rhamnose-containing glycans (Library P) and others.

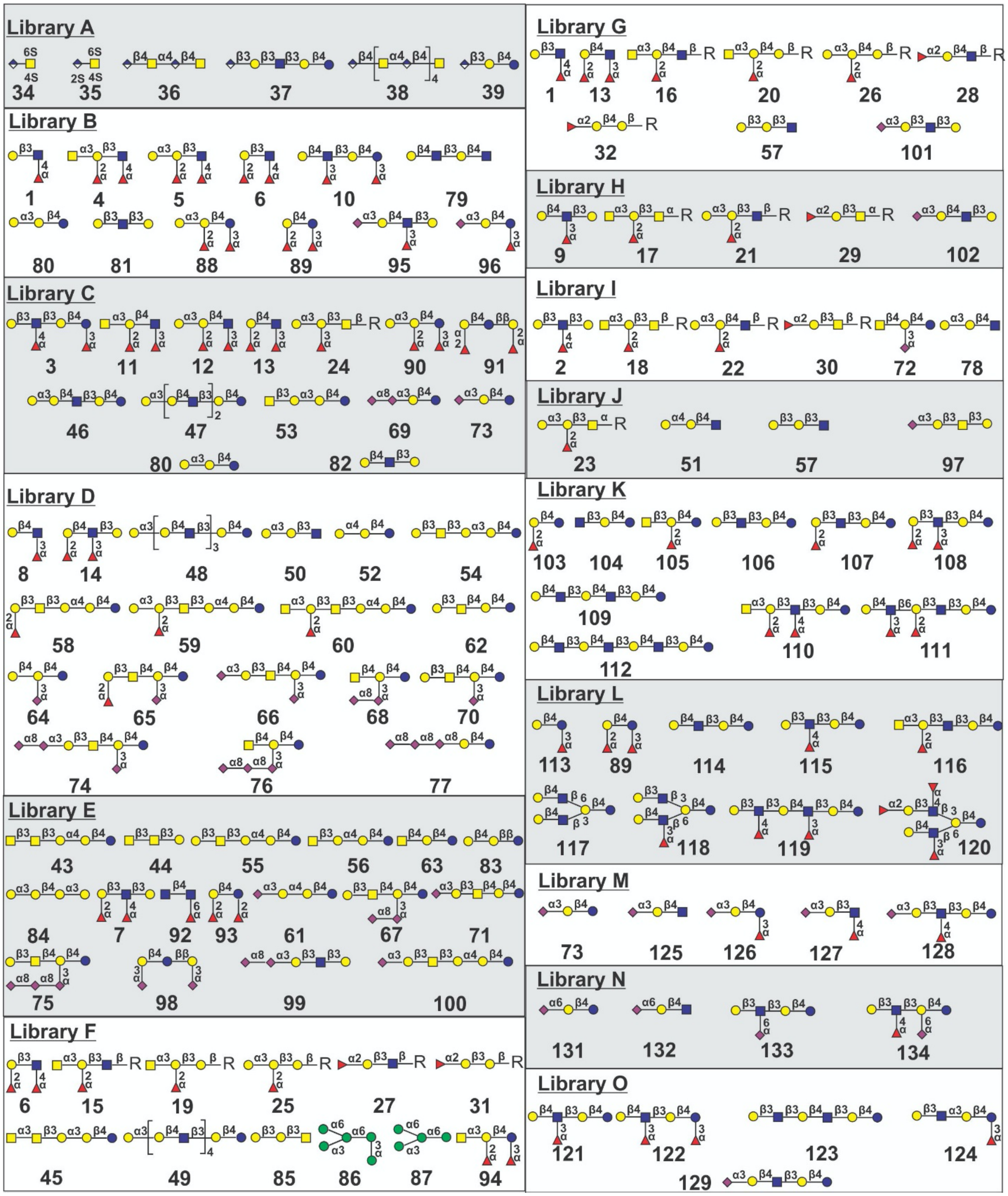


**Supplementary Figure 3.** Sub-libraries (A-O) of defined glycans used for CaR-ESI-MS screening.

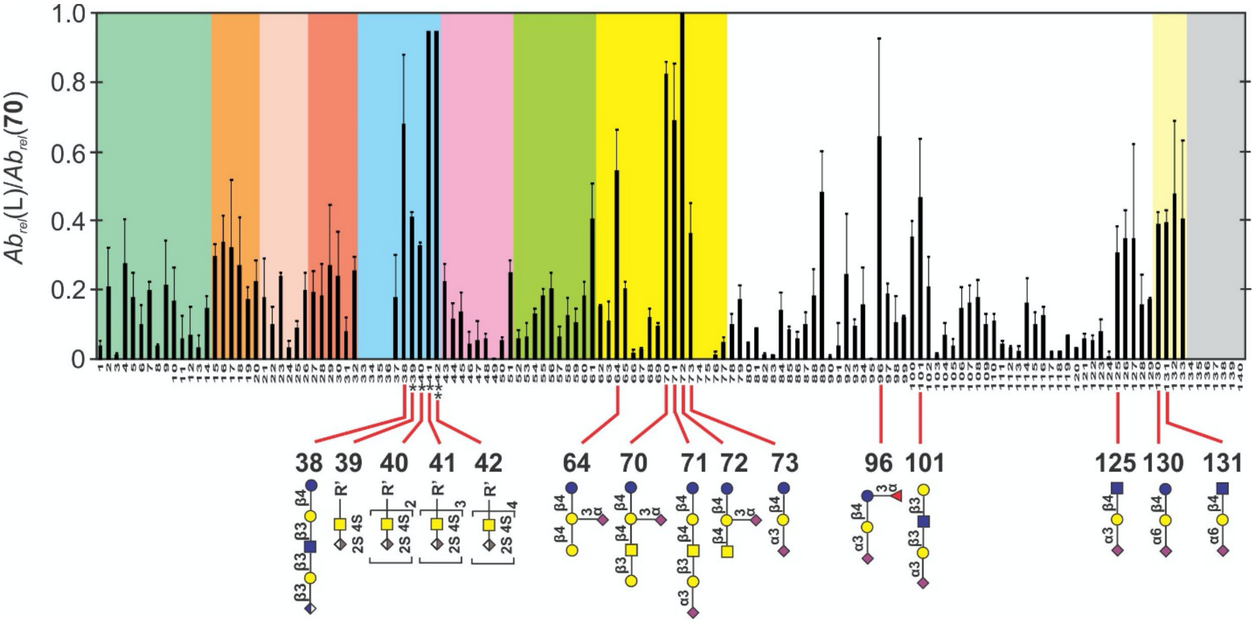


**Supplementary Figure 4. Glycan library screening for RBD.** Normalized abundances of released glycans from the SARS-CoV-2 RBD by CaR-ESI-MS at 25 $^{o}$C. Summary of the charge-normalized (relative to 70) abundances of released ligands measured by CaR-ESI-MS screening of *Library* A – O against SARS-CoV-2 RBD. Measurements were performed in negative ion mode with a UHMR Orbitrap mass spectrometer at an HCD energy of 50 V. Aqueous ammonium acetate (100 mM, pH 6.9) solutions of SARS-CoV-2 RBD (13 μM) and glycan library (containing 50 nM of each glycan) at were used for screening. The different classes of oligosaccharides are distinguished by colour: mint green - Lewis antigens (**1**-**14**); light orange - blood group A antigens (**15**-**20**); light pink - blood group B antigens (**21**-**26**); tropical pink - blood group H antigens (**27**-**32**); ice blue - sulfated compounds (**34**-**38**) and heparan sulfates (**39**-**42**), light violet - antigen-related glycans (**43**-**51**), chartreuse - globo (**52**-**61**), yellow - ganglioside oligosaccharides (**62**-**77**), white – HMOs and other glycans (**78**-**129**), light yellow - Neu5Acα2-6-linked oligosaccharides (**130**-**133**), and grey – rhamnose-containing compounds (**134**-**140**). * relative abundances (in CaR-ESI-MS) estimated from their relative (to **70**) $K_d$ values. ** **42** sample also contained some **41** and the affinity reflects the estimated relative concentrations of **41** and **42**.

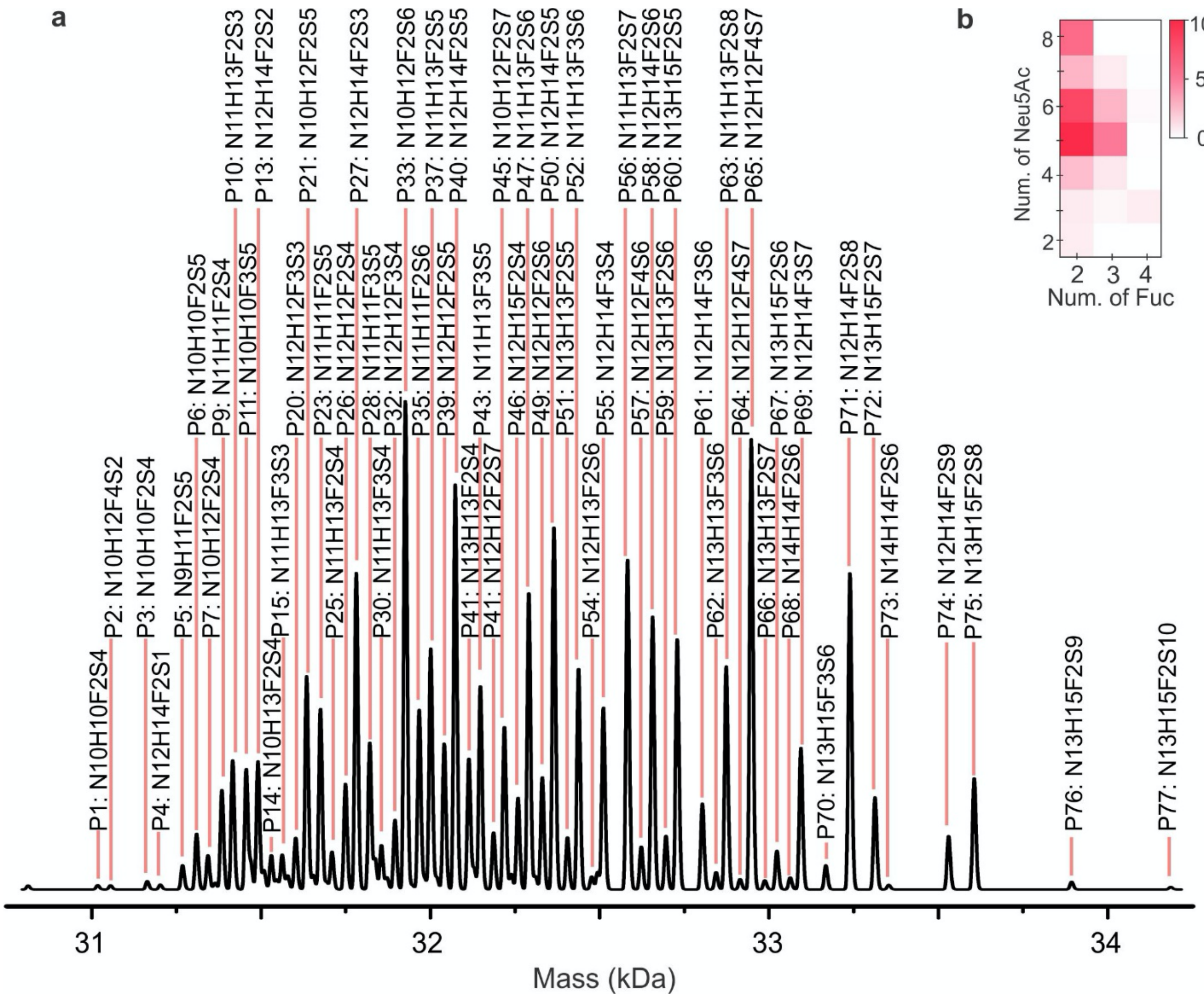


**Supplementary Figure 5. a,** Zero-charge mass spectrum of RBD acquired in positive ion mode with a UHMR Orbitrap mass spectrometer for an aqueous ammonium acetate (100 mM, pH 6.9) solution of SARS-CoV-2 RBD (13 μM). Deconvolution was performed using the Thermo BioPharma Finder Software™. The glycan composition of each species is indicated (N ≡ HexNAc, H ≡ Hex, F ≡ Fuc, S ≡ Neu5Ac); compositions highlighted in red correspond to the species used for direct ESI-MS binding measurements. **b,** Heatmap showing the distribution of the number of Neu5Ac and Fuc residues in the RBD glycoforms.

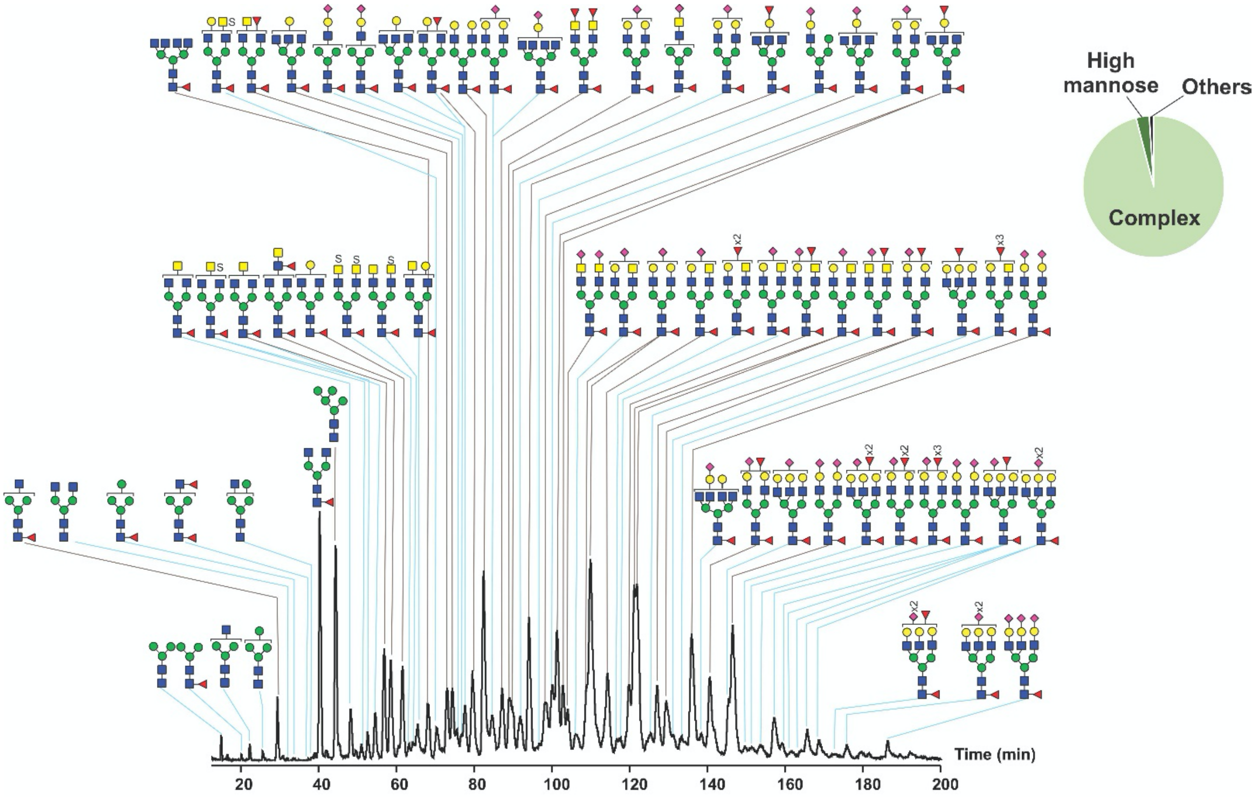


**Supplementary Figure 6.** Chromatogram of 2-AB-labeled *N*-glycans released from SARS-CoV-2 RBD treated with PNGase F acquired using HILIC-UHPLC with fluorescence detection. Summary of the relative abundances of principal *N*-glycan types also shown.

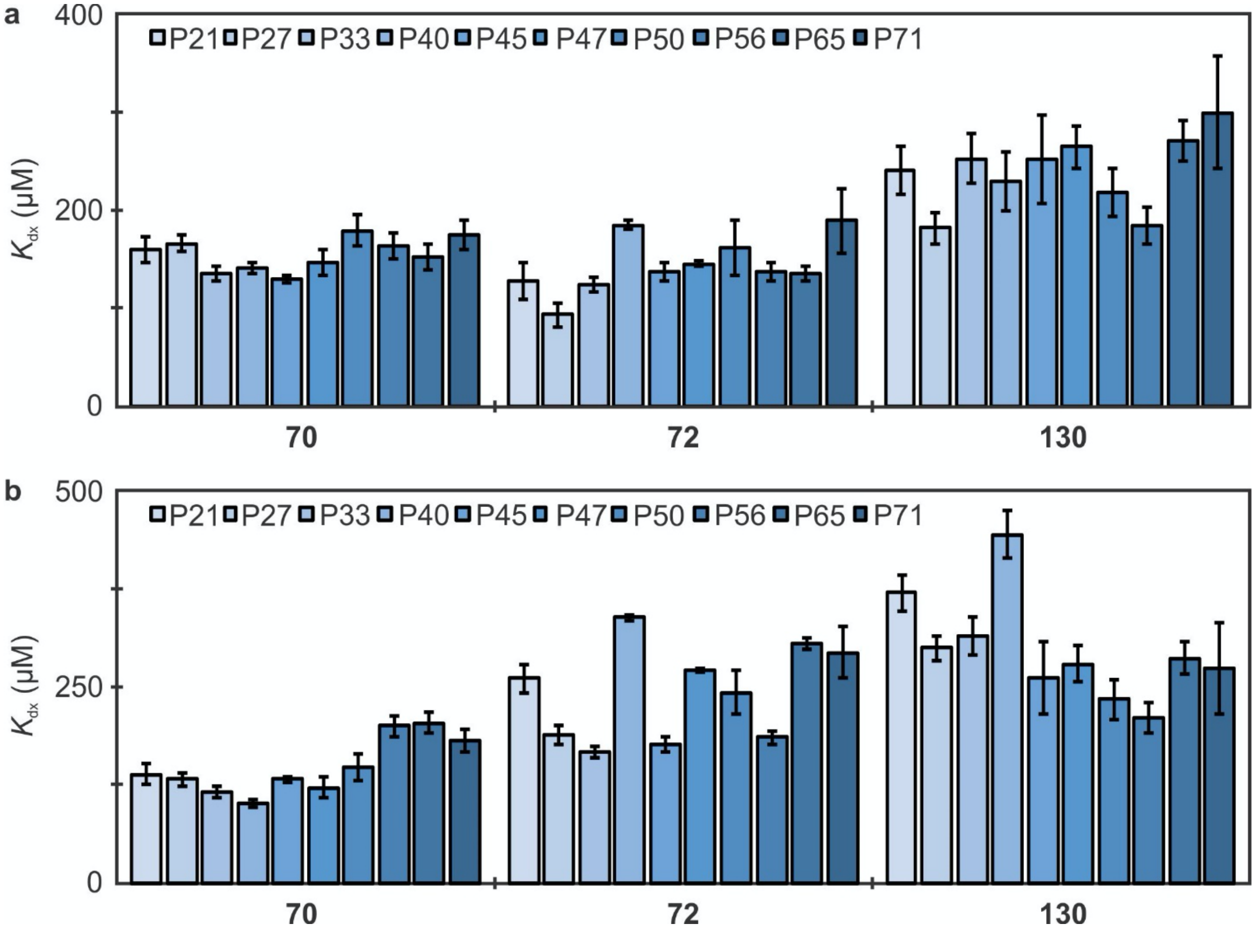


**Supplementary Figure 7.** The $K_d$ of 10 most abundant RBD glycoforms (P21, P27, P33, P40, P45, P47, P50, P56, P65, and P71) in 100 mM ammonium acetate solution (pH 6.9) and RBD (5 μM) at 3 different concentrations of glycans **70**, **72**, and **130** at 25 $^oC$ (**a**) and 37 $^oC$ (**b**).

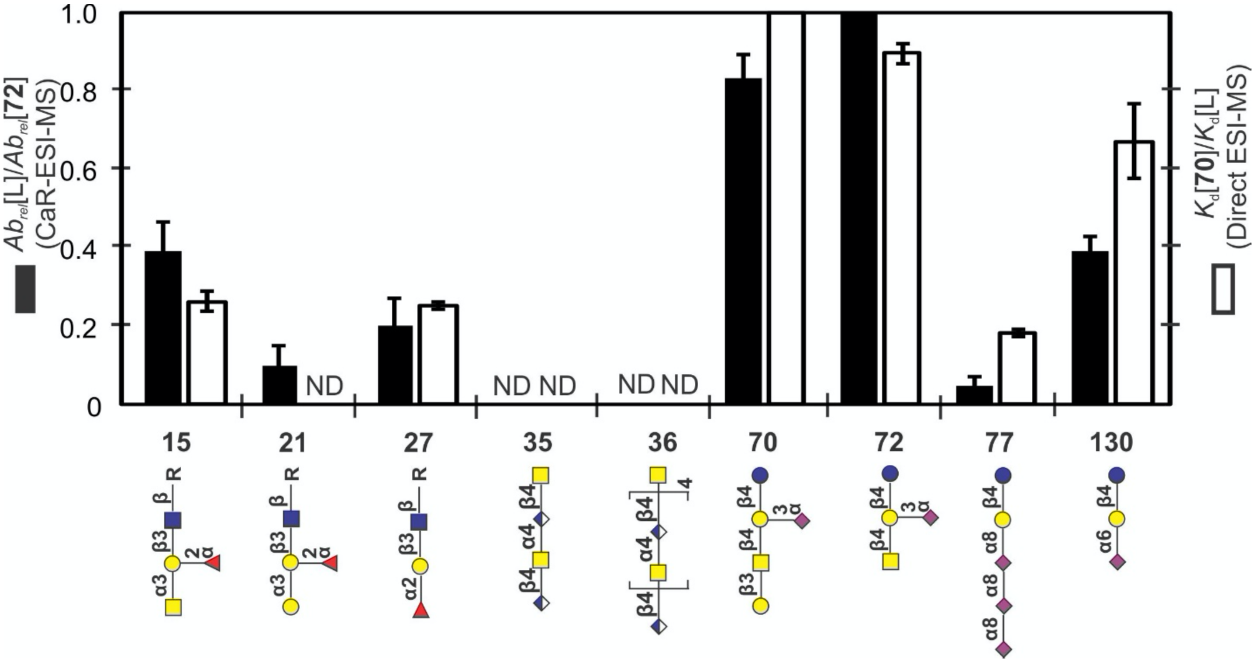


**Supplementary Figure 8.** Comparison of glycan (**15**, **21**, **27**, **35**, **36**, **70**, **72**, **77** and **130**) affinities for RBD measured by ESI-MS and their relative abundances measured by CaR-ESI-MS screening (performed at 25 °C). Black bars represent the ratio of the abundance of released glycans normalized to that of **72** under same conditions. White bars represent the ratios of dissociation affinities of RBD against **70** to dissociation affinities of RBD against glycan (L). ND ≡ not detected.

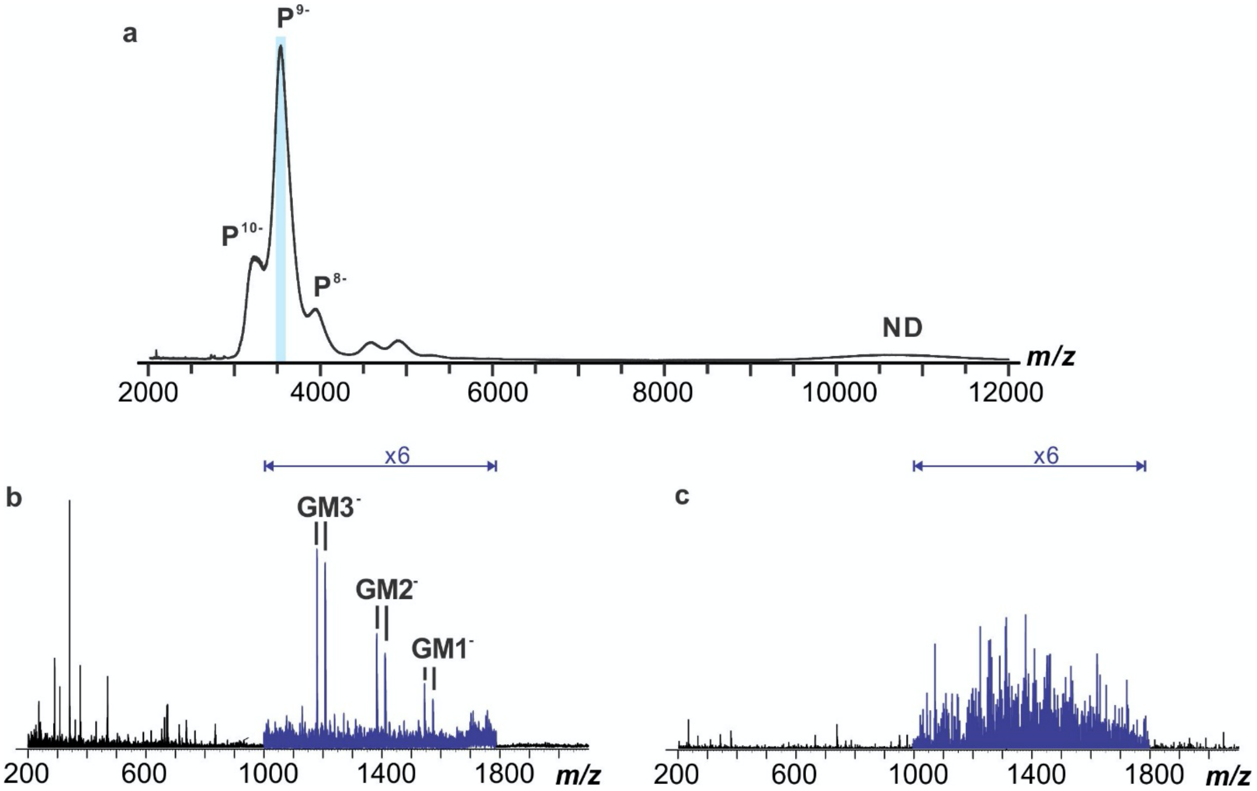


**Supplementary Figure 9. a,** ESI mass spectrum acquired in negative ion mode for an aqueous ammonium acetate solution (100 mM, 25 °C and pH 7.4) of RBD (13 µM) and 5 µM ganglioside-containing ND (1% GM1, GM2, GM3, GD1a, GD2, and GT1b, each at approximately 10 µM). **b,** CID mass spectrum acquired for ions centered at m/z 3,540, with a window width of 100 m/z units (the region highlighted in light blue in (**a**)) using a Trap collision energy of 50 V. **c,** CID mass spectrum acquired in negative ion mode for ions centered at m/z 3,540, with a window width of 100 m/z units, produced from an aqueous ammonium acetate solution (100 mM, 25 °C and pH 7.4) of and 10 µM ganglioside-containing ND; the Trap collision energy was 50 V.

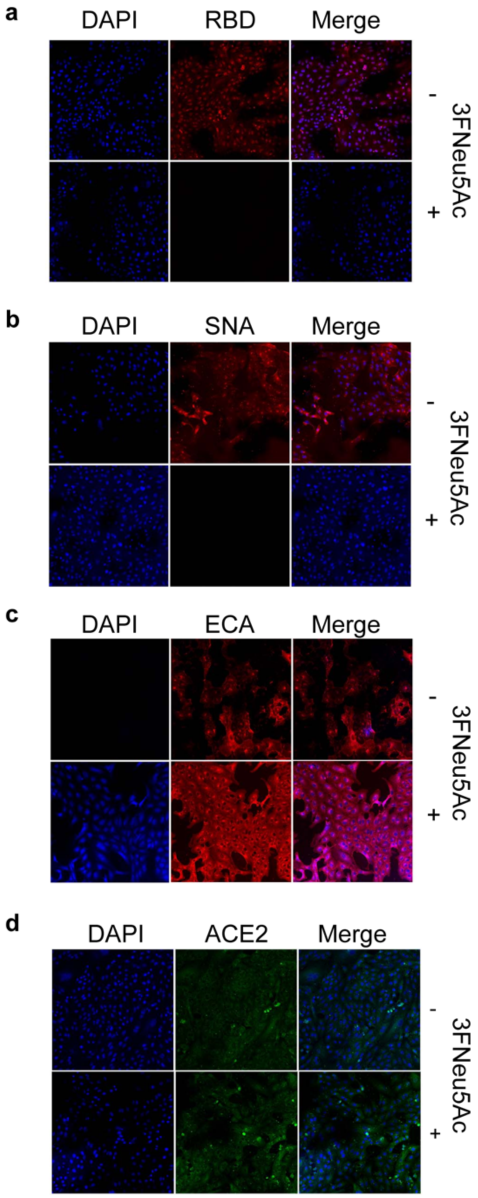


**Supplementary Figure 10. Immunofluorescence staining in 3FNeu5Ac treated Vero-E6 cells.** DAPI was used for nuclear staining. **a,b,** RBD binding (**a**) and SNA (**b**) staining decreased with 3FNeu5Ac treatment. **c,** ECA staining increased with 3FNeu5Ac treatment. **d,** No changes were seen in ACE2 binding when Vero-E6 cells were treated with 3FNeu5Ac.

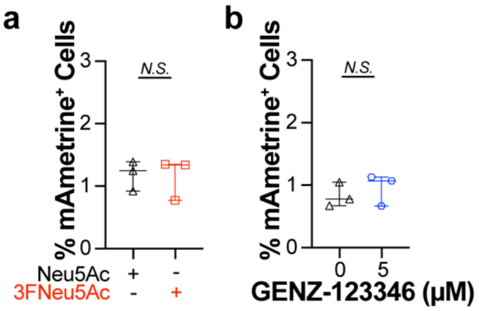


**Supplementary Figure 11. Infection of control lentivirus that does not encode SARS-CoV-2 in HEK293 cells expressing ACE2. a,** RP172 lentivirus (10 µL) 1 hr infection followed by a 23 hr incubation showed no differences when cells are treated with Neu5Ac (300 µM) or 3FNeu5Ac (300 µM). Level of infection quantified by % of mAmetrine+ cells as determined by flow cytometry. **b,** DMSO or GENZ-123346 (5 µM) treated HEK293 cells expressing ACE2 infected with RP172 lentivirus for 1 hr followed by a 23 hr incubation. Infection quantified by % of mAmetrine+ cells observed using flow cytometry. Error bars represent ± standard deviation of three replicates. Statistical significance calculated based on two-tailed unpaired Student's *t*-test.

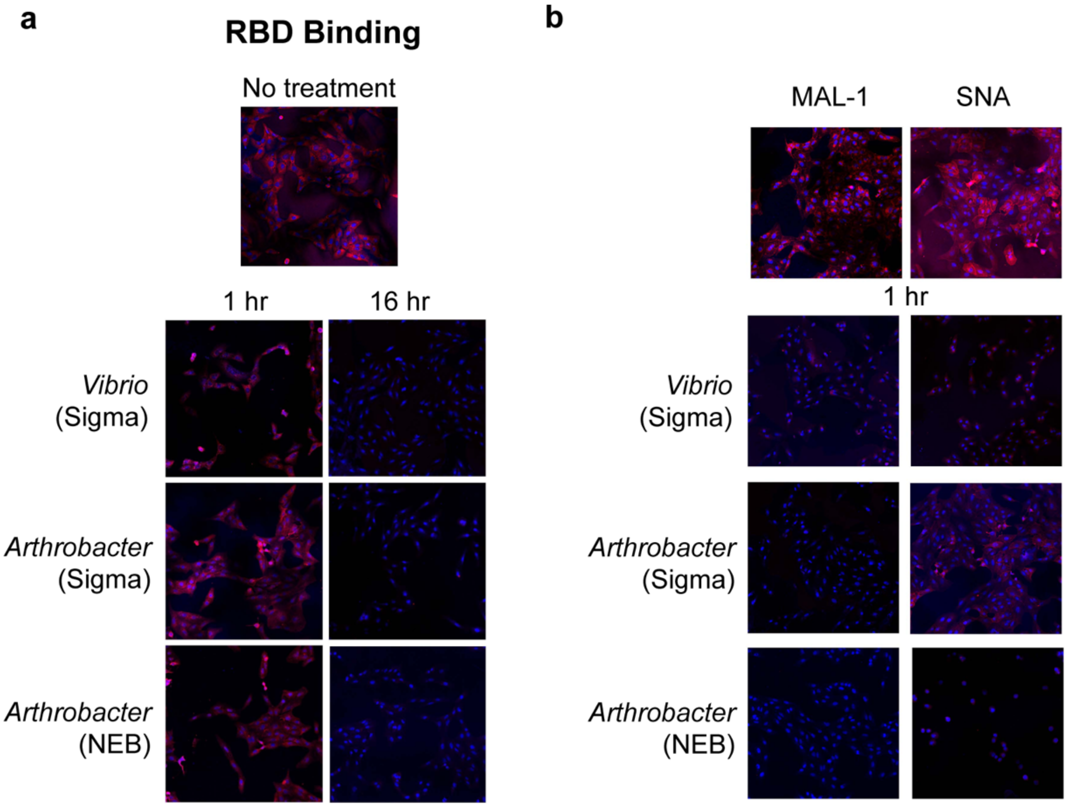


**Supplementary Figure 12. Immunofluorescence staining showing RBD binding in neuraminidase treated Vero-E6 cells. a**, SARS-CoV-2 RBD binding was decreased in Vero-E6 cells treated with *Vibrio* (Sigma), *Arthobacter* (Sigma), and *Arthrobacter* (New England Biolabs, NEB) neuraminidases for 16 hours. **b,** After 1 h of neuraminidase treatment, with *Vibrio* (Sigma), *Arthobacter* (Sigma), or *Arthrobacter* (NEB), significant changes in RBD binding were not observed in Vero-E6 cells.

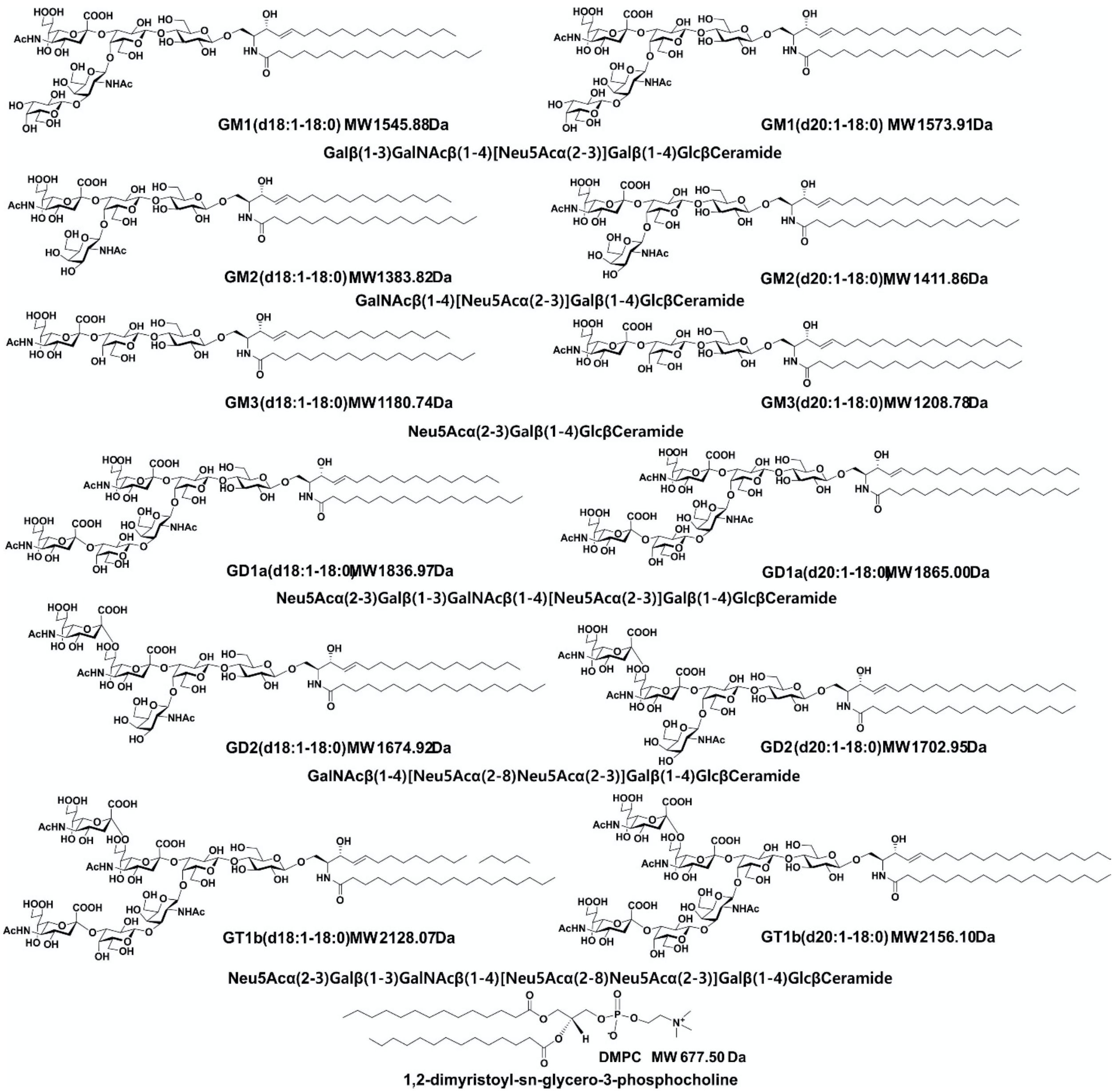


**Supplementary Figure 13.** Structures of the gangliosides (GM1, GM2, GM3, GD1a, GD2 and GT1b; two major isoforms (*d*18:1-18:0 and *d*20:1-18:0) shown), and the phospholipid (DMPC) used in this study.

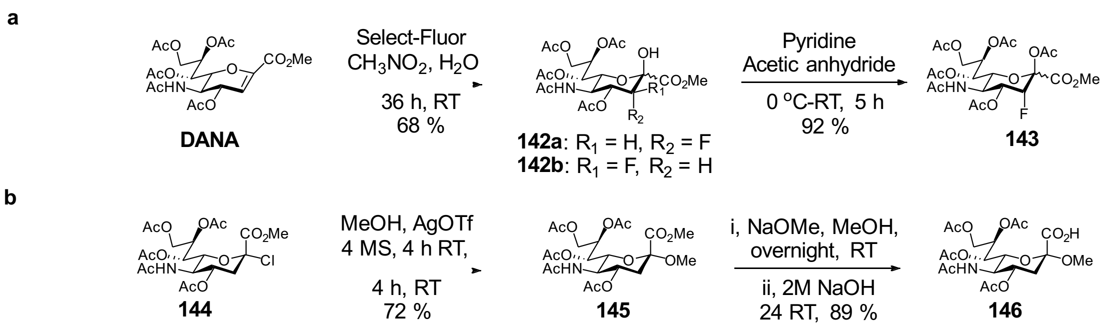

**Supplementary Figure 14**. Synthesis scheme of per-*O*-acetylated 3$F_{ax}$-Neu5Ac (**143**) (**a**), and methyl-α-Neu5Ac (**146**) (**b**).

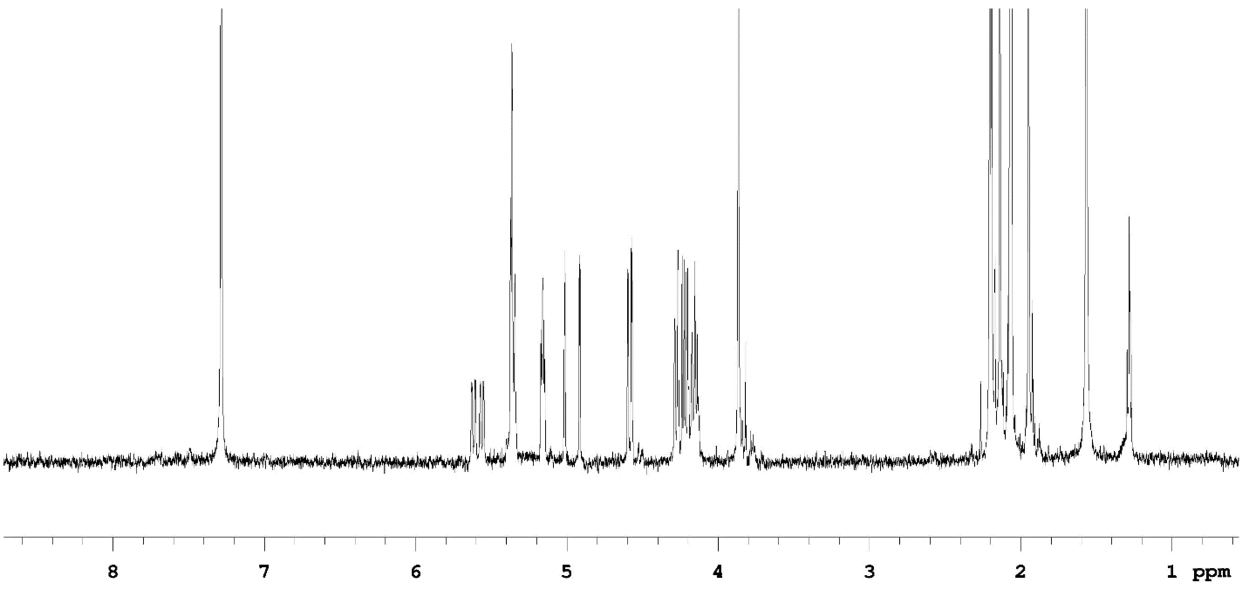


**Supplementary Figure 15.** $^{1}$H NMR spectrum of per-*O*-acetylated 3Fax-Neu5Ac (**143**) (600 MHz, $D_2O$)

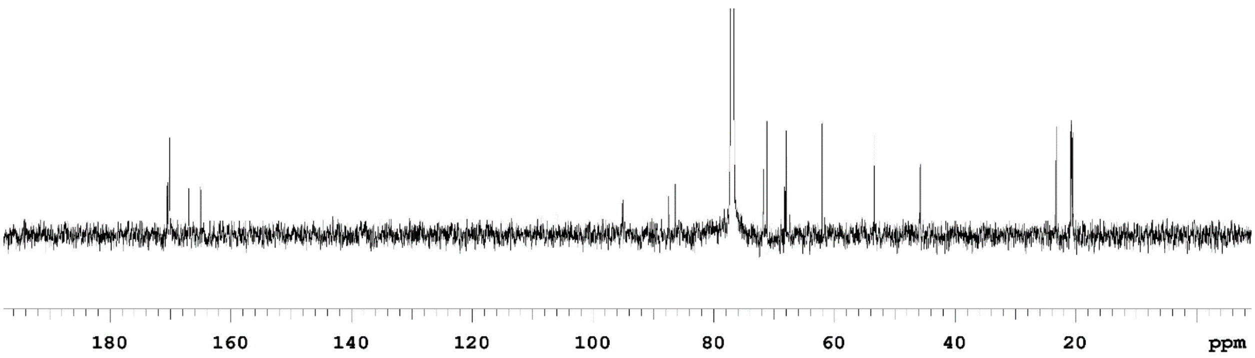


**Supplementary Figure 16.** $^{13}C$ NMR spectrum of per-*O*-acetylated 3Fax-Neu5Ac (**143**) (125 MHz, $D_2O$)

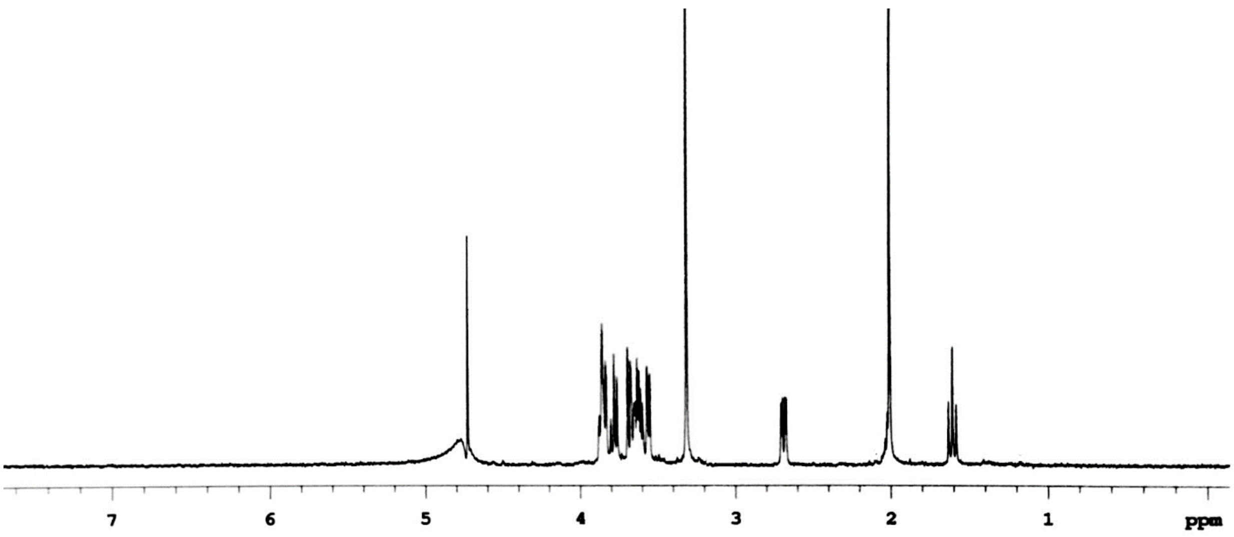


**Supplementary Figure 17.** $^{1}$H NMR spectrum of methyl-α-Neu5Ac (**146**) (500 MHz, $D_2O$).

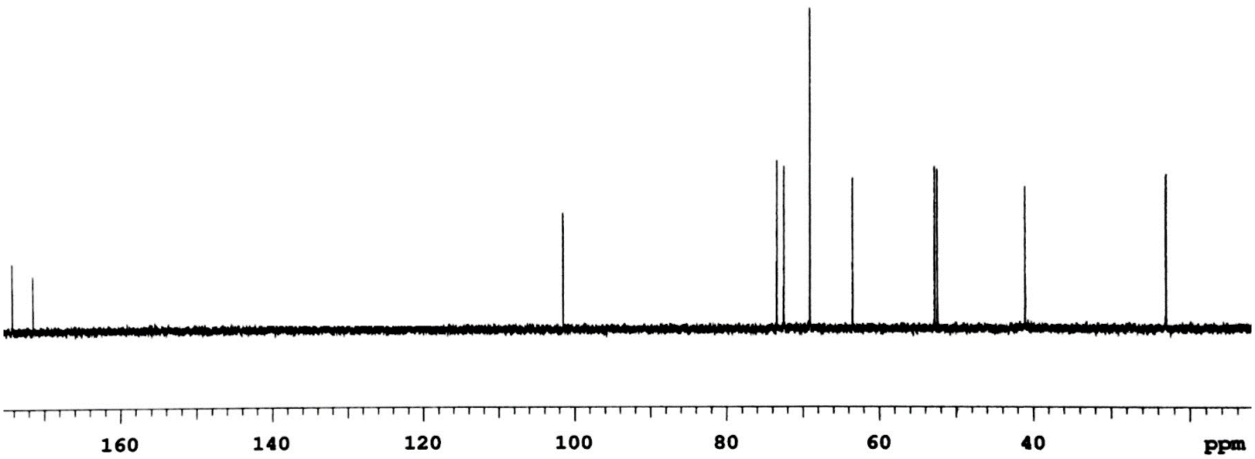


**Supplementary Figure 18.** $^{13}C$ NMR spectrum of methyl-α-Neu5Ac (**146**) (125 MHz, $D_2O$).

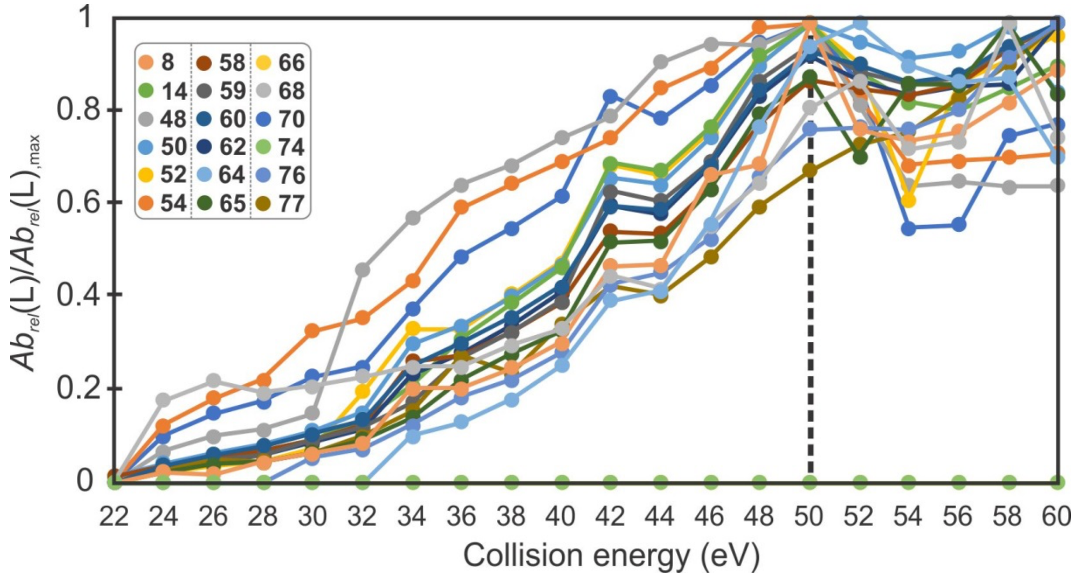


**Supplementary Figure 19.** Influence of collision energy (eV) on glycan ligand release for CaR-ESI-MS screening of RBD (13 μM) against Library D (50 nM of each glycan) in ammonium acetate (100 mM, pH 6.9) containing SNA ($P_{ref}$, 4 μM) measured at 25 °C. The abundance of each glycan was normalized according to the maximum, charge-normalized signal detected.

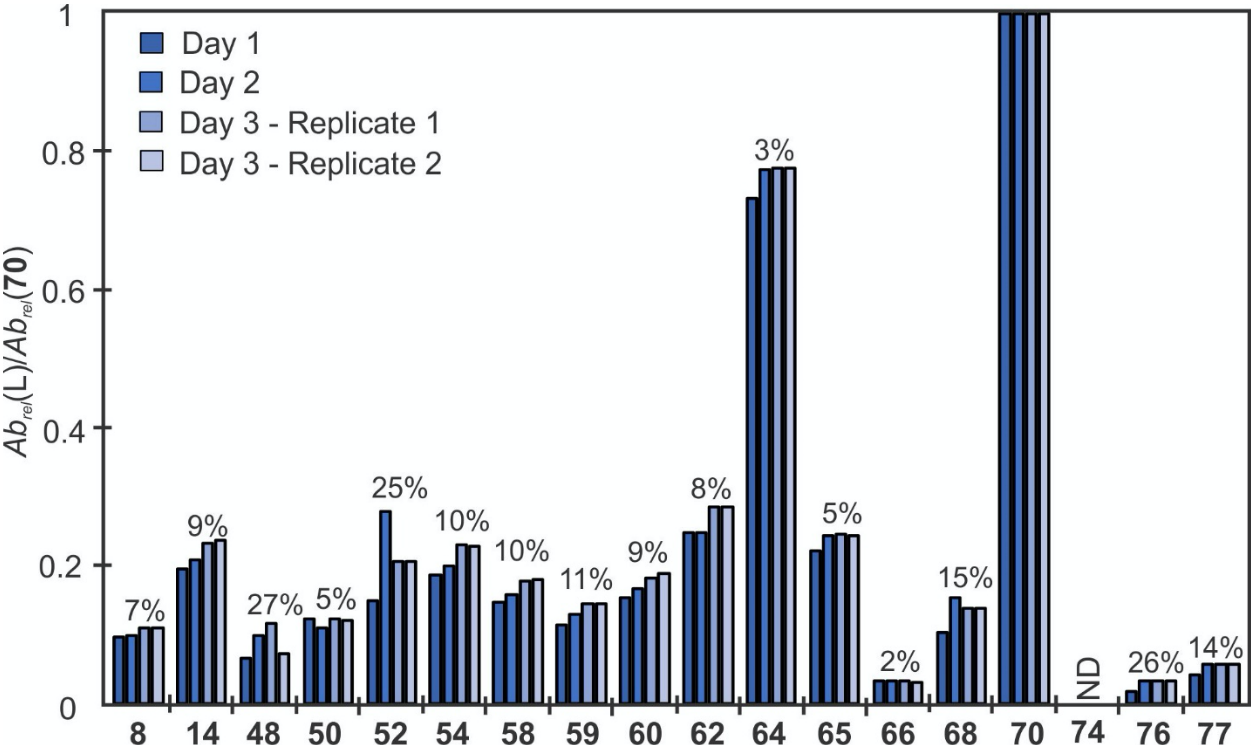


**Supplementary Figure 20.** Reproducibility of CaR-ESI-MS glycan library screening data. Normalized (to **70**) abundances of released ligands measured for aqueous ammonium acetate (100 mM, pH 6.9, 25 $^{o}$C) solutions of RBD (13 μM), SNA ($P_{ref}$, 4 μM) and Library D (50 nM of each glycan). HCD was performed using a collision energy 50 eV. The percentages shown correspond to the relative standard deviation of the normalized abundances. ND ≡ not detected.
